# Statistical design of a synthetic microbiome that suppresses diverse gut pathogens

**DOI:** 10.1101/2024.02.28.582635

**Authors:** Rita A. Oliveira, Emma McSpadden, Bipul Pandey, Kiseok Lee, Mahmoud Yousef, Robert Y. Chen, Conrad Triebold, Fidel Haro, Valeryia Aksianiuk, Riya Patel, Kaavya Shriram, Ramaswamy Ramanujam, Seppe Kuehn, Arjun S. Raman

## Abstract

Engineering functional microbiomes is challenging due to complex interactions between bacteria and their environments^1–6^. Using a set of 848 gut commensal strains and clearance of multi-drug resistant *Klebsiella pneumoniae* (*Kp*-MH258) as a target function, we engineered a functional 15-member synthetic microbiome—SynCom15—through a statistical approach agnostic to strain phenotype, mechanism of action, bacterial interactions, or composition of natural microbiomes. Our approach involved designing, building, and testing 96 metagenomically diverse consortia, learning a generative model using community strain presence/absence as input, and distilling model constraints through statistical inference. SynCom15 cleared *Kp*-MH258 across *in vitro*, *ex vivo*, and *in vivo* environments, matching the efficacy of a fecal microbiome transplant in a clinically relevant murine model of infection. The mechanism of suppression by SynCom15 was related to fatty acid production coupled with environmental acidification. SynCom15 also suppressed other pathogens—*Clostridioides difficile*, *Escherichia coli*, and other *K. pneumoniae* strains—but through different mechanisms. Sensitivity analysis revealed models trained on strain presence/absence captured the statistical structure of pathogen suppression, illustrating that community representation was key to our approach succeeding. Our framework, ‘Constraint Distillation’, could be a general and efficient strategy for building emergent complex systems, offering a path towards synthetic ecology more broadly.

## Main

Engineering communities of microbes for desired functions (‘synthetic ecology’) is of fundamental importance and holds great practical promise for addressing many problems facing humanity^7–9^. Across environmental, agricultural, and human health applications, designed communities have previously been created for specific functions. Yet despite these case-by-case successes, synthetic ecology remains largely empirical, driven by phenotypic screens, substantial prior knowledge, and context-specific information. In application to the human gut microbiome, such empiricism has typically entered design efforts in three ways. First, by mining large human cohorts or the microbiotas of laboratory mice to extract the natural statistics of existing gut communities (species prevalence and abundance, covariation structure, modules/guilds, compositional structures linked to phenotypes of interest)^10–15^. Second, by using intact or minimally processed natural communities (fecal microbial transplants, donor enrichments, fractionation/serial dilution) to transfer function (i.e. ‘top-down’ approach)^16–20^. Or third, by assembling defined consortia from individual bacterial strains but with the inclusion of rich biological priors (strain growth characteristics, functional screens of strains, pairwise interaction maps, metabolic capabilities, resource-interaction networks)^21–27^.

Why is such knowledge typically used and broadly deemed necessary for engineering consortia? A central reason is complexity. Microbiomes comprise many species whose interactions are non-linear, redundant, and context-dependent^2,10,28,29^. Higher-order effects often defeat logic learned about system function derived from single strains or pairwise interactions between strains^3,6,30^. Thus, the combinatorial complexity of *N* strains is not additive, but scales as 2*^N^* thereby becoming prohibitively expansive with respect to design efforts: a hypothetical consortium comprised of 100 strains encodes 10^30^ possible strain combinations. In physics and chemistry, engineering principles are scaffolded by ‘forward models’—generative theory that maps parts to function in a deterministic fashion^31^. However in biology, such forward models are absent and therefore functional properties cannot be inferred from components alone. Collective biological behaviors—so-called ‘emergent’ properties—arise from interaction patterns that are difficult to observe, let alone encode *a priori*. Thus, lacking a physics of emergence and confronted with daunting complexity, current bacterial consortia design strategies default to embedding statistical, ecological, or mechanistic priors. Inclusion of these priors leads to two key limitations with respect to consortia design: (i) biasing outcomes towards what nature already makes feasible rather than exploring a truly synthetic space and (ii) resulting in bespoke solutions that exist on a case-by-case basis rather than transferable design principles that can be broadly employed. Both limitations place a ceiling on the potential of synthetic ecology with respect to future application towards desired target functions.

There has been a recent substantial rise in the capacity to culture bacteria and create strain banks^11,16,21,27,32^. This trend along with the observations regarding consortia design noted above motivate a natural question of both fundamental and practical importance: are microbiomes programmable? By ‘programmable’, we specifically mean two nested capabilities. First, the capacity to create a reliable map relating microbiome composition to function learned (e.g. predictive capacity) without the need to specify context-dependent mechanistic or functional priors^33^. But also second, going beyond prediction, the ability to use that map to build microbiomes that satisfy a desired target function with performance that holds across environments (i.e. generative capacity and functional robustness)^10,34^. Taking inspiration from engineering, in circuit theory existing principles enable the construction of apparently complex, functional, and robust circuits from a list of parts. Analogously, can design principles be derived that enable construction of synthetic, functionally robust microbiomes from strains alone?

When forward models are unavailable or limiting, statistical physics has addressed design by solving the so-called ‘inverse problem’: inferring constraints that reflect statistical patterns within observational data alone, then using only these constraints to synthesize new systems^31,35,36^. Classic examples include the approach of maximum-entropy to learn energy models from correlation structures in data; more recent examples include using sophisticated statistical networks (e.g. artificial intelligence-based models) to represent relationships and generate complex systems like proteins and materials^37–40^. These examples suggest a promising strategy forward with respect to consortia design. Rather than encode any priors involving strain phenotype, strain interactions, or mechanism of desired target function, solving the inverse problem would involve inferring a statistical model of latent constraints residing within an ensemble of synthetic consortia that is predictive of the target function and is generative. If successful, this strategy would enable the construction of truly synthetic microbiomes—consortia that do not resemble natural microbiomes—through establishing design principles that are statistical in nature (e.g. ‘statistical design’) and thereby reframe consortia design as a data-driven engineering problem.

In this work, we engineered a complex 15-member synthetic microbiome—SynCom15—that suppressed a multi-drug resistant (MDR) pathogenic strain of *Klebsiella pneumoniae* (*Kp*-MH258) across *in vitro*, *ex vivo*, and *in vivo* conditions, matching the rapidity, efficacy, and robustness of a whole fecal microbial transplant (FMT) in a specific pathogen-free (SPF) mouse model of infection. Rather than relying on biological knowledge involving strain phenotype, strain interactions, natural microbiome composition, or mechanism of suppression to accomplish this task, we engineered SynCom15 by first solving the inverse problem through statistically learning a generative model of consortia design *de novo* using an ensemble of just 96 synthetic consortia spanning various sizes that we evaluated for pathogen suppression, then distilling taxonomic constraints learned by the model. Critically, the only biological information used in our process was strain genome sequence for characterizing genomic diversity amongst our strain collection. Through a series of experiments, we found that SynCom15 (i) suppressed *Kp*-MH258 through fatty acid production coupled with environmental acidification, (ii) suppressed other pathogenic strains of *K. pneumoniae*, *Escherichia coli*, and *Clostridioides difficile*, (iii) promoted recovery of antibiotic-induced dysbiosis *in vivo*, and (iv) was not identifiable from sequencing statistics of healthy human gut microbiomes. We pose that our statistical approach—‘Constraint Distillation’— may be generally useful for engineering bacterial consortia towards executing desired target functions.

## Results

### Suppression of K. pneumoniae by engineered, diverse microbiomes does not translate across environments

To design a synthetic bacterial consortium that suppresses MDR *K. pneumoniae*, we first created a strain bank of 848 gut commensals from a set of 28 healthy human volunteers and whole-genome sequenced all strains (Methods). A subset of these strains have been described and characterized previously^32,41^. These commensals collectively reflected a breadth of phylogenetic diversity typically found in human gut microbiomes (**Extended Data Fig. 1**, **Supplementary Table 1A**). Without incorporating more information, the total possible design space encompassed by this strain bank spanning all individual strains, pairs of strains, and higher-order combinations was ∼10^250^—a space that is impractical from the standpoint of design. Moreover, constructing consortia that constitute greater than 50 strains is markedly challenging from the perspective of experimental culture. We therefore sought to reduce the number of possible strains we could use to design consortia.

We performed a UMAP-based (Uniform Manifold Approximation and Projection) dimension-reduction on an alignment of gene content across all strains (Methods). This defined a latent embedding of strain relatedness that clustered strains by their phylum-level phylogenetic designation, thereby providing a coarse-grained view of genomic diversity across our strain bank (**Extended Data Fig. 2**, **Supplementary Table 1B**). We next sought to select a subset of strains from our original 848 strains that comprised a manageable set from the perspective of engineering synthetic consortia in the laboratory yet retained the structure of phylogenetic diversity captured by the UMAP embedding. We therefore selected 46 strains that retained the phylogenetic diversity of our strain bank—a number of strains that was within the range of previously published designed consortia (**Fig. 1A**, **Supplementary Table 2**) (Methods)^16,27^.

**Figure 1.**
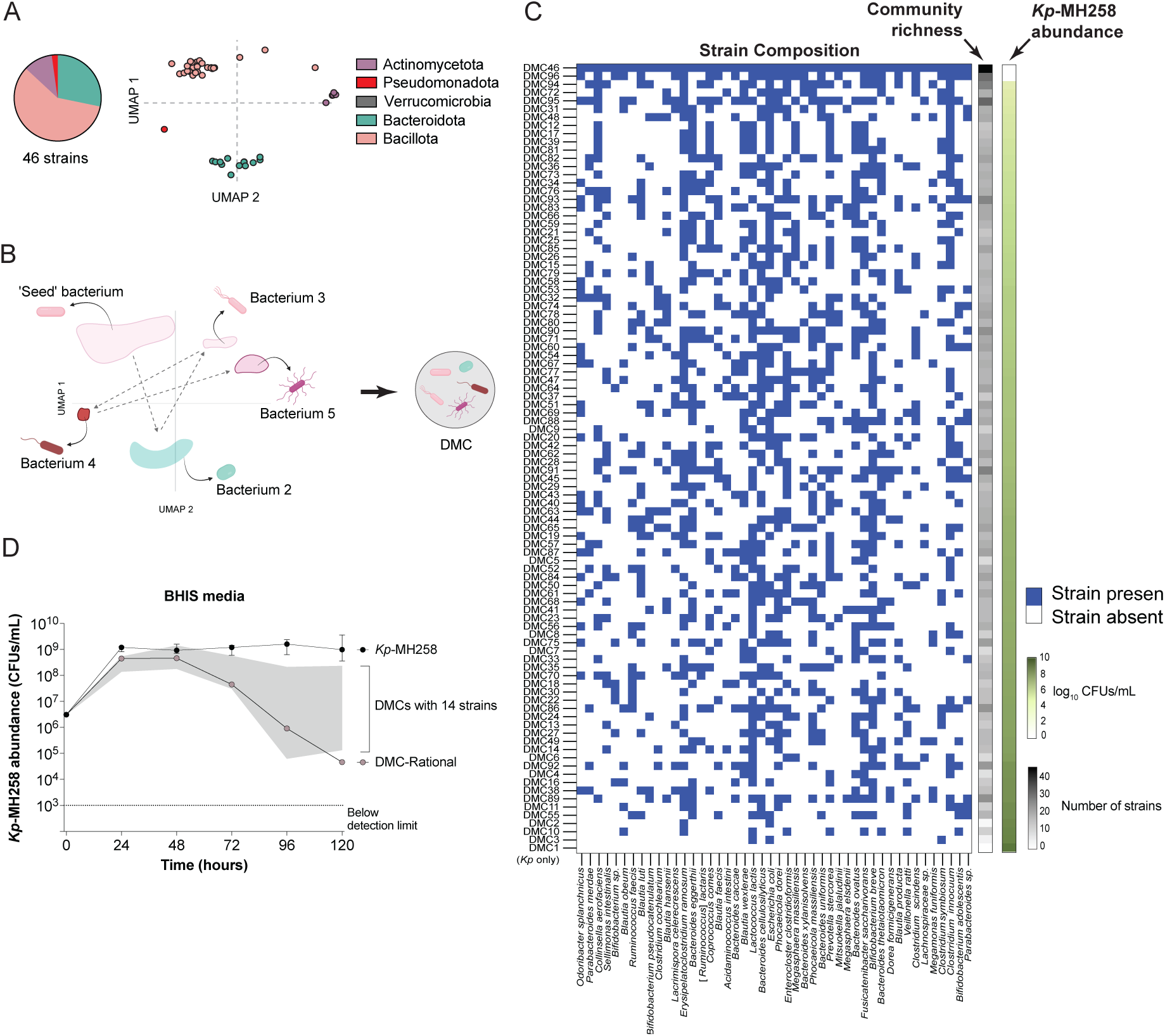
Designing, building, and testing synthetic microbiomes for suppression of *Kp*-MH258. **(A)** Subset of commensal strain bank used to engineer synthetic consortia; strains are dots (n = 46) within Uniform Manifold Approximation Plot (UMAP) axes. **(B)** Schematic of algorithm used to create Designed Microbial Communities (DMCs). **(C)** Matrix of 96 engineered DMCs (rows) as well as *Kp*-MH258 in monoculture (bottom row) by strains (columns). Blue pixel indicates strain was included in the DMC, white pixel indicates strain was not included in the DMC. Each row is labeled by ‘Community richness’—the number of strains included in the design of the DMC—and ‘*Kp*-MH258 abundance’—the colony forming units (CFUs) per milliliter of *Kp*-MH258 remaining after 120 hours of co-culture with the DMC in Brain Heart Infusion media supplemented with cysteine (BHIS media). **(D)** Abundance of *Kp*-MH258 (y-axis) across time (x-axis) in monoculture (black dots), in co-culture with all DMCs from panel C comprised of 14 strains (grey distribution), or in co-culture with DMC-Rational (14 strains) in BHIS media.

Despite down-sampling our commensal strain bank from 848 strains to 46 strains, the resulting combinatorial complexity (∼35.2 billion possibilities) was still unmanageable. Therefore, we sought to impose a constraint on the consortia we designed—construct metagenomically diverse consortia. The rationale behind this choice was to capture a large possible functional capacity of each designed consortium. Previous studies have constructed diverse consortia by simply combining phylogenetically diverse strains^11,27^. However, no algorithmic process currently exists for creating ensembles of diverse consortia spanning various membership sizes from strain banks. Primarily this is because the hierarchical nature of phylogeny makes defining differences between strains in a quantitative manner challenging^32^. We therefore developed an algorithm that constructs consortia directly off the two-dimensional UMAP coordinates shown in **Fig. 1A** (**Supplemental Discussion**) (Methods). Briefly, for a consortium consisting of *N* members, the algorithm proceeds as follows. First, a random seed bacterium is chosen; next, a second bacterium is chosen such that it is distant from the seed in the UMAP coordinate space, a third bacterium is chosen such that it is distant from the first two bacteria, and so on until *N* members are chosen (**Extended Data Fig. 3A**). This strategy of design contains no biological information outside of strain genomes for engineering microbial consortia nor any information regarding strain phenotype or pathogen suppression *a priori*. We employed our design strategy and engineered 96 ‘Designed Microbial Communities’ (DMCs) (**Fig. 1B,C**). The 96 DMCs spanned a size of 2 to all 46 strains (**Supplementary Table 3A**). We found that the communities comprising greater than 10 strains asymptotically reached a degree of taxonomic diversity represented by the community containing all 46 strains (‘DMC46’) (**Extended Data Fig. 3B**). Thus, our algorithm succeeded in creating an ensemble of metagenomically diverse consortia.

We then tested all DMCs for their capacity to suppress an MDR strain of *K. pneumoniae*—*Kp-*MH258—after 120 hours of co-culture in Brain Heart Infusion Media supplemented with cysteine (BHIS) (**Extended Data Fig. 4A**) (Methods). The MH258 strain was isolated from a patient at Memorial Sloan Kettering Hospital and was representative of the epidemic multilocus sequence type (ST) 258 clone harboring the *bla*_KPC_-encoded carbapenemase. We chose this strain to use as our target for suppression because it is amongst the most multi-drug resistant strains that have been previously characterized, exhibiting resistance against a diversity of antibiotics^42^. BHIS media was chosen because all 46 gut commensal strains robustly grew in this environment (**Extended Data Fig. 4B**, **Supplementary Table 3B**). We found that the set of 96 DMCs exhibited a continuous range of suppressive capacities in a reproducibly robust fashion (**Fig. 1C**, **Extended Data Fig. 4C,D**, **Supplementary Table 3C,D**). During the first 48 hours of co-culture, the abundance of *Kp-*MH258 remained uniformly high, after which community effects began to manifest—some consortia suppressed *Kp-*MH258 greater than four orders of magnitude while others had minimal impact on its fitness (**Extended Data Fig. 4E**, **Supplementary Table 3D**).

We evaluated three different compositional qualities of our DMCs that could be associated with suppression of *Kp-*MH258. First, previous studies have suggested the importance of including up to ten strains that are phylogenetically diverse in consortia designed to suppress *K. pneumoniae*^27^. We found that the suppressive capacity of DMCs was not associated with the number of species within our diverse consortia (**Extended Data Fig. 5A**). As one example, DMC17 contained 12 strains and suppressed *Kp-*MH258 four orders of magnitude while DMC92 contained 19 strains and suppressed *Kp-*MH258 less than one order of magnitude (**Extended Data Fig. 5B**, left). As another example, there were a total of 16 DMCs that contained 14 strains, but their observed capacity to suppress *Kp*-MH258 varied from one to four orders of magnitude (**Extended Data Fig. 5B**, right). Second, previous studies have suggested the importance of including *Escherichia coli* strains in the background of large diverse consortia for suppressing *K. pneumoniae* due to hypothesized overlap in metabolic niche^27^. Our set of 46 strains contained one *E. coli* strain. We found no relationship between the membership size of consortia that contained our *E. coli* strain and the capacity of a consortium to suppress *Kp-*MH258 (**Extended Data Fig. 5C**). As an example, DMC92 contained an *E. coli* strain plus 18 other strains and DMC03 contained an *E. coli* strain and two other strains. Despite the difference in consortia size, both DMCs exhibited similar suppressive capacity of less than one order of magnitude (**Extended Data Fig. 5C,D**). Finally, we tested whether the presence of any single strain was uniformly responsible for the suppressive capacity of DMCs. Our results showed that the presence of 14 individual strains was significantly associated with the suppression of *Kp-*MH258 (**Extended Data Fig. 6**). However, we also found that the presence of any one of these strains on its own did not guarantee suppression by the DMC. DMCs containing these strains showed a range of suppression that overlapped extensively with DMCs not containing these strains (**Extended Data Fig. 7**). Collectively, these results illustrated that previously described taxonomic heuristics—diversity/richness of consortium, presence/absence *of E.* coli—and presence/absence of single strains were insufficient to describe our results.

Recent work has suggested that simplistic statistical models that do not consider higher-order interactions between strains can be successful at predicting community phenotype from the presence or absence of individual strains. A study that fit simple regression models across a large number of previously published studies found that additive models alone that do not consider even pairwise interactions between species can predict community phenotype^33^. Inspired by this work, we next combined all 14 strains whose individual presence was significantly associated with *Kp-*MH258 suppression (see **Extended Data Fig. 6**, yellow box). This consortium was not within the 96 DMCs we had engineered in **Fig. 1C** and was designed using only the results from our functional screen. We therefore termed this consortium ‘DMC-Rational’. Notably, DMC-Rational included strains displaying the most extreme regression coefficients derived from a linear regression model that successfully related *Kp*-MH258 abundance with the additive effect of individual strains alone (r^2^ of ∼0.8) (**Supplementary Table 3E**) (Methods). We found that DMC-Rational suppressed *Kp-*MH258 greater than four orders of magnitude and was the best performing DMC of all the DMCs comprised of 14 strains (**Fig. 1D**). This result demonstrated that taking a simplistic statistical approach—constructing a consortium based on the suppressive effect of each strain considered individually without interaction effects—enabled designing a consortium that suppressed *Kp-*MH258 in BHIS.

Previous attempts at translating the suppression of *K. pneumoniae* by engineered bacterial consortia from *in vitro* to *in vivo* settings have demonstrated, at best, limited success. As an example, recently published work engineering a large diverse consortium of gut symbionts showed robust *K. pneumoniae* suppression *in vitro* but only mild suppression *in vivo* with suppression lasting only one day after infection^27^. Moreover, tests of pathogen suppression by engineered consortium *in vivo* have primarily involved germ-free mouse models of infection—a model system unrelated to clinically relevant courses of infection by *K. pneumoniae* where treatment by broad-spectrum antibiotics creates an open niche for pathogen colonization while patients retain a background microbiome and are subject to colonization by bacteria in the environment^11,16,27^.

Given these observations, we next tested a select group of DMCs for their capacity to suppress *Kp*-MH258 in a more clinically realistic *in vivo* model of infection. Using a specific pathogen-free (SPF) mouse model of *Kp-*MH258 infection, we administered several DMCs spanning a range of suppressive capacities observed in BHIS media. Singly-housed mice were treated with broad-spectrum antibiotics (metronidazole, neomycin, vancomycin) for four days, colonized with *Kp-*MH258 on the day after stopping antibiotics, and subsequently gavaged two days later with either phosphate buffered saline (PBS), a whole-stool fecal microbiota transplant from mice not given antibiotic treatment (FMT), or a specific DMC (**Fig. 2A**) (Methods). We chose to evaluate four DMCs—DMC46 (comprised of all 46 strains), DMC96 (comprised of 28 strains), DMC94 (comprised of 24 strains), and DMC-Rational (comprised of 14 strains)—because they suppressed *Kp-*MH258 in BHIS media at least four orders of magnitude. We also chose to evaluate DMC86 (comprised of 20 strains) because this DMC was amongst the worst performing with respect to pathogen suppression in BHIS media—suppressing *Kp*-MH258 within one order of magnitude—while containing a similar number of strains as DMC94.

**Figure 2.**
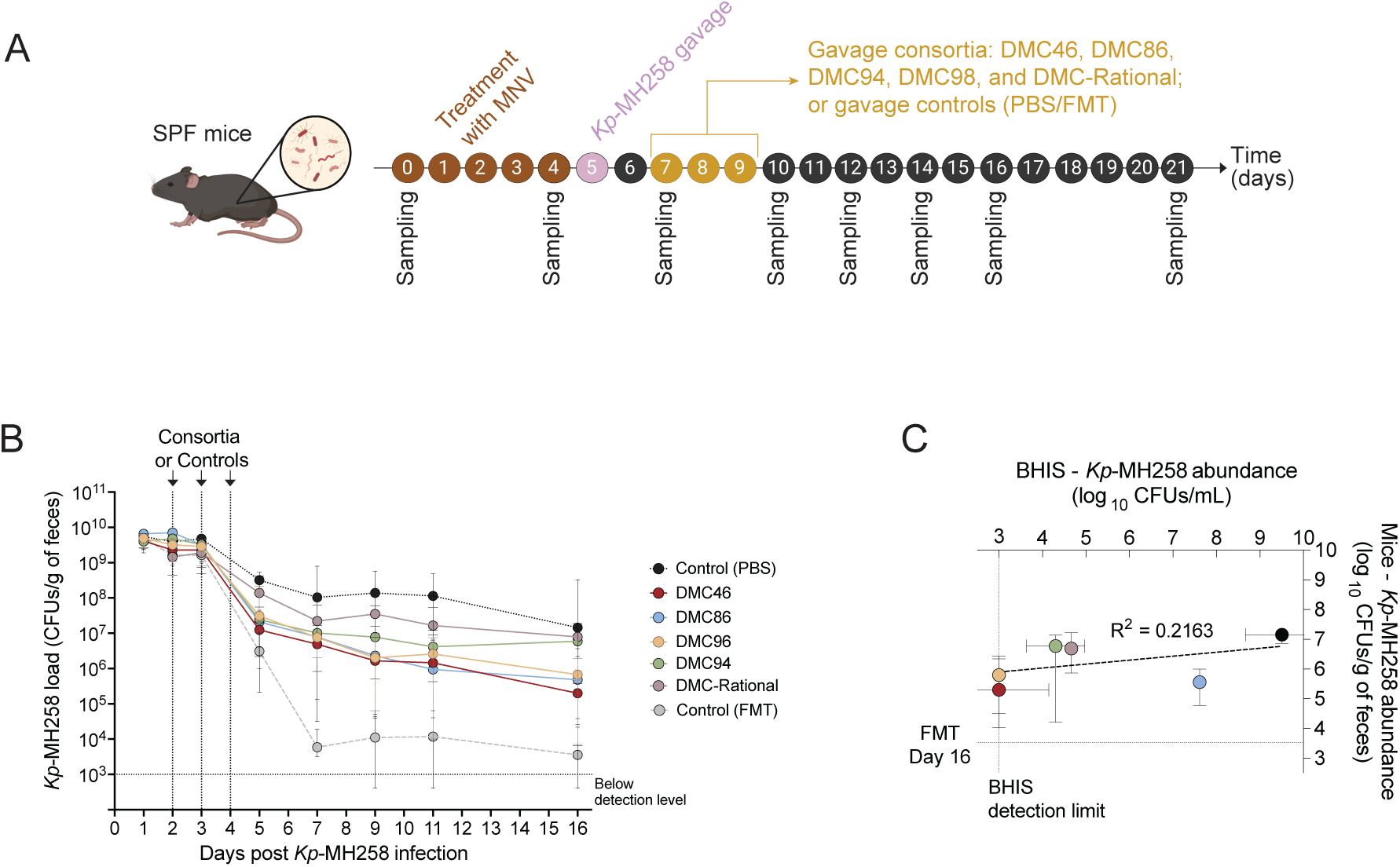
Suppression of *Kp*-MH258 by DMCs *in vitro* does not translate *in vivo*. **(A)** Experimental workflow for *in vivo* model of *Kp*-MH258 infection. Specific-pathogen-free (SPF) mice were treated with metronidazole, neomycin, and vancomycin (‘MNV’) in drinking water for four days, gavaged with *Kp*-MH258 (‘*Kp*-MH258 gavage’), then gavaged with either phosphate buffered saline (PBS), fecal microbial transplant (FMT), or one of several DMCs three days in a row. **(B)** Dynamics of fecal *Kp*-MH258 abundance through time for mice gavaged with either PBS, FMT, or one of five DMCs (see color key). Lines represent median values and error bars represent interquartile range. (C) Suppressive capacity of DMCs (colored dots; color scheme follows panel B) observed in BHIS at 120 hours of co-culture (x-axis) versus SPF-infected mice at day 16 post *Kp*-MH258 infection (y-axis). PBS gavage in mice and monoculture of *Kp*-MH258 alone in BHIS is represented by the black dot. Error bars represent interquartile range.

First, our results showed that no consortium suppressed *Kp-*MH258 to the same extent as an FMT. By three days after gavage of the FMT, fecal abundance of *Kp*-MH258 dropped six orders of magnitude to 10^4^ colony forming units (CFUs) per gram of feces (**Fig. 2B**, **Supplementary Table 4A**). In contrast, the best performing consortium in BHIS media—DMC46—was three orders of magnitude worse than the FMT intervention three days after the last gavage. Second, we found that seven days after the last gavage of the intervention (Day 11 of the experiment), the suppressive effect of all consortia were within 1.5 orders of magnitude of each other, yielding a fecal abundance of *Kp-*MH258 between 10^6^ and 10^8^ CFUs/gram of feces. This result illustrated that all tested consortia performed in a roughly similar manner to each other within our *in vivo* model (**Fig. 2B**, **Supplementary Table 4A**). Third, despite suppressing *Kp-*MH258 several orders of magnitude in BHIS, DMC-Rational most closely resembled the effect of saline intervention on *Kp-*MH258 colonization in mice. Thus DMC-Rational lost its suppressive potential when evaluated in our *in vivo* setting. A systematic analysis of our results showed that the suppressive capacity of a DMC in BHIS media was not associated with the capacity to suppress *Kp-*MH258 in our *in vivo* model (**Fig. 2C**, **Supplementary Table 4B**).

Overall, our results showed that using simple heuristics—size/diversity of consortium, presence of *E. coli*, or the cumulative effect of statistically important individual strains—was insufficient for engineering a microbiome that suppressed *Kp*-MH258 across different environments. A parsimonious explanation for these results is that environmental idiosyncrasies demand bespoke consortia designs, making generally effective consortia inherently difficult to engineer. However, prior work suggests that translatability can indeed be achieved by identifying modules of interacting taxa. As a notable example, statistical analysis of human developing microbiomes led to the identification of a module of interacting species that, when combined into a consortium, recapitulated key taxonomic dynamics in a porcine model of microbiome development^10^. More broadly, the idea of modularity has been proposed as a construct for enhancing robustness across changing contexts in network science, protein biophysics, and gene-regulatory circuits^34,43–48^. We therefore posed an alternative hypothesis: the capacity to suppress *Kp*-MH258 across different environments may reside in groups of strains whose joint effects form a collective unit such that translatable suppression is a property of the interaction architecture rather than any single taxon or simple rule. Because such groups are not directly observable from composition alone and because there does not exist obvious priors for encoding translatability of function into consortia design, we turned to statistical learning as a path forward.

### Discovering a 15-member bacterial consortium through machine learning and distillation of model constraints

Machine-learning (ML) used towards designing synthetic bacterial consortia has typically incorporated a number of detailed strain qualities including (i) individual growth rates, (ii) growth rates in co-culture, and (iii) metabolic capacities that were either experimentally determined or inferred from gene content^22–24,33^. On the other hand, using patterns of strain presence/absence with regression-based statistics including interactions have accurately predicted ecosystem structure-function relationships but is limited in detecting greater than pairwise interactions between strains in the limit of finite sampling that may be critical for going beyond prediction and towards generative design^33^. As ML models are capable of modeling non-linear interactions, we reasoned that using ML but considering only the pattern of strain presence/absence across our DMCs could aid in creating a statistical generative model of consortia design for suppressing *Kp-*MH258 without requiring the need to consider biological priors.

We trained two models—a Random Forests (RF) ML model and a deep learning-based neural network (DNN)—using the pattern of strain presence/absence across our 96 DMCs as input and the CFUs of *Kp-*MH258 after 120 hours of co-culture with a DMC as output (Methods). Notably, neither model included any information regarding (i) survival of microbes in culture, (ii) dynamics of bacterial strains, (iii) phenotypic qualities of strains, (iv) nature of interactions between microbes, or (v) interaction with *Kp*-MH258. Instead, the input to the model was a 46-dimensional vector defined by the 46 strains that could be used to design a DMC. Each DMC was characterized by ‘1’s and ‘0’s where a ‘1’ reflected that a specific strain was included in the DMC and a ‘0’ reflected that a specific strain was left out of the DMC. We observed an in-sample validation r^2^ value of 0.98 for the RF model and 0.74 for the DNN model, indicating well-trained models (**Supplementary Table 5**).

We next evaluated the capacity of the RF and DNN models to predict 60 new DMCs that were not included in the original 96 DMCs we constructed. These 60 new DMCs spanned a range of membership sizes and predicted capacities for *Kp-*MH258 suppression (**Supplementary Table 6A**). We found that both the RF and DNN models accurately predicted the suppressive capacity of the 60 new DMCs with the RF model mildly outperforming the DNN model (r^2^ of 0.6 versus 0.52) where variability in model performance was predictably large at low values of *Kp*-MH258 abundance (**Fig. 3A**, top; **Supplementary Table 6B**). Thus, considering only the pattern of included strain presence/absence for a given DMC was sufficient to yield generative models of consortia engineered to suppress *Kp*-MH258 in BHIS media.

**Figure 3.**
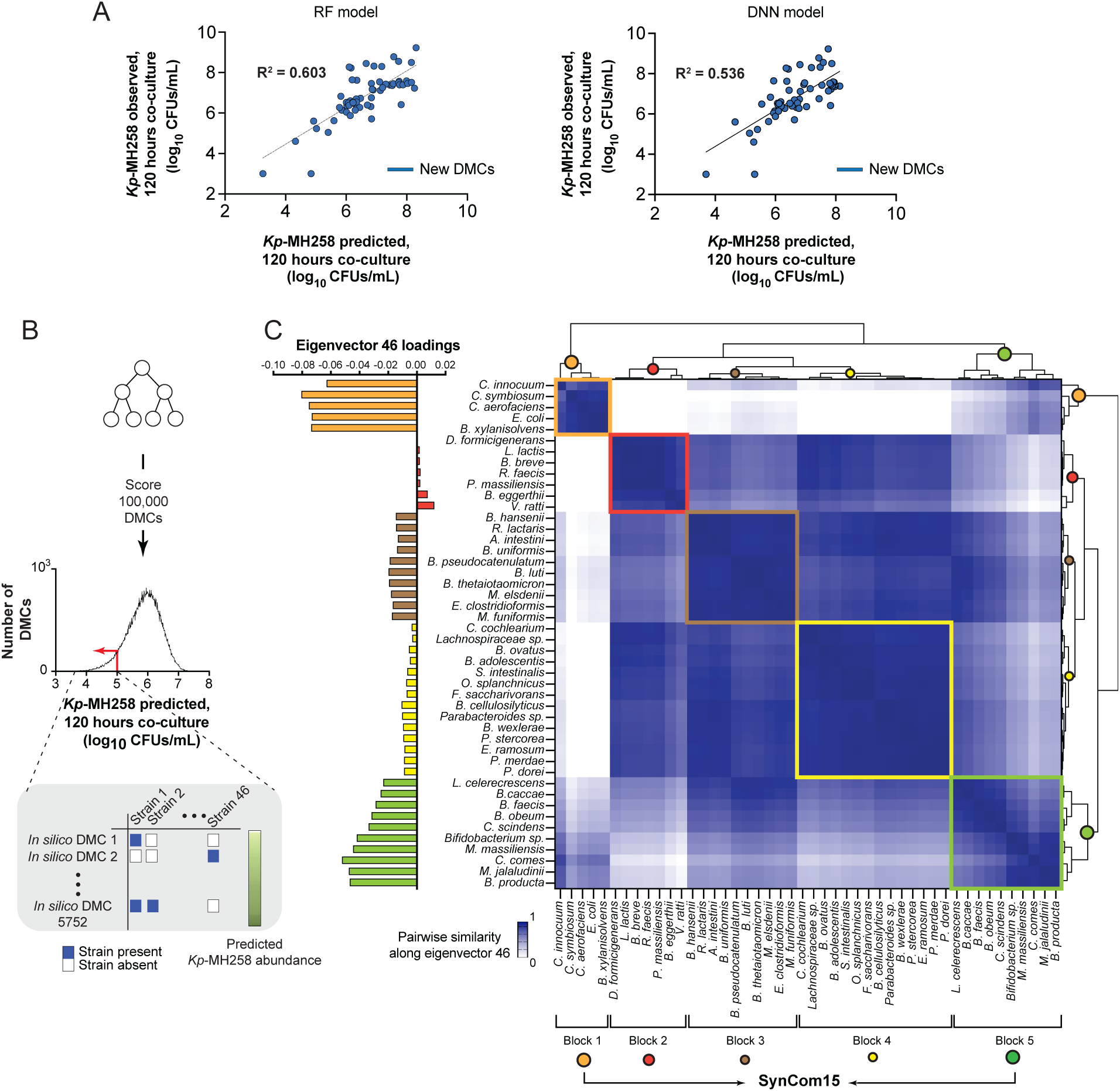
Distillation of constraints learned by generative model yields SynCom15. **(A)** Predicted abundance of *Kp*-MH258 at 120 hours of co-culture with DMCs (x-axis) from a Random-Forests model (RF, left) and a deep-learning based neural network (DNN, right) trained on data from **Fig. 1C** versus observed abundance (y-axis). Each dot is one of 60 possible out-of-sample DMCs not represented in the 96 DMCs shown in **Fig. 1C**. **(B)** 100,000 DMCs were generated *in silico* and scored by the trained RF model. Histogram of 100,000 DMCs is shown as a function of predicted *Kp*-MH258 abundance for each DMC after 120 hours of co-culture. DMCs for which the RF model predicts less than 10^5^ CFUs/mL of *Kp*-MH258 remaining after 120 hours of co-culture are selected and converted into a matrix (inset). Rows are each of 5,752 possible DMCs (‘*In silico* DMC’); columns are one of 46 strains used to create DMCs in **Fig. 1C**; entries reflect whether a strain is present (blue pixel) or absent (white pixel) in the DMC. **(C)** Loadings of each of the 46 strains onto eigenvector 46 of the matrix defined in panel B (left). Similarity between all pairs of strains along eigenvector 46 was computed and resulting matrix was hierarchically clustered resulting in distinct blocks of strains (right). Block 1 (orange) plus Block 5 (green) is defined as SynCom15.

We therefore next sought to use these models to design a suppressive consortium. However, we noted this would be challenging: our ML models exhibited substantial variation in the prediction of *Kp*-MH258 abundance below 10^5^ CFUs because the input data was naturally limited within this regime (**Fig. 3A**). Moreover, studying the models themselves was not a viable solution either. While ML models like RF and DNN-based algorithms can be useful for prediction, their internal logic is opaque, making it difficult to extract actionable design principles or understand compositionally why particular consortia were effective. With RF models, feature importance scores are useful for discriminatory tasks but do not capture combinatorial, redundant, and context-dependent relationships necessary for revealing statistical interactions underlying the success of model prediction and therefore necessary for generation of a system. In the case of DNNs, no mapping exists between the architecture of the model and the input dataset since DNNs are inherently non-linear, opaque transformations.

Given the above considerations, one option would have been to collect data on more consortia such that the suppressive regime would be better sampled. However, it was not clear how this could be done. First, simply creating more consortia would likely not have led to sampling a greater number of suppressive communities. Second, the high variance of the ML predictions in consortia performance below *Kp*-MH258 values of 10^5^ would likely necessitate testing lots more consortia. Given these issues, we desired a way to turn noisy ML predictions into actionable guidance without conducting a large new experimental screen. We therefore posed an alternative hypothesis: a large ensemble of outputs from our ML models could be leveraged for extracting new taxonomic information about *Kp*-MH258 suppression. Our rationale for this hypothesis was as follows. If predictions are unreliable in the extreme low-abundance regime but are globally reasonable, then what can be trusted is the relative prediction of the model across many consortia. Thus, we could use the ML models as rankers—assigning relative rankings such that certain consortia are placed consistently lower than others—rather than relying on the precise numerical prediction for a specific consortium. Relatedly, we recognized that any single ‘top-ranked’ consortium could be inaccurately predicted. However, aggregation across many independently top-ranked consortia could potentially reveal taxonomic regularities that are shared. Patterns that recur across many such consortia would be far more likely to reflect true design signals than model idiosyncrasies. This approach—(i) broadly query a complex and opaque model, (ii) keep only the consistent answers, and (iii) summarize those answers into a compact rule set—is analogous to the logic of ‘distillation’ in the field of artificial intelligence (AI)—a framework used to make ‘black-box’ AI models understandable^49–51^. We therefore adopted the ethos of distillation by generating a large *in silico* ensemble of consortia, selecting the consortia that were ranked as strong-suppressors, and performing statistical inference via spectral analysis to extract compositional constraints (which strains co-occur in association with *Kp*-MH258 suppression, which strains are allowable substitutions, which strain combinations should be avoided) (see **Supplementary Discussion** for further details). We chose to use our RF model for this approach as it performed better than the DNN model.

We generated 10^5^ DMCs by randomly sampling strain combinations from the 46-strain pool where the totality of DMCs spanned a range of predicted *Kp*-MH258 suppression values as determined by the RF model. We chose 10^5^ communities to sample greater than 10^3^ of the complexity of the design space as defined by each strain independently (46 total strains corresponds to a space of 4.6 × 10^4^ considering 10^3^ oversampling). Each DMC was represented as a binary vector of strain presence/absence and scored using the RF model, resulting in a matrix of 10^5^ DMCs (rows) by 46 strains (columns) where each DMC was annotated with a predicted abundance of *Kp*-MH258. We then filtered this design space by DMCs predicted to suppress *Kp*-MH258 to a value below 10^5^ CFUs, yielding a subset of 5,752 DMCs that were predicted to be high-performing (**Fig. 3B**, **Supplementary Table 7A**). Next, to uncover model constraints on consortia composition, we performed Singular Value Decomposition (SVD) of this set of DMCs defined by their patterns of strain presence/absence. This resulted in a spectrum of eigenvectors that defined axes of variation across DMCs predicted to function particularly well. We associated the pattern of loadings of DMCs onto each eigenvector with the predicted capacity to suppress *Kp*-MH258 for all 5,752 DMCs (Methods) (**Supplementary Table 7B**,**C**). This analysis revealed that eigenvector 46—encoding <1% of data variance—defined taxonomic variation amongst *in silico* generated DMCs that was most strongly associated with *Kp-*MH258 suppression (**Extended Data Fig. 8**). This axis therefore represented a functionally relevant statistical constraint latent within our RF model.

Two groups of strains contributed similarly onto eigenvector 46: one group of five strains (‘Block 1’) comprising *Clostridium innocuum*, *Clostridium symbiosum*, *Colinsella aerofaciens*, *E. coli*, and *Bacteroides xylanisolvens*, and another group of ten strains (‘Block 5’) comprising *Lacrimispora celerecrescens, Bacteroides caccae*, *Blautia faecis*, *Blautia obeum*, *Clostridium scindens*, a *Bifidobacterium* species, *Megasphaera massiliensis*, *Coprococcus comes*, *Mitsuokella jalaludinii*, and *Blautia producta* (**Fig. 3C**) (Methods). Three other blocks of strains contributed relatively less towards eigenvector 46 (‘Block 2’, ‘Block 3’, and ‘Block 4’) (**Fig. 3C**). We further observed that the strains belonging to Blocks 1 and 5 co-contributed to eigenvector 46 with each other more than with strains defining Blocks 2, 3, and 4 (**Fig. 3C**, **Supplementary Table 7D-F**). Thus, the 15 strains collectively comprising Blocks 1 and 5 formed the statistically-inferred putative core within DMCs predicted to suppress *Kp*-MH258—the basis of a designed consortium that we termed SynCom15 (**Fig. 3C**).

### SynCom15 suppresses Kp-MH258 across diverse contexts

We evaluated the capacity for SynCom15, as well as Blocks 1 through 5, to suppress *Kp*-MH258 when co-cultured in BHIS media as well as two *ex-vivo* contexts—media created from the cecal contents of germ-free mice (‘GF cecal content media’) and media created from the cecal contents of SPF mice that were treated with antibiotics for four days then given normal drinking water for three days (‘Ab-SPF cecal content media’)—a scenario mimicking our *in vivo* infection model (**Fig. 4A**) (Methods). The two *ex-vivo* conditions contained information regarding the host, host diet, and host microbiome, and thus the collection of *in vitro* and *ex-vivo* contexts reflected vastly different environments for evaluating the suppressive capacity of a consortium on *Kp*-MH258. We found that of all the consortia tested, SynCom15 was the most suppressive community across all three environments (**Fig. 4B**, **Supplementary Table 7G**).

**Figure 4.**
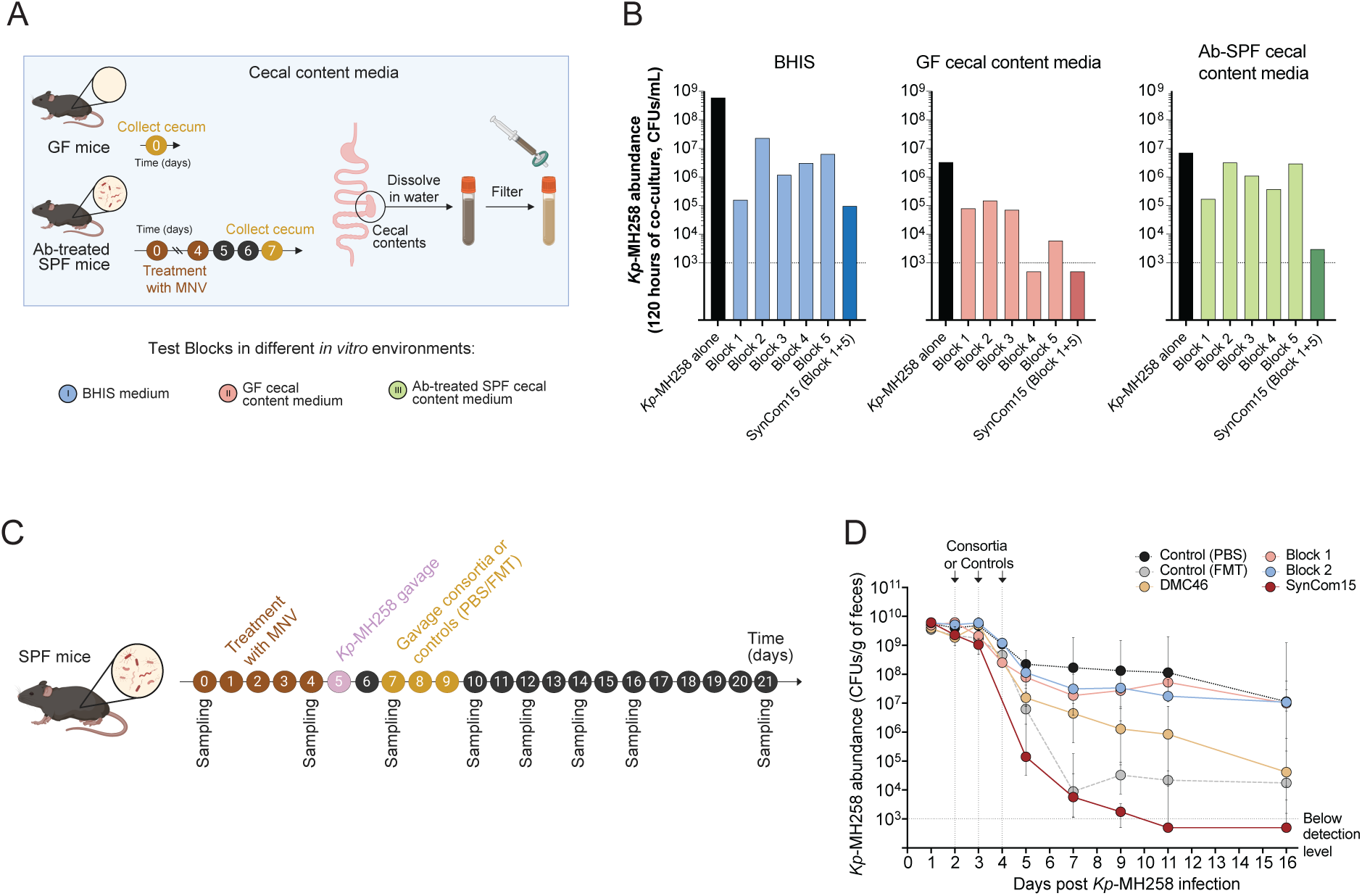
**SynCom15 suppresses MDR *Kp-*MH258 across diverse environments. (A)** Workflow for creating cecal content media from germ-free (GF) or antibiotic-treated specific pathogen free (Ab-treated SPF) mice (box). **(B)** Abundance of *Kp*-MH258 at 120 hours of co-culture (y-axis) with each Block defined in Fig. 3C, SynCom15, and *Kp*-MH258 in monoculture (x-axis) across BHIS media and medias defined in panel A. **(C)** Workflow for assaying suppressive capacity of Blocks 1, 2, DMC46, SynCom15, and controls (PBS and FMT) in *in vivo* model of *Kp-*MH258 infection (similar scheme as Fig. 1D). **(D)** *Kp*-MH258 fecal abundance versus time for mice gavaged with various interventions (color key). Lines represent median values; error bars represent interquartile range.

We next tested the efficacy of SynCom15 in our SPF mouse model of infection (**Fig 4C**). We found SynCom15 suppressed *Kp*-MH258 five orders of magnitude on the first day after the third gavage, six orders of magnitude the third day after the third gavage, seven orders of magnitude the fifth day after the third gavage and maintained a level of *Kp*-MH258 below detection the seventh day after the third gavage (**Fig. 4D**). Notably, *Kp*-MH258 suppression due to SynCom15 was more rapid and a greater magnitude than that of an FMT. We compared the suppressive effect of SynCom15 with that of saline and three other consortia: Block 1, Block 2, and DMC46. We evaluated Block 1 because it effectively suppressed *Kp*-MH258 in BHIS media and was identified as consistently comprising a group of strains associated with a suppressive phenotype towards DMCs by three independent statistical descriptions—(i) the effect of strains considered individually (see **Extended Data Fig. 6**, yellow box), (ii) the feature importance scores resulting from the RF model, and (iii) the results of our distillation analysis (**Extended Data Fig. 9**, **Supplementary Table 7H**). We evaluated Block 2 as a plausible negative control as it failed to suppress *Kp*-MH258 in BHIS, GF cecal content media, and Ab-SPF cecal content media, and this group of strains were associated with poor suppressive capacity from our distillation analysis. We re-evaluated DMC46 because it contained all SynCom15 strains, and it was the consortia that exhibited the highest suppressive capacity of all DMCs tested from **Fig. 1C** in our *in vivo* infection model. We found that Blocks 1 and 2 did not suppress *Kp*-MH258 greater than saline intervention (**Fig. 4D**, **Supplementary Table 7I**). Additionally, we found that DMC46 suppressed *Kp*-MH258 several orders of magnitude less on days 5 through 11 after infection by *Kp*-MH258 and was more gradual than SynCom15, illustrating that removing 31 strains from DMC46 was beneficial for achieving a heightened suppressive capacity of *Kp-*MH258 *in vivo* (**Fig. 4D**, **Supplementary Table 7I**). We note that the increased suppressive capacity of DMC46 observed between days 11 and 16 compared to the results shown in **Fig. 2B** was also observed with the PBS intervention. Thus, pathogen suppression during this time period was likely due to the natural course of microbiota repair as our murine model was not germ-free but was open to the environment.

Collectively these results illustrated that (i) SynCom15 suppressed *Kp*-MH258 across diverse, unrelated conditions and (ii) SynCom15 intervention resulted in a marked, rapid, and sustained clearance of *Kp*-MH258 in a pre-clinically relevant *in vivo* mouse model of infection. These findings motivated us to understand the mechanism of action of SynCom15 and to evaluate its effect on gut pathogens other than *Kp*-MH258.

### SynCom15 ameliorates antibiotic-induced dysbiosis, suppresses Kp-MH258 by acidification and metabolic remodeling, and suppresses other gut pathogens through various mechanisms

To understand how SynCom15 suppresses *Kp*-MH258, we started by characterizing the suppressive capacity of SynCom15 from a taxonomic perspective. First, we evaluated the suppressive capacity of each single strain on *Kp*-MH258 in BHIS media. We found that only four strains suppressed *Kp*-MH258 more than two orders of magnitude in BHIS while the remaining 11 strains suppressed *Kp*-MH258 less than one order of magnitude (**Extended Data Fig. 10A**). This result illustrated that interactions between strains were key for specifying the suppressive capacity of SynCom15, consistent with prior literature demonstrating that the capacity to decolonize *K. pneumoniae* in *in vitro* culture was not a property of single strains^27^. Additionally, the Block 1 consortium comprised all four of the strains that suppressed *Kp*-MH258 more than two orders of magnitude yet failed to suppress *Kp*-MH257 *in vivo*, suggesting that the translatable capacity of SynCom15 could not be explained simply by the effects of individual strains *in vitro*. Second, as SynCom15 includes a strain of *E. coli*—a species whose inclusion in diverse consortia was previously shown to be important for designing consortia that suppress *K. pneumoniae*—we evaluated the effect of removing *E. coli* from SynCom15 on pathogen suppression **(Extended Data Fig. 10B**)^27^. We found that SynCom15 without *E. coli* matched the suppressive capacity of SynCom15 in both BHIS and GF cecal content media (**Extended Data Fig 10C**,**D**). This result illustrated that the effect of SynCom15 was not merely due to the inclusion of *E. coli* in the consortium. Third, we evaluated whether the suppressive capacity of SynCom15 could be inferred via taxonomic information gleaned from our mouse experiments. Taxonomic profiling of fecal samples collected through the course of our experiment revealed that 10 of the 15 strains in SynCom15 engrafted in at least one of the mice within the cohort with one such strain being *E. coli*. Additionally, dynamics of SynCom15 strains showed that 5 of the 10 strains were present at detectable fractional abundances throughout the course of the experiment—*C. symbiosum*, *B. xylanisolvens*, *C. innocuum*, *B. obeum*, and *B. caccae* (**Extended Data Fig. 10B**, inset)

(Methods). We therefore built three communities—(i) a community constituting strains that consistently engrafted the mice (10 species), (ii) a community constituting strains that consistently engrafted the mice but without *E. coli*, and (iii) a community constituting strains that were detected in mice across all timepoints (5 species). The 10-member community and the 10-member community without *E. coli* suppressed *Kp*-MH258 two orders of magnitude in BHIS and did not exhibit suppression in GF cecal content media. The 5-member community suppressed *Kp*-MH258 one order of magnitude in BHIS and did not exhibit suppression in GF cecal content media (**Extended Data Fig. 10C,D**). Together these results suggested that elucidating a mechanism of action of SynCom15 from the perspective of taxonomic composition alone would be challenging. We therefore turned to a more function-centric view for understanding how SynCom15 suppresses *Kp*-MH258.

We leveraged the results of our mouse experiments and analyzed the longitudinal dynamics of fecal microbiota structure to evaluate the effect of administering SynCom15 after antibiotic exposure and infection by *Kp*-MH258 (Methods). We found that the effect of administering SynCom15 mirrored the FMT intervention, returning the state of the microbiota to that observed prior to antibiotic treatment (**Fig. 5A**, **Extended Data Fig. 11A,B**, **Supplementary Table 8A,B**). Moreover, we noted that while treatment with FMT resulted in the return of mouse-specific gut bacteria (e.g. *Duncaniella* and *Paramuribaculum*), treatment with SynCom15 resulted in the detectable presence of human-specific gut bacteria (e.g. genera belonging to the genera *Bacteroides* and a bloom of *Bifidobacterium*) (**Extended Data Fig. 11C**, **Supplementary Table 8C**). Histology of the mouse colon showed that SynCom15 was well tolerated as an intervention showing no evidence of inflammation or tissue insult (**Extended Data Fig. 11D**). Together, these results demonstrated that SynCom15 restores community diversity of the gut microbiota as measured by coarse phylogenetic composition but results in a compositional signature distinct from natural murine microbiota. Our findings motivated a deeper investigation into the functional mechanisms by which SynCom15 suppresses pathogens.

**Figure 5.**
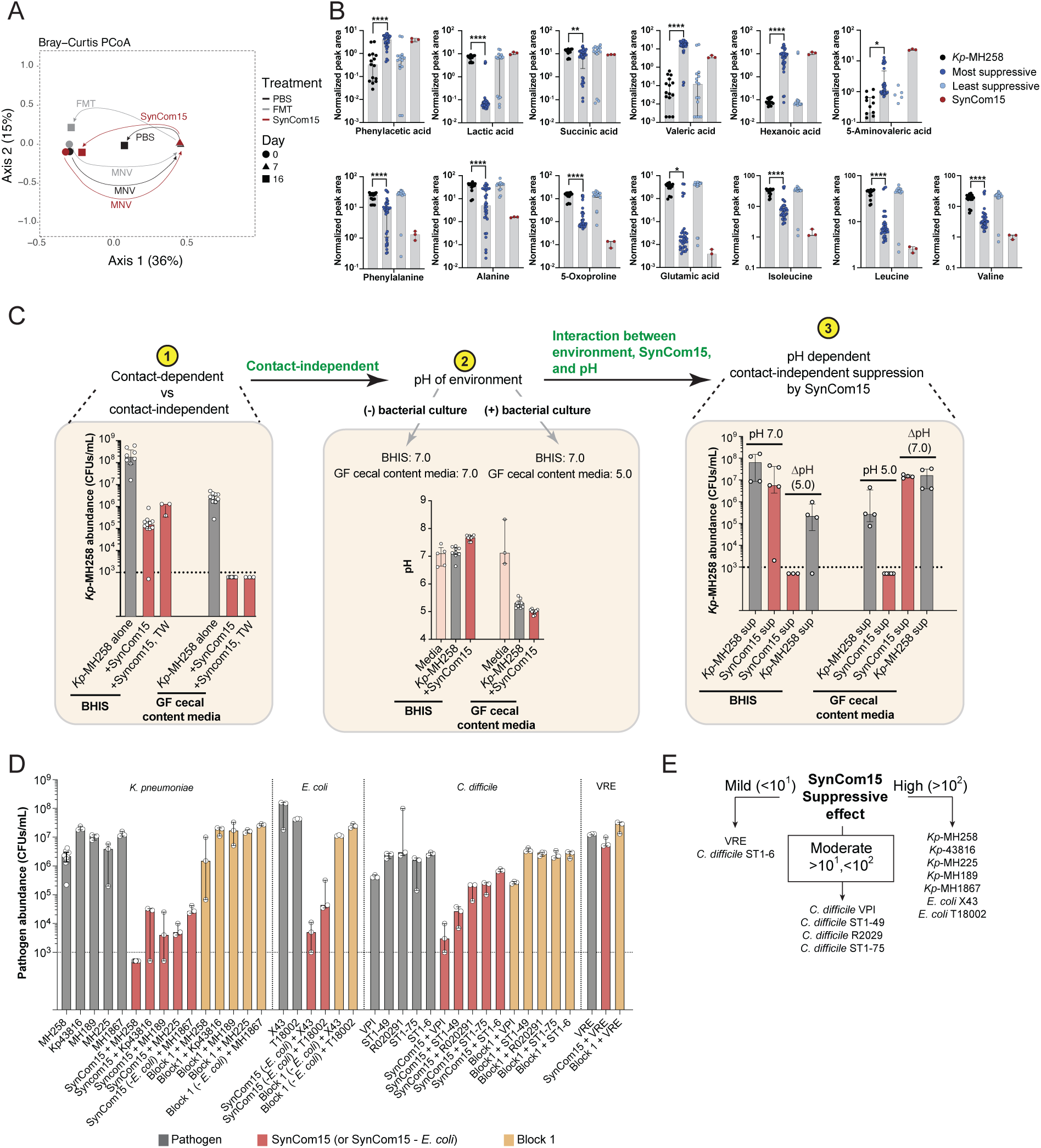
SynCom15 rescues antibiotic-induced dysbiosis, suppresses *Kp*-MH258 through metabolic remodeling plus environmental acidification, and suppresses a diversity of other gut pathogens. (A) PCoA of fecal microbiota for SPF mice on day 0, 7, and 16 of experiment shown in Fig. 4C; colored shape is centroid for indicated cohort. (B) Bar plots show distribution of normalized peak areas (y-axis) for each metabolite feature distinguishing the most suppressive from least suppressive DMCs (individual panels) across three different culture settings at the 72-, 96-, and 120-hour culture timepoint: *Kp*-MH258 in monoculture (black), *Kp*-MH258 in co-culture with the most suppressive DMCs (dark blue), *Kp*-MH258 in co-culture with the least suppressive DMCs (light blue), and *Kp*-MH258 in co-culture with SynCom15 (maroon). *p < 0.05; **p < 0.01; ****p < 0.0001. (C) (Left) Abundance of *Kp*-MH258 in BHIS and GF cecal content media in monoculture, co-culture with SynCom15, or co-culture with SynCom15 in a transwell culture setup (SynCom15 TW) after 120 hours. (Middle) pH of BHIS remains constant at 7.0 with and without bacterial culture; pH of GF cecal content media drops from 7.0 to 5.0 with bacterial culture. (Right) Abundance of *Kp*-MH258 in BHIS and GF cecal content media when co-cultured for 120 hours with supernatants collected from *Kp*-MH258 cultured alone (*Kp*-MH258 sup) or SynCom15/*Kp*-MH258 co-culture (SynCom15 sup). Both supernatants in BHIS media exhibited a natural pH of 7.0; supernatants were then acidified to a pH of 5.0 (ΔpH). Both supernatants in GF cecal content media exhibited a natural pH of 5.0; pH of supernatants was then increased to 7.0 (ΔpH). (D) Abundance of various pathogens (y-axis) in monoculture (grey), co-culture with SynCom15 (red), co-culture with SynCom15 without *E. coli* strain (‘*-E. coli*’) (red), or co-culture with Block 1 (defined in Fig. 3C). Experiments were performed in GF cecal extract media. In panels C and D, bars represent median values and error bars represent interquartile range. (E) Summary of suppressive effects of SynCom15 on abundance of all tested pathogens.

To begin interrogating the functional output of SynCom15, we performed targeted metabolic profiling on supernatants from the five most suppressive and least suppressive DMCs, as well as SynCom15, when co-cultured with *Kp*-MH258 in BHIS media (Methods). This metabolic panel encompassed metabolites central to microbiome functional capacities—fatty acids, indoles, amines, and amino acids—and has been used previously to describe human gut microbiomes (Methods)^32,41,52,53^. Our analysis revealed two distinguishing signatures of suppressive consortia—select amino acid depletion and fatty acid production (**Fig. 5B**, **Extended Data Fig. 12A**, **Supplementary Table 9A,B**). Relative to *Kp-*MH258 monoculture, the most suppressive DMCs and SynCom15 exhibited elevated levels of phenylacetic acid, valeric acid, hexanoic acid, and 5-aminovaleric acid while SynCom15 uniquely showed additional increases in lactic and succinic acid (**Fig. 5B**, **Supplementary Table 9A,C**). Additionally, multiple amino acids including those with aromatic and non-polar sidechains—phenylalanine, alanine, isoleucine, leucine, and valine—plus glutamate and 5-oxoproline were significantly depleted in the suppressive DMCs as well as SynCom15 (**Fig. 5B**, **Supplementary Table 9A**,**C**). Thus, elevated levels of medium-chain fatty acids (valeric acid, hexanoic acid) with aromatic and amino-acid derived acids (phenylacetic acid and 5-aminovaleric acid) suggested environmental acidogenesis consistent with the previously described mechanism involving pH-dependent suppression of gram-negative *Enterobacteriaceae*^54^. Additionally, depletion of branched-chain (isoleucine and leucine), aromatic (phenylalanine), and nitrogen handling (glutamate, 5-oxoproline, alanine) amino acids suggested nutrient competition that could potentially limit *K. pneumoniae* growth, in accord with previous studies illustrating the importance of branched-chain, aromatic, and glutamate family amino acids (glutamate and proline) for *K. pneumoniae* fitness^55–57^. Consistent with these results, metabolic profiling of fecal samples collected from our mouse model of *Kp*-MH258 infection showed a significant increase in fatty acids at day 10 of the experiment and depletion of amino acids at day 12 of the experiment when mice were given the SynCom15 intervention compared to intervention with saline (**Extended Data Fig. 12B**). Together, our findings suggested that metabolic remodeling of the environment by SynCom15 was important for suppression of *Kp*-MH258.

Following our metabolomic analysis, we performed a series of experiments to evaluate the mechanistic basis of suppression by SynCom15 in BHIS and GF cecal content media. First, we evaluated whether the metabolic environment created by SynCom15, as opposed to contact-dependent direct physical interaction, was the driver of *Kp*-MH258 suppression. To test this idea, we co-cultured SynCom15 with *Kp*-MH258 in a transwell system where SynCom15 was physically separated from *Kp*-MH258 but could still interact with *Kp*-MH258 in a contact-independent manner via metabolites (Methods). We found that in both BHIS media and GF cecal content media, SynCom15 suppressed *Kp*-MH258 to nearly the same degree in the transwell system as when in co-culture (**Fig. 5C**, left; **Supplementary Table 10A**). This result illustrated that the mechanism of *Kp*-MH258 suppression by SynCom15 was through contact-independent interaction with the pathogen.

Previous literature has demonstrated that acidification of the environment can substantially augment the effect of fatty acid production on suppression of gram negative *Enterobacteriaceae*^54^. Because suppression in the GF cecal content media condition was substantially higher than that of BHIS, we evaluated the pH of each media condition with and without bacterial culture. Both media conditions exhibited a baseline pH of 7.0. However, SynCom15 co-cultured with *Kp*-MH258 in GF cecal content media resulted in a drop in pH to 5.0 whereas the pH of the co-culture in BHIS media remained at 7.0. We next evaluated whether a pH drop by itself was sufficient to suppress *Kp*-MH258. We grew *Kp*-MH258 alone in BHIS and GF cecal content media and found that the pH of the culture in BHIS was maintained at 7.0 while the pH of the culture in GF cecal content media dropped to 5.0. However, despite the acidification of the culture in GF cecal content media, the fitness of *Kp*-MH258 remained high (**Fig. 5C**, left and middle; **Supplementary Table 10B**). This result illustrated that environmental acidification alone was not sufficient to explain suppression of *Kp*-MH258.

Our results so far showed that both SynCom15/*Kp*-MH258 co-culture and *Kp*-MH258 monoculture drop environmental pH, but that SynCom15/*Kp*-MH258 co-culture also produces fatty acids. We therefore hypothesized that SynCom15 suppresses *Kp-*MH258 through a combination of fatty acid production and acidification of the media—a mechanism consistent with previous studies interrogating gut microbiome-mediated decolonization of gram negative *Enterobacteriaceae*^54^. To test this hypothesis, we first collected the supernatant from SynCom15 co-cultured for 120 hours with *Kp*-MH258 in BHIS and GF cecal content media. Second, we acidified the supernatant derived from BHIS media to a pH of 5.0 and buffered the supernatant derived from GF cecal content media to a pH of 7.0 (Methods). Finally, we inoculated the acidified and buffered supernatants with *Kp*-MH258 for 120 hours and compared the resulting pathogen abundance in supernatants that were not acidified and buffered. We found that acidification of the supernatant in BHIS suppressed *Kp*-MH258 greater than five orders of magnitude; buffering of the supernatant in GF cecal content media increased the abundance of *Kp*-MH258 by greater than four orders of magnitude (**Fig. 5C**, right; **Supplementary Table 10C**). To evaluate whether this effect was specific to SynCom15, we collected the supernatant of *Kp*-MH258 monocultured for 120 hours in BHIS and in GF cecal content media. We then followed the same protocol as above. We found that *Kp*-MH258 cultured in its own supernatant in BHIS and GF cecal content media maintained at their native pH retained its fitness, decreasing less than an order of magnitude in each condition (**Fig. 5C**, right; **Supplementary Table 10C**). We also found that changing the pH in either condition affected the fitness of *Kp*-MH258 but not to the same degree as the supernatant collected from SynCom15/*Kp*-MH258 co-culture (**Fig. 5C**, right; **Supplementary Table 10C**). We next evaluated whether the suppressive effect of the supernatant collected from the co-culture of SynCom15/*Kp*-MH258 was because of the inclusion of *Kp*-MH258. We collected the supernatant of SynCom15 cultured on its own for 120 hours in both BHIS and GF cecal content media, followed by the same protocol of acidification and buffering with inoculation of *Kp*-MH258. We found that the presence of *Kp*-MH258 in the co-culture with SynCom15 did not affect the suppressive phenotype of the supernatant nor the effect of pH on the suppressive phenotype (**Extended Data Fig. 12C**, **Supplementary Table 10D**). Collectively, these results suggest that environmental acidification with fatty acid production was a plausible mechanism by which SynCom15 suppresses *Kp*-MH258 across different environments.

Several previous studies have illustrated the connection between fatty acid production, nutrient depletion, and environmental acidification in naturally occurring microbiomes with suppression of various pathogens^54,58–61^. We therefore sought to evaluate the suppressive effect of SynCom15 against a library of phylogenetically distinct gut pathogens. These included different strains of MDR *K. pneumoniae*, *E. coli*, and *C. difficile* as well as a single strain of vancomycin resistant *Enterococcus* (VRE). We chose these pathogens because of (i) their clinical prevalence with respect to disease, (ii) the difficulty associated with their treatment, and (iii) because they cause disease through different biological mechanisms—*C. difficile* colitis through toxin production and *K. pneumoniae*, VRE, and *E. coli* infection through induction of colitis and/or gut barrier translocation leading to sepsis. We evaluated the co-culture of SynCom15 with these pathogens in GF cecal content media. We note that due to antibiotic selection overlap, we evaluated the fitness of *Kp-*MH1867 and MDR *E.coli* strains when co-cultured with SynCom15 without its *E. coli* strain. With respect to strains of *Enterobacteriaceae*, we found that co-culture with SynCom15 suppressed all strains of *K. pneumoniae* and *E. coli* three to four orders of magnitude (**Fig. 5D**, **Supplementary Table 11A**). We found that SynCom15 suppressed two *C. difficile* strains—VPI and ST1-49—by two orders of magnitude, two other strains—R020291 and ST1-75—greater than one order of magnitude; and one strain—ST1-6—less than one order of magnitude (**Fig. 5D**, **Supplementary Table 11A**). Finally, we found there to be minimal effect of SynCom15 on suppressing VRE (**Fig. 5D**, **Supplementary Table 11A**). Across all pathogens, we found that the Block 1 consortium was generally not suppressive, resulting in the same level of pathogens and the pathogen cultured alone (**Fig. 5D**, **Supplementary Table 11A**). Thus, SynCom15 was able to suppress several strains of phylogenetically diverse gut pathogens.

We next evaluated mechanisms by which SynCom15 suppressed a selection of *K. pneumoniae*, *E. coli*, and *C. difficile* strains. We co-cultured SynCom15 with *Kp*-MH225, *E. coli* X43, and *C. difficile* VPI and ST1-49 in GF cecal content media using a transwell culture system for 120 hours. We found that separating SynCom15 from *Kp*-MH225 abrogated the suppression of the pathogen: SynCom15 suppressed *Kp*-MH225 three orders of magnitude in co-culture but less than one order of magnitude in the transwell culture system (**Extended Data Fig. 13**). In contrast, the transwell co-cultures of SynCom15 with *E. coli* X43 and *C. difficile* VPI and ST1-49 strains recapitulated pathogen suppression observed in co-culture. Using the same protocol of buffering the GF cecal content media condition used in **Fig. 5C**, we found that suppression of *E. coli* X43 and *C. difficile* VPI and ST1-49 were pH dependent (**Extended Data Fig. 13**, **Supplementary Table 11B**).

Collectively, our interrogation of pathogen suppression by SynCom15 revealed several findings. First, environmental acidification coupled with fatty acid production appeared to be a mechanism by which SynCom15 suppressed *Kp*-MH258, possibly augmented by depletion of specific amino acids. Second, though designed to suppress *Kp*-MH258, SynCom15 was able to suppress gut pathogens belonging to diverse phylogenies that manifest pathology in humans by different biological mechanisms. Third, the capacity to suppress these pathogens was dependent on different mechanisms—through indirect, pH-dependent suppression for *E. coli* X43, *C. difficile* VPI, and *C. difficile* ST1-49 strains versus through direct physical interaction for *Kp*-MH225—illustrating the degeneracy of mechanisms for pathogen suppression encoded by SynCom15. Lastly, beyond only pathogen suppression, we found that SynCom15 repairs the microbiota after antibiotic-induced dysbiosis, promoting its return to a healthy and diverse state.

### Strains of SynCom15 are not prevalent or abundant in gut microbiomes of healthy humans

We explored the extent to which SynCom15 was represented across healthy humans who provided stool samples from which we created our strain bank. We first interrogated the prevalence of the genera constituting SynCom15 strains in these fecal samples. We found that SynCom15 was composed of a diversity of genera observed across the set of healthy gut microbiomes (**Fig. 6A**). Next, we interrogated the prevalence of the SynCom15 species across the fecal samples of the healthy donors (Methods). We found that no healthy human microbiome contained more than eleven of the SynCom15 species above a fractional abundance of 0.1% (**Fig. 6B**; **Supplementary Table 12**). Moreover, we found certain SynCom15 species to be sparse in their prevalence across donors as detected by metagenomic sequencing. *M. jalaludinii* was not detectable in any donor; *M. massiliensis* was detectable in two donors; *C. symbiosum* in three donors; and *C. scindens* in four donors. Amongst strains that were most prevalent, *B. obeum* and *B. faecis* were detectable in 20 donors; *L. celerecrescens* in 14 donors; *C. comes* in 13 donors; *B. caccae* in 12 donors. Finally, we interrogated the fractional abundance of SynCom15 species across the fecal samples. We found SynCom15 species were present at a relative abundance of less than 5% across all donors, with most species being found at a relative abundance of less than 0.5% (**Fig. 6C**).

**Figure 6.**
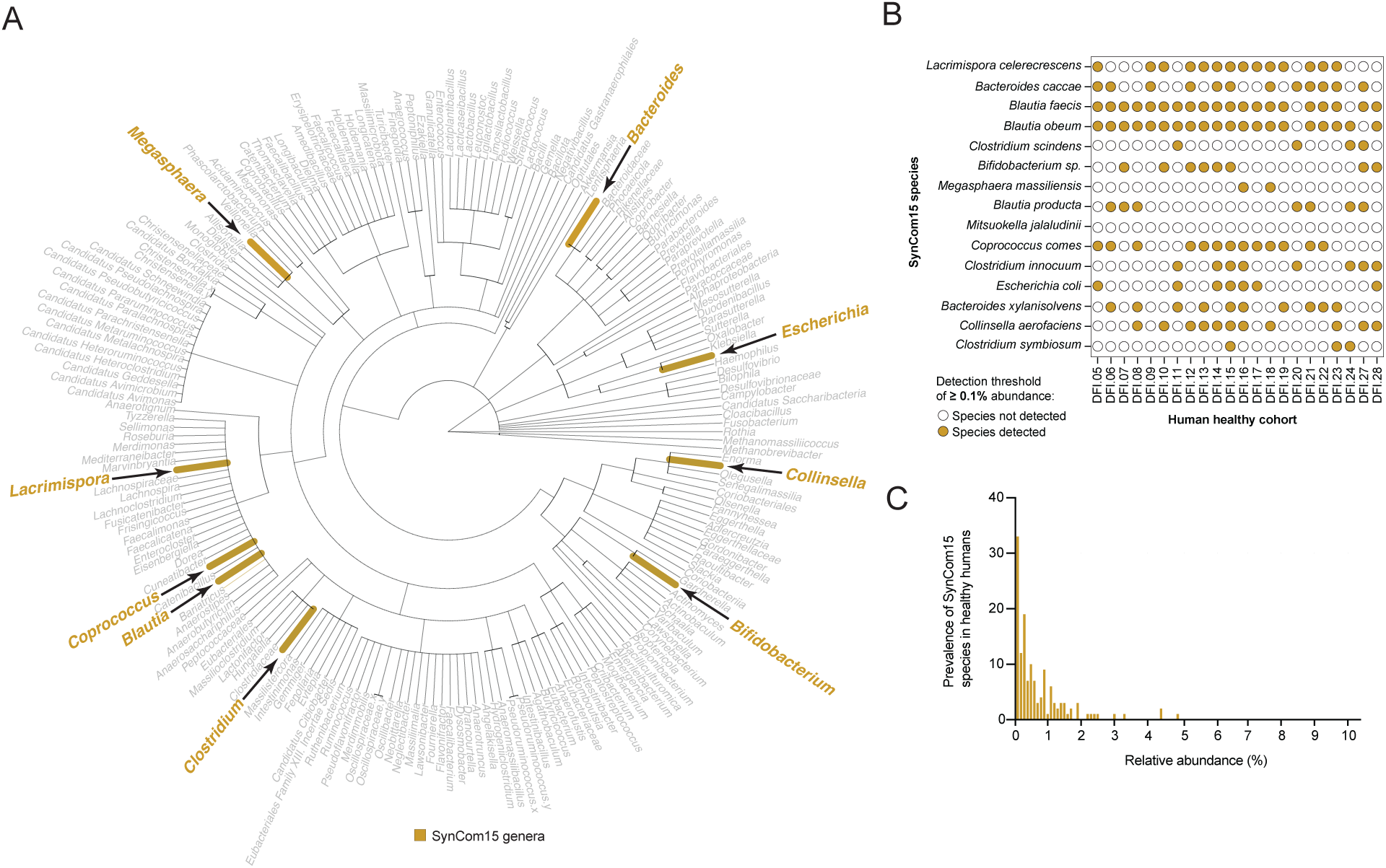
Comparison of SynCom15 composition with composition of healthy human microbiomes. (A) Phylogenetic tree of genera present across fecal microbiomes of human donors. Brown genera are those found across SynCom15 strains. (B) Prevalence pattern for species of SynCom15 (rows) across donor fecal microbiomes (DFI is Duchossois Family Institute; columns). (C) Histogram of relative abundance for SynCom15 species (x-axis) across all fecal samples from population of healthy human donors.

Together, these findings illustrated two conclusions. First, the composition of SynCom15 was distinct from that found across healthy human gut microbiotas determined by metagenomic sequencing. This was either because SynCom15, as a community, did not exist in the healthy samples from our cohort or because several of SynCom15 strains were undetectable by our sequencing methods due to their low abundance. Second, the strains comprising SynCom15 were low prevalence and abundance amongst fecal samples of healthy donors. This result highlighted the power of generating and using broadly diverse strain banks for engineering synthetic bacterial communities as compared to strain banks reflecting the compositional abundance and prevalence distributions gleaned from analysis of natural human microbiomes—an approach previously used to design synthetic microbiomes^10–12^.

### Describing communities by strain presence/absence was important for learning a generative model of pathogen suppression

Our overall approach to deriving SynCom15 followed six sequential steps: (1) <u>C</u>ompress the design space and (2) <u>D</u>esign consortia implementing an algorithmic constraint based on selection for genomic diversity, (3) <u>B</u>uild and (4) <u>T</u>est consortia for function, (5) <u>L</u>earn a statistical model of function from patterns of strain presence/absence of designed consortia, and (6) Infer constraints learned by the model. As this approach is distinct from the typical ‘Design-Build-Test-Learn’ (DBTL) approach used in systems theory engineering, we term this approach ‘Constraint Distillation’. We note that for the specific function of *Kp*-MH258 suppression, Constraint Distillation was not iterative: SynCom15 was directly derived from just one round of Compress/Design-Build/Test-Learn-Infer. Thus, we next explored two facets of Constraint Distillation: (i) its experimental efficiency and (ii) key choices we made that enabled our framework to converge on SynCom15.

To quantify the experimental efficiency of Constraint Distillation in generating SynCom15, we computed an effective compression score that considers three measurements: (i) the total complexity of information prior to implementing each step of Constraint Distillation, (ii) the total complexity of information remaining after implementing each step of Constraint Distillation, and (iii) the information needed to converge on the output of each step of Constraint Distillation (**Fig. 7A**) (**Supplementary Discussion**). Starting with a strain bank of 848 gut commensal strains and reducing the possible space of strains to 46 required whole genome sequencing of all strains—a total of nearly 10^12^ base pairs of information—and subsequently implementing a UMAP-based dimension-reduction which required no additional information. This step reduced the space of design from 848 to 46 strains resulting in a ∼10^230^ compression of information yet left greater than 35 billion strain combinations as the space of design. To arrive at SynCom15 required performing 179 experiments where each experiment evaluated a single DMC for its capacity to suppress *Kp*-MH258 in BHIS media (96 experiments to train and validate the predictive model; 23 experiments to evaluate reproducibility of assay; 60 experiments to demonstrate the generative capacity of the predictive model). This yielded a compressive power of ∼10^11^ (**Fig. 7B**). These findings illustrated that Constraint Distillation enabled navigation of a remarkably high-dimensional combinatorial space to ultimately arrive at SynCom15 using only the genome sequences of the individual strains as prior biological knowledge.

**Figure 7.**
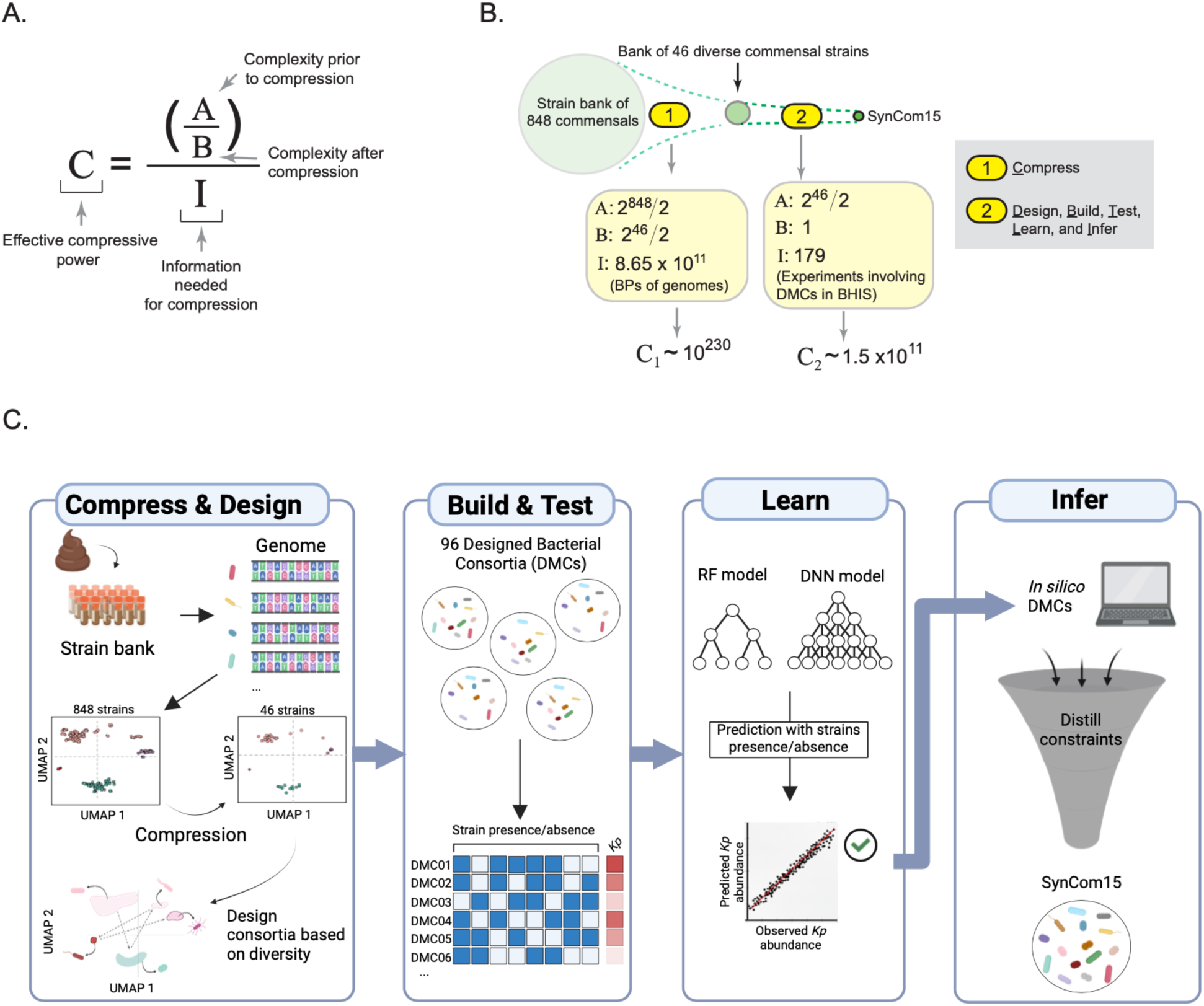
Constraint Distillation: A strategy for the statistical design of synthetic functional consortia from sequenced bacterial strains. **(A)** Computing effective compressive power. **(B)** Compression from starting point—bank of 848 gut commensal strains—to SynCom15 required two sets of information. The first was the genomes of all strains to compress the design space from 848 to 46 strains (a total of information content of 8.65 × 10^11^). The second was 179 experiments comprised of evaluating DMCs in BHIS media used in the Design-Build-Test-Learn-Infer steps to engineer SynCom15. **(C)** Schematic of Constraint Distillation defining inputs—genome-sequenced strain bank—and outputs—statistically designed consortium for target function.

We next evaluated the choices that were critical for the efficiency of Constraint Distillation. The Compress step resulted in 46 strains that recapitulated the diversity of the 848 strains in our strain bank. However, our results showed that DMC46—the consortium containing all 46 strains—was not effective at suppressing *Kp*-MH258 *in vivo*. Thus, simply performing the Compress step was insufficient for engineering a consortium that translated pathogen suppression across environments. The Design step of leveraging genetic diversity to construct 96 DMCs from 46 strains resulted in a substantial reduction in complexity. However, our results from screening 96 DMCs illustrated that merely designing for genetic diversity proved insufficient for engineering consortia that predictably suppressed *Kp*-MH258. Regarding the Test function itself, we found that suppression of *Kp*-MH258 was not explainable by simple heuristics—(i) richness and diversity of consortia, (ii) the presence/absence of individual strains, (iii) or the combination of richness/diversity with the presence of strains previously shown to metabolically overlap with *K. pneumoniae* like *E. coli*. Moreover, the result of inference on our RF model could have yielded a phylogenetically constrained group of strains, e.g. all strains of the same genera. However, this was not the case as SynCom15 remained phylogenetically diverse. Thus, the test function we chose to target—suppressing *Kp-*MH258—was complex enough to require a specific set of diverse, interacting microbes to satisfy suggesting that the experimental efficiency of our process was likely not merely due to a simplicity of test function. These observations shifted our focus to the final two steps of Constraint Distillation: Learning and Inference.

With respect to the Learning step, different types of ML models—RF and deep learning-based neural networks—proved to be useful in statistically learning an input-output relationship with our data. Thus, the specific class of ML model was not critical for the efficiency of Constraint Distillation. However, a specific choice we made in the Learning step was to describe each of the DMCs by their pattern of strain presence/absence. Notably, this choice neglected information about dynamics of the co-culture, physiologic state of the microbes in the culture, interaction with *Kp-*MH258 throughout the time-course of the co-culture, or mechanistic information regarding pathogen suppression. Additionally, our efforts at characterizing the mechanism of action of SynCom15 showed that the supernatant of the co-culture after 120 hours was sufficient for suppressing *K. pneumoniae*, illustrating the importance of the metabolic environment created by SynCom15 towards pathogen suppression. As such, we sought to understand whether a generative model of consortia design could be created using the metabolic profiles of DMCs as defined at 120 hours of co-culture with *Kp-*MH258. Our rationale was that pursuing this goal would enable comparing two types of information—strain presence/absence of DMCs versus metabolite content at 120 hours of co-culture—that were leveraged by us to (i) design SynCom15 or (ii) explain how SynCom15 works respectively.

We performed targeted metabolic profiling of DMCs described in **Fig. 1C** in addition to the 60 DMCs that collectively served as the out-of-sample set when co-cultured with *Kp*-MH258 for 120 hours (**Supplementary Table 13A,B**). The metabolic panel used was the same used for characterizing consortia in **Extended Data Fig. 12A**. We trained and validated an RF model and DNN model on the metabolic profile of DMCs to predict *Kp*-MH258 abundance after 120 hours of co-culture (Methods). We found that these models were well trained, exhibiting an r^2^ value of 0.9 and 0.7 respectively with the validation set of data (**Extended Data Fig. 14A**, **Supplementary Table 13C**). However, these models failed to generalize to out-of-sample communities (**Extended Data Fig. 14B**, **Supplementary Table 13D**). Our finding that the models performed well on the validation dataset but were unable to generalize out-of-sample pointed to model overfitting. We emphasize that this result does not suggest that metabolomics is uninformative. Indeed, multiple factors could limit its generative utility in our data, including incomplete coverage of relevant metabolites, the use of a single endpoint (e.g., 120 hours of co-culture) rather than a time-course based input, and missing important variables like supernatant pH—a trait we found was crucial for explaining the mechanism of pathogen suppression by SynCom15. However, these results provided us the unique opportunity to further understand Constraint Distillation as a process: what statistical characteristics of an input space that describes communities enable a generative design model to extract beyond the training ensemble?

We therefore examined the eigenspectrum of variation in the two input spaces used: strain presence/absence and metabolites. Examination of each eigenspectrum with respect to the capacity to suppress *Kp*-MH258 revealed a stark difference between the way information was encoded in the two spaces. In the strain presence/absence space, the top eigenvectors were the most strongly associated with *Kp*-MH258 suppression; additionally, this association decayed with eigenvector rank (**Extended Data Fig. 14C**, **Supplementary Table 13E**). This alignment between the statistical structure of variation and our target function concentrated in a few high-variance, low-noise directions, yielded a low-dimensional coordinate along which suppression is ordered thereby defining conditions where ML models can generalize. In the metabolite space, this alignment was absent. A high-variance mode (the second eigenvector harboring >20% variance) showed little association with suppression while multiple low-variance modes (harboring <5% variance each) were more associated with suppression (**Extended Data Fig. 14D**, **Supplementary Table 13F**). This observation provided an explanation as to why overfitting was occurring: models fit the training data by over-indexing on fluctuant, low-variance modes but these modes are unstable across new communities thereby degrading out-of-sample performance.

Thus, our results shed light on specific critical qualities of Constraint Distillation—a series of sequential steps that took as input the genome sequences of strains and output a statistical model of targeted, function-specific consortia design that directly led to the construction of SynCom15 (**Fig. 7C**). Namely, the capacity to create a useful generative model depended on describing the 96 DMCs in a way that enabled informational resonance between the statistical structure of DMCs and variation in the target function. When statistical signal associated with function is concentrated in the top variance modes of the input representation, learning models can become generative. When statistical signal associated with function resides in low-variance modes, models may fit but fail to generalize.

## Discussion

We have engineered a defined, synthetic microbiome, SynCom15, that (i) suppresses *Kp*-MH258 (a clinical MDR strain of *K. pneumoniae*) across diverse environments, (ii) matches the efficacy of a whole stool transplant in a pre-clinically relevant *in vivo* mouse model of infection, and (iii) is compositionally distinct from natural human gut microbiomes. We achieved this using Constraint Distillation—a process that is agnostic to bacterial interactions, strain phenotypes, data from cohort studies, qualities of natural microbiomes, or any mechanism of action. Instead, Constraint Distillation comprised learning a map of taxonomic structure to desired function from observational data alone to yield SynCom15. A plausible mechanism by which SynCom15 suppressed *Kp*-MH258 was through fatty acid production and acidification of the environment. This mechanism was shared across suppression of multiple, phylogenetically distinct gut pathogens including strains of *E. coli* and *C. difficile.* A sensitivity analysis of the Constraint Distillation framework demonstrated that describing consortia by their pattern of strain presence/absence was critical for achieving an accurate generative model of design. This was because of informational overlap between the structure of taxonomic variation in the 96 engineered consortia that were functionally screened and the target function of suppressing *Kp-*MH258.

We acknowledge clear limitations regarding (i) the effect of SynCom15 and (ii) Constraint Distillation as an approach for engineering consortia. Regarding SynCom15, we note that our *in vivo* model considered only one dietary background—normal mouse chow. However, it is known that dietary background can have a substantial effect on the composition of the microbiome as well as the capacity for probiotic interventions to be efficacious^62,63^. Normal mouse chow is fiber rich; recent studies have demonstrated the limitations of more westernized diets with respect to ameliorating antibiotic-induced dysbiosis^64^. Additionally, our *in vivo* model also only spanned a single genetic background—C5J/Bl6. These experimental choices limit the extent to which SynCom15 can be considered a generally useful consortia for rescuing dysbiosis of the gut microbiome. It will be important for future efforts to vary such environmental variables that likely play an important role in determining the efficacy of a live biotherapeutic intervention. Regarding Constraint Distillation, our results showed that we could learn a generative model of consortia design towards *K. pneumoniae* suppression using 96 synthetic consortia as an initial ensemble. Importantly, this test function could be evaluated using *in vitro* culture. For test functions that involve extra-enteric effects in the host (e.g. the capacity of the gut microbiome to influence the tumor microenvironment, gut brain axis, immune system, or other organs), we note that assay throughput may become a significantly limiting factor. To demonstrate that Constraint Distillation can engineer synthetic microbiomes functions that, at baseline, require murine models, it will be important to further develop theoretical frameworks that enable designing consortia in a more informed manner such that the subsequent Build-Test/Learn-Infer steps can be rapidly iterated upon if necessary.

As synthetic ecology efforts continue to evolve within application to the human gut microbiome, we contrast the composition and functionality of our consortium with other consortia that have previously been created as surrogates for ‘healthy’ microbiomes. Recent studies have emphasized the power of large consortia, comprising greater than 40 strains, for pathogen suppression and other functions associated with a healthy, diverse gut microbiome^11,27^. One rationale for such an approach is to maximize the functional potential of a consortium by simply adding more diverse strains. Our data demonstrate that it is possible to create a synthetic consortium that is sparse—comprising only 15 members—yet can clear *K. pneumoniae* from the gut as well as rescue the effects of antibiotic-induced dysbiosis. Moreover, our data illustrate that a larger consortium comprising more gut commensals that is inclusive of SynCom15—DMC46—was in fact less efficacious with respect to pathogen clearance, thereby illustrating the limitation of using larger commensal consortia as therapeutics. Instead of directly inoculating the gut with the large diversity of strains that may be present in a whole stool sample, the intervention of SynCom15 appears to be more indirect—allowing the rescue of bacterial diversity through creating an environmental milieu for commensal growth. We reason this is the property underlying the sustained pathogen resistance we see well beyond gavage of SynCom15. In this sense, SynCom15 represents a qualitatively different strategy for repopulating the gut microbiome after perturbation relative to a full stool transplant: by creating an appropriate gut environment, SynCom15 promotes the re-diversification of the gut ecosystem. From a more practical perspective, to realize the value of synthetic ecology it will be important to attempt synthesizing consortia as live biotherapeutics and establishing clinical trials for their efficacy. Given the challenges associated with the regulatory landscape and manufacturing of live biotherapeutics, having a defined mixture of 15 strains constitute the investigational new drug (IND) as opposed to much larger consortia or other types of live biotherapeutic constructs that are undefined (e.g. stool transplants or purified contents from healthy donor stool) will be a substantial benefit in ultimately transitioning synthetic consortia to clinical settings.

Due to the complexity of microbiomes, there have been many recent efforts that augment design with statistical formalism^21–24,65,66^. These efforts have used some level of biological behavior to inform the statistical approach. Inclusion of these behaviors—metabolic qualities, growth qualities, presumed architecture of interactions via consumer-resource models, interaction characterization, functional screens—constitute the equivalent of a forward model imposed by the experimenter. We note, this is not statistical design. Statistical design involves solving the inverse problem—learning a model from observation alone. In other fields of biology and science spanning protein design, material science, neuroscience, and complex systems-based macro-ecology, implementation of statistical design involves (i) creating an ensemble of systems reflecting extant diversity, (ii) learning a probabilistic model of constraint, and (iii) using the model to generate synthetic systems^37–40,67–71^. In our work, we closely mirrored this approach with the difference being that instead of collecting microbiome data from the wild and therefore representing extant microbiome diversity, we created an ensemble of microbiomes using genetic diversity as merely a sampling strategy. Thus, our approach fell directly within the purview of statistical design. What are benefits of not including priors and only employing the strategy of statistical design? First, this approach filters priors that are not general. For instance, our data show that the previously published prior of consortia diversity and presence of *E. coli* to specify *K. pneumoniae* suppression could not explain our results. Second, statistical design makes possible the discovery of mechanism rather than anchoring on prior mechanisms as routes of design. In our work, we first constructed SynCom15 then evaluated its mechanism of action rather than the reverse. Third, learning a statistical model of design opens the possibility of truly synthetic ecology—the creation of products that are unnatural and thereby expand the functional space beyond what nature has created. We showed that SynCom15 was not detectable within natural human gut microbiomes and encoded different mechanisms by which it achieved pathogen suppression across phylogenetically diverse gut pathogens. While future work will test the limits of generality with respect to using statistical design for engineering consortia against other functions, our work has demonstrated that such a strategy is viable, efficient, and can lead to the creation of novel synthetic products.

As increasingly more phenotypes are associated with the gut microbiome, we pose that a useful test for judging the strength of association is through design of synthetic consortia that affect the phenotype. We note that Constraint Distillation is agnostic to the specific function being optimized. The ‘Test’ module (‘T’) can be adapted to any phenotype for which there is an assay. Because implementing Constraint Distillation to engineer synthetic ecological systems does not depend on natural community composition, known mechanisms, or pre-existing pathway annotations, this approach offers a potentially general route to the statistical design of synthetic microbiomes. More broadly however, Constraint Distillation is, in theory, also agnostic to the specific type of ‘part’ used to construct a collective whole. That is, given an ensemble of parts (e.g. bacterial strains) described by degrees of freedom (bacterial genomes), the process of Constraint Distillation can be employed to create an emergent system. In this sense, Constraint Distillation offers more than a procedure for specifically designing synthetic microbiomes but could be a more abstract strategy for using collections of heterogeneous parts to engineer synthetic emergent systems—systems where higher-order, unintuitive interactions between parts create a functional collective. As physics-based principles for building emergent systems *de novo* do not currently exist, testing this idea across both ecological and non-ecological domains represents an exciting future direction.

Our results challenge prevailing assumptions about what is required to design functional microbial communities for therapeutic effect. Prior studies have emphasized the need for high-dimensional measurements, detailed mechanistic understanding, or complex dynamical models to capture structure-function relationships that govern microbial ecosystems. Community design efforts have typically drawn on a wide array of information sources including: (i) quantitative modeling of interspecies interactions, (ii) knowledge of molecular mechanisms underlying target functions, (iii) pathway-level genome annotations, (iv) human microbiome composition as a blueprint, or (v) down-selection from natural communities exhibiting desirable traits such as pathogen resistance (**Supplementary Table 14**, **Supplemental Discussion**)^10,11,13,16,22–25,27,72^. Moreover, strategies for translating microbiome function across unrelated, diverse contexts have involved learning even more information about environmental differences and context-specific bacterial interactions for consortia optimization^3,73^. Our findings paint a substantially different picture. We find that generative principles of consortia design can emerge from far simpler inputs. A binary encoding of just 96 metagenomically diverse synthetic communities by strain presence/absence coupled with empirical functional measurements was sufficient to achieve two key results: the capacity to train a statistical model mapping community structure with function and guide the construction of a sparse community (SynCom15) that translated successfully to diverse environments. Constraint Distillation proved substantially compressive, navigating a large design space with minimal information relative to the full combinatorial complexity of design. In computational terms, this workflow achieved an efficient search over a high-dimensional space using a low-dimensional, statistically learnable representation. We hypothesize that the success of this approach stems from an underappreciated property of many biological systems: that despite their apparent complexity, biological systems are often governed by low-dimensional rules that can be successfully inferred directly from observational data. Our results regarding the informational resonance between patterns of strain presence/absence and pathogen suppression supports this idea. Placing learned statistical patterns before biological phenomenology or mechanistic understanding has found growing support across biology, including recent breakthroughs in data-driven protein design and the adoption of AI-driven approaches at the scale of genomes and tissues^74–81^. Our results suggest that the same principle may hold for microbial ecosystems.

---

**Extended Data Fig. 1.**
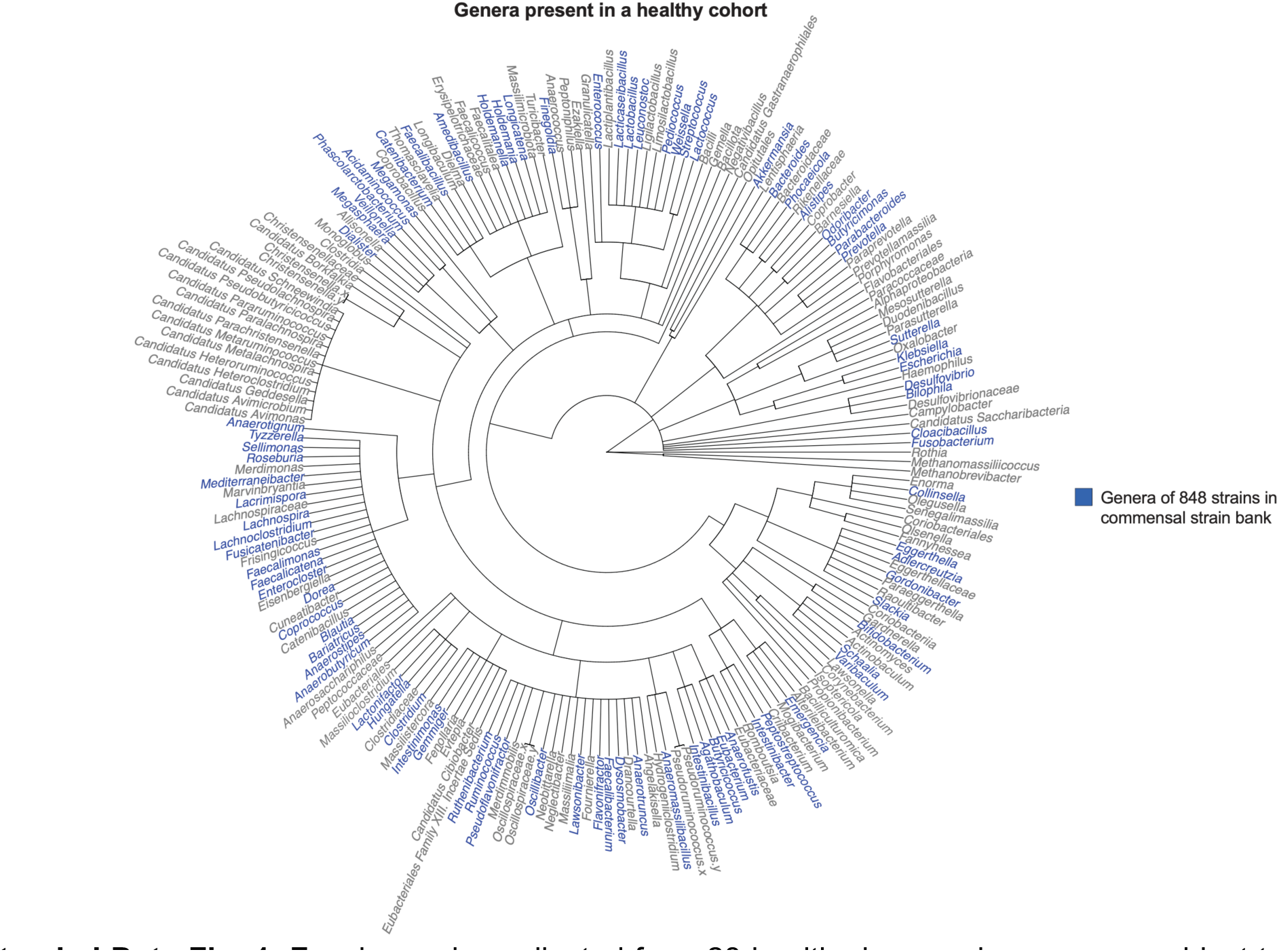
Fecal samples collected from 28 healthy human donors were subject to shotgun metagenomic sequencing. Tree of the genera comprising all fecal microbiomes is shown here. Colored in blue are the distribution of genera observed in our bank of 848 commensal strains.

**Extended Data Fig. 2.**
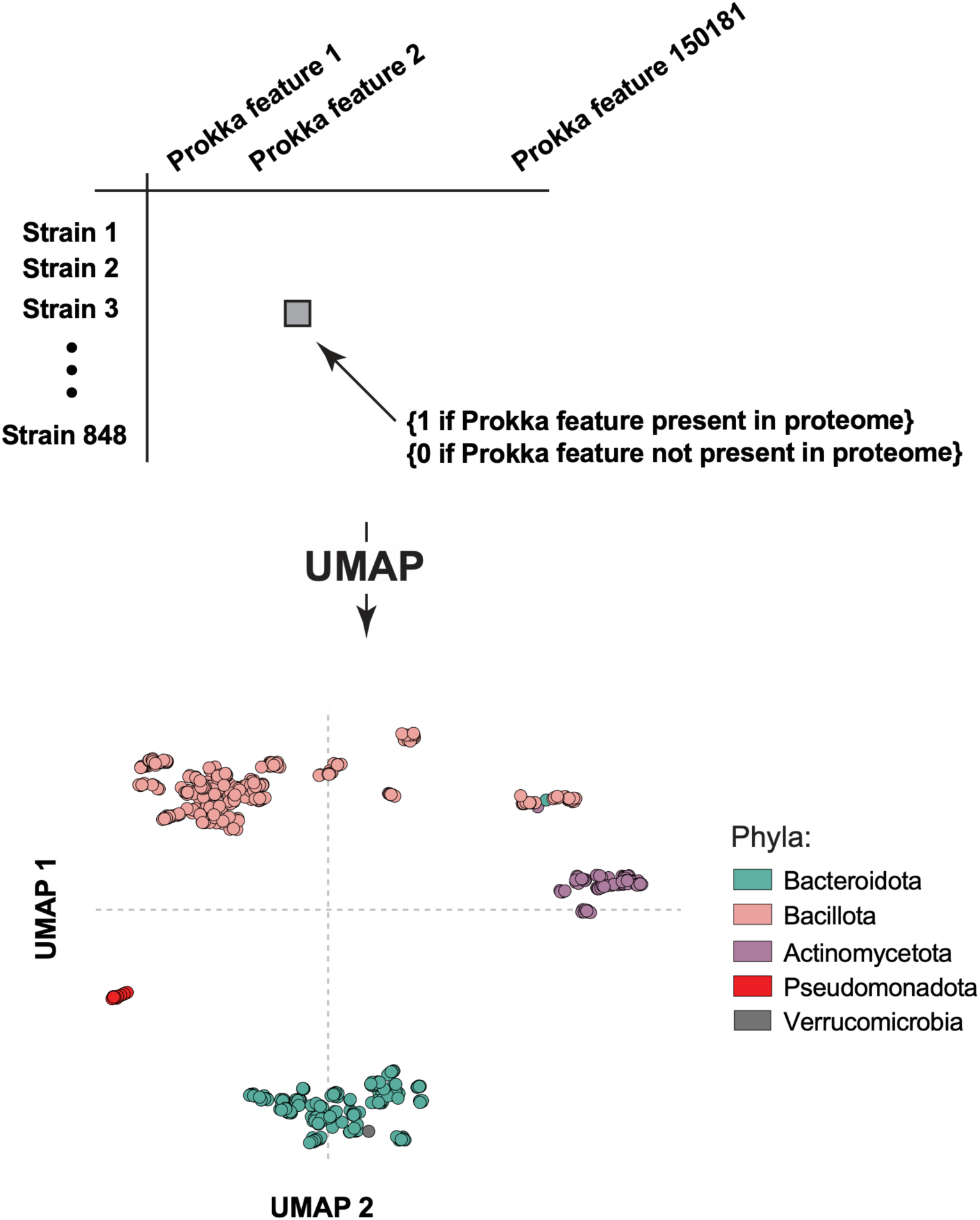
Matrix of strain (rows) by Prokka feature (columns) for 848 gut commensal strains in our strain bank was created where entries are a ‘1’ if the Prokka feature is present in the strain proteome, and ‘0’ if Prokka feature is absent in the strain proteome (top). Matrix was subject to UMAP decomposition resulting in two axes (UMAP1 and UMAP2) used to define clusters of bacteria; each dot in this plot is an individual bacterial strain (bottom).

**Extended Data Fig. 3.**
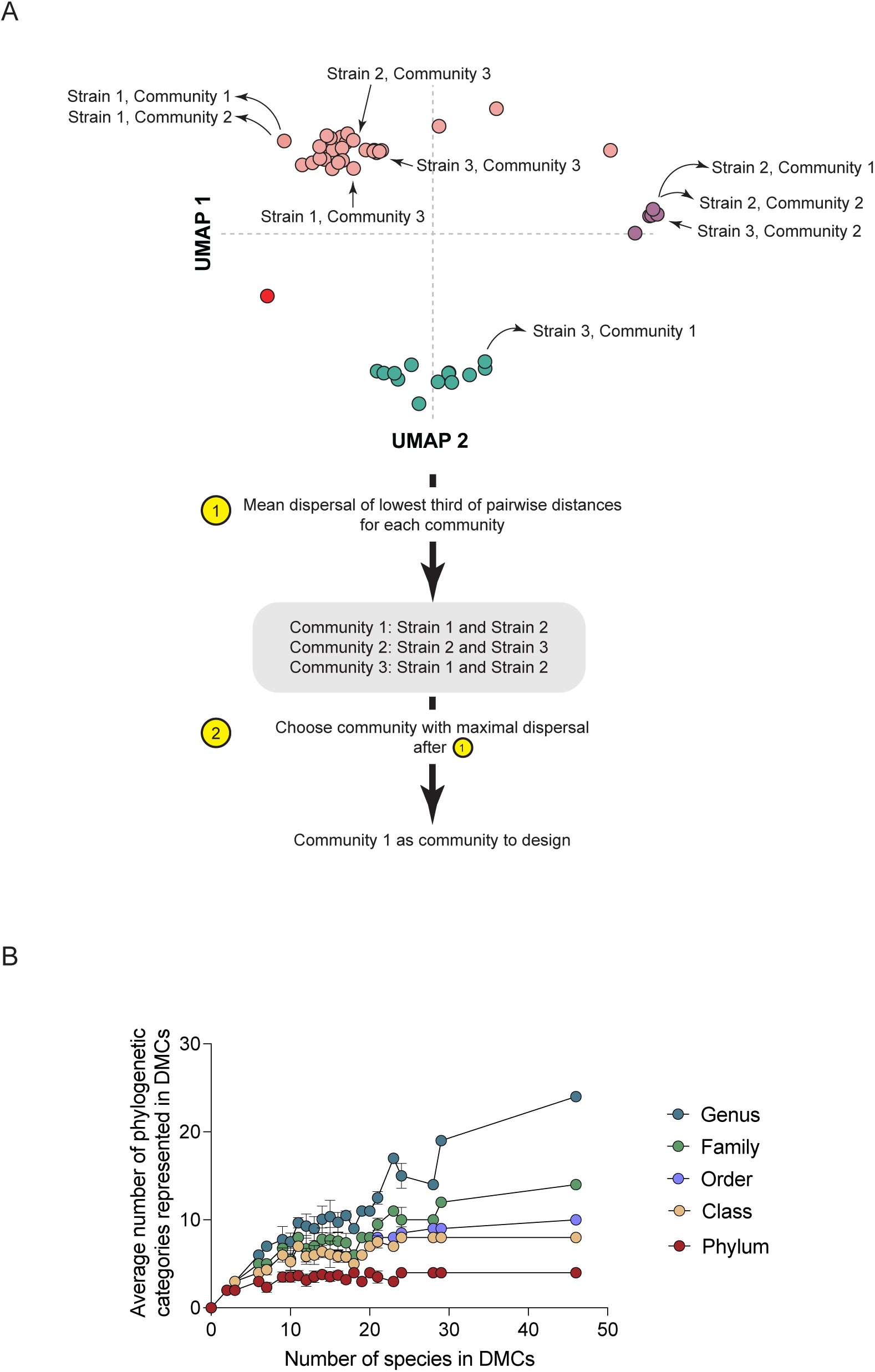
**(A)** Workflow for algorithm used to design synthetic consortia. Communities with three bacterial strains are shown as an example. Given three possible communities that could be created, the first step is to choose the mean dispersal of the lowest third of pairwise distances between strains for each community. In the example shown here, the lowest third is equivalent to the minimum pairwise distance for each community due to the communities being comprised of only three strains (gray box). The second step is to choose the community with the maximal dispersion per Step 1. In the case shown here, ‘Community 1’ would be chosen as a DMC to create. **(B)** Phylogenetic diversity (y-axis) versus number of species in DMCs (x-axis). Each dot is the averaged prevalence of a specific phylogenetic category spanning Phylum, Class, Order, Family, and Genus (see color key) across all DMCs of a specific membership size. The maximum possible prevalence of each phylogenetic category is set by the DMC containing all 46 strains used to engineer DMCs (the right-most set of dots).

**Extended Data Fig. 4.**
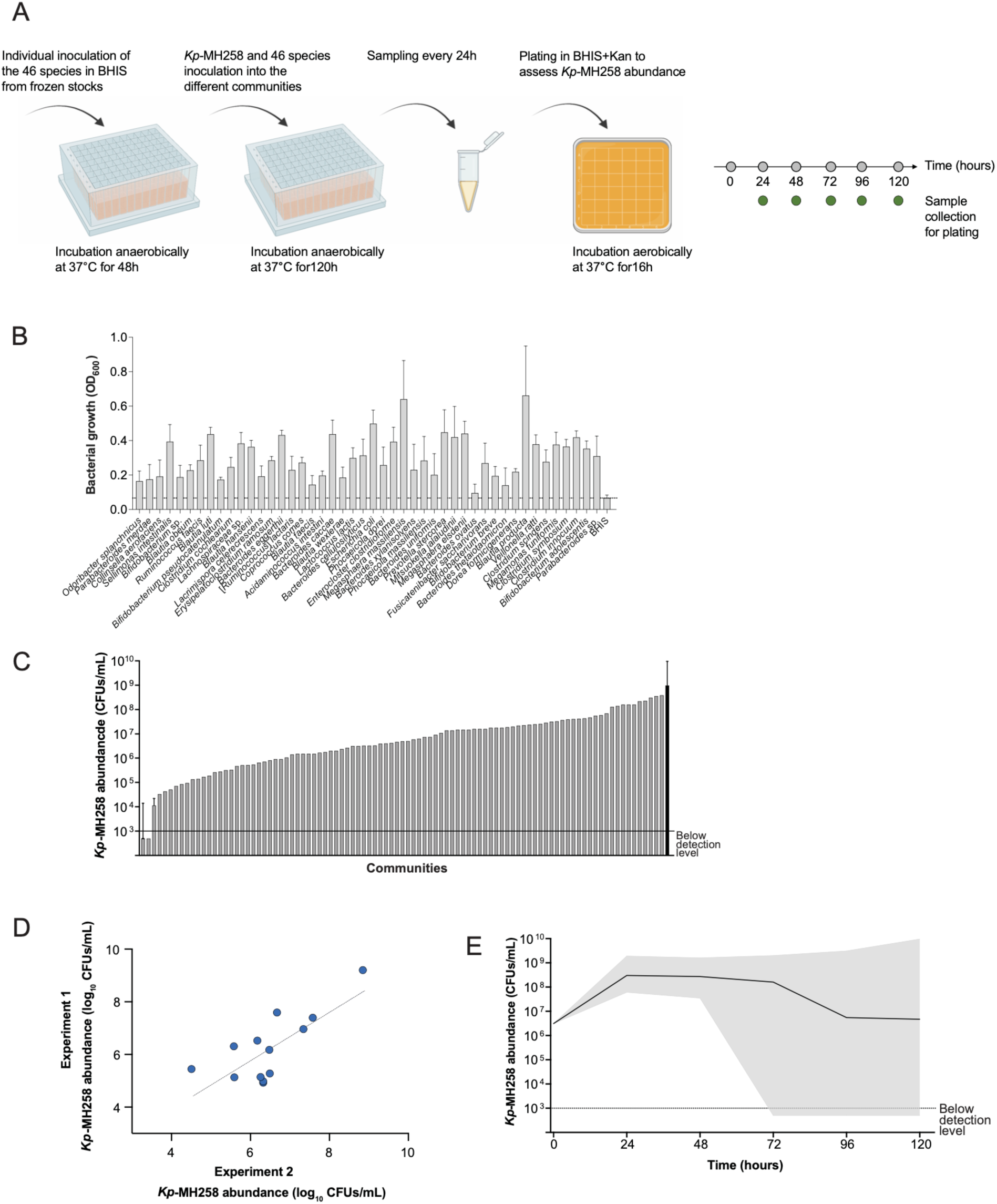
(A) Workflow for evaluating clearance capacity of DMCs for *Kp-*MH258 *in vitro* (‘BHIS’ is Brain Heart Infusion media supplemented with cysteine, ‘Kan’ is kanamycin). (B) Bacterial growth (OD_600_, y-axis) of each of the 46 strains comprising the columns of the matrix in Fig. 1C grown in BHIS media for 48 hours. Bars represent average values, error bars represent standard deviation. (C) *Kp*-MH258 abundance (y-axis) after co-culture with each DMC (x-axis) for 120 hours where the x-axis is ordered by the most suppressive DMCs (left) to the least suppressive (right) DMCs. (D) Correlation between the suppressive capacity of several different DMCs across two experimental replicates in BHIS media. (E) Time course of *Kp-*MH258 abundance (y-axis) when co-cultured with each DMC described in the rows of Fig. 1C. Solid line represents the median value of *Kp*-MH258 abundance in co-culture across all 96 DMCs at a given timepoint; grey shade represents the distribution of *Kp*-MH258 abundance in co-culture across all 96 DMCs at a given timepoint. ‘Below detection level’ delineates limit of the assay to detect suppressive capacity of a DMC.

**Extended Data Fig. 5.**
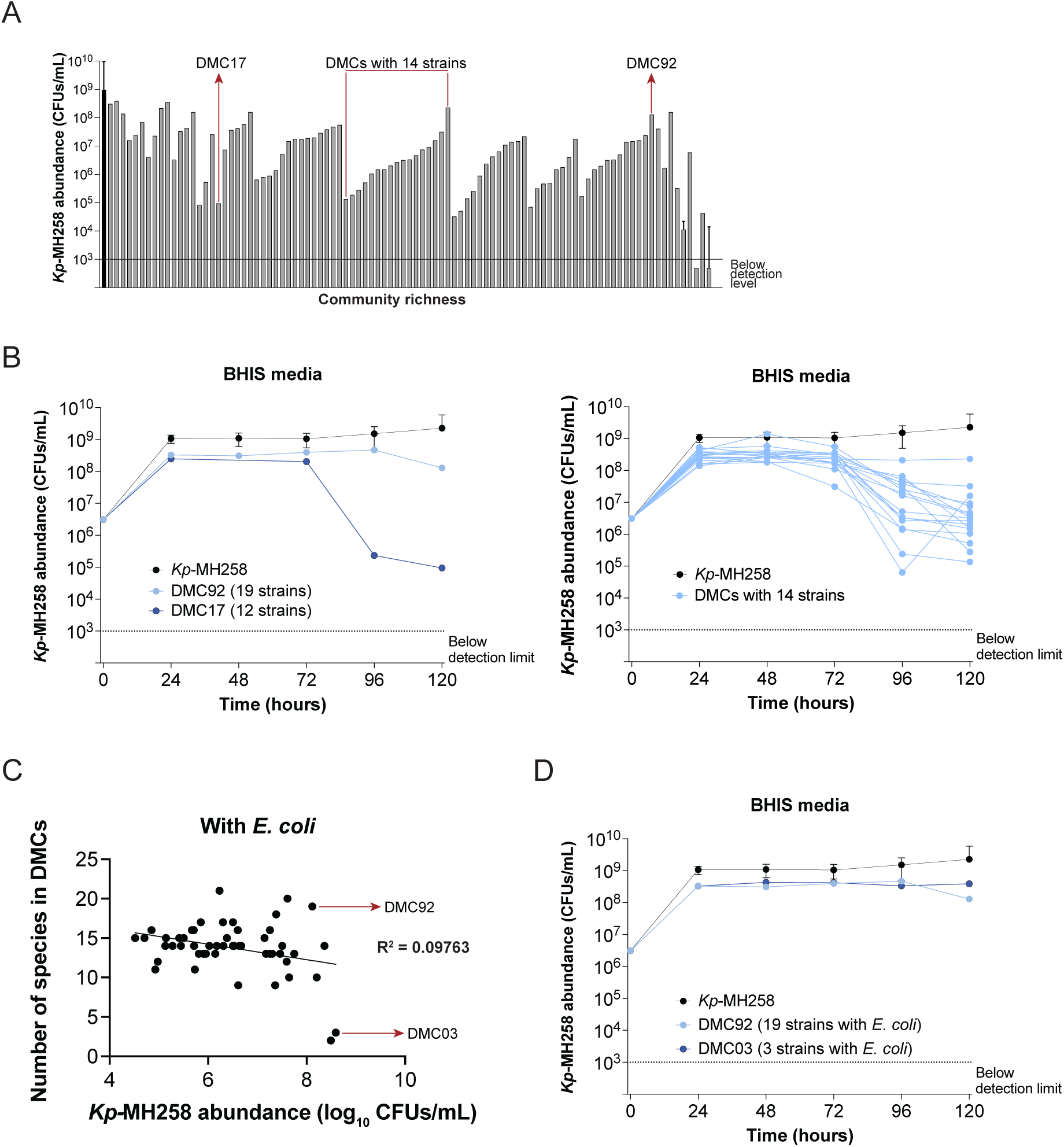
**(A)** Communities ordered from least rich to most rich (x-axis) versus *Kp*-MH258 abundance after 120 hours of co-culture (y-axis). Solid line indicates detection level of the assay; first entry on the x-axis is *Kp-*MH258 abundance in monoculture (black bar). **(B)** Time course of *Kp*-MH258 abundance when co-cultured with DMC92 and DMC17 (left); Time course of *Kp*-MH258 abundance when co-cultured with DMCs containing 14 strains (right). **(C)** DMCs containing *E. coli* as a bacterial strain (dots) are separated by their capacity to suppress *Kp*-MH258 in co-culture (x-axis) and the number of species they contain in their design (y-axis). Highlighted are two DMCs (DMC92 and DMC03) with similar suppressive capacities that represent markedly different degrees of community richness. **(D)** Time course of *Kp*-MH258 abundance when co-cultured with DMC92 and DMC03. In panels B and D, Time course of *Kp*-MH258 is represented by median values and error bars are represented by interquartile range.

**Extended Data Fig. 6.**
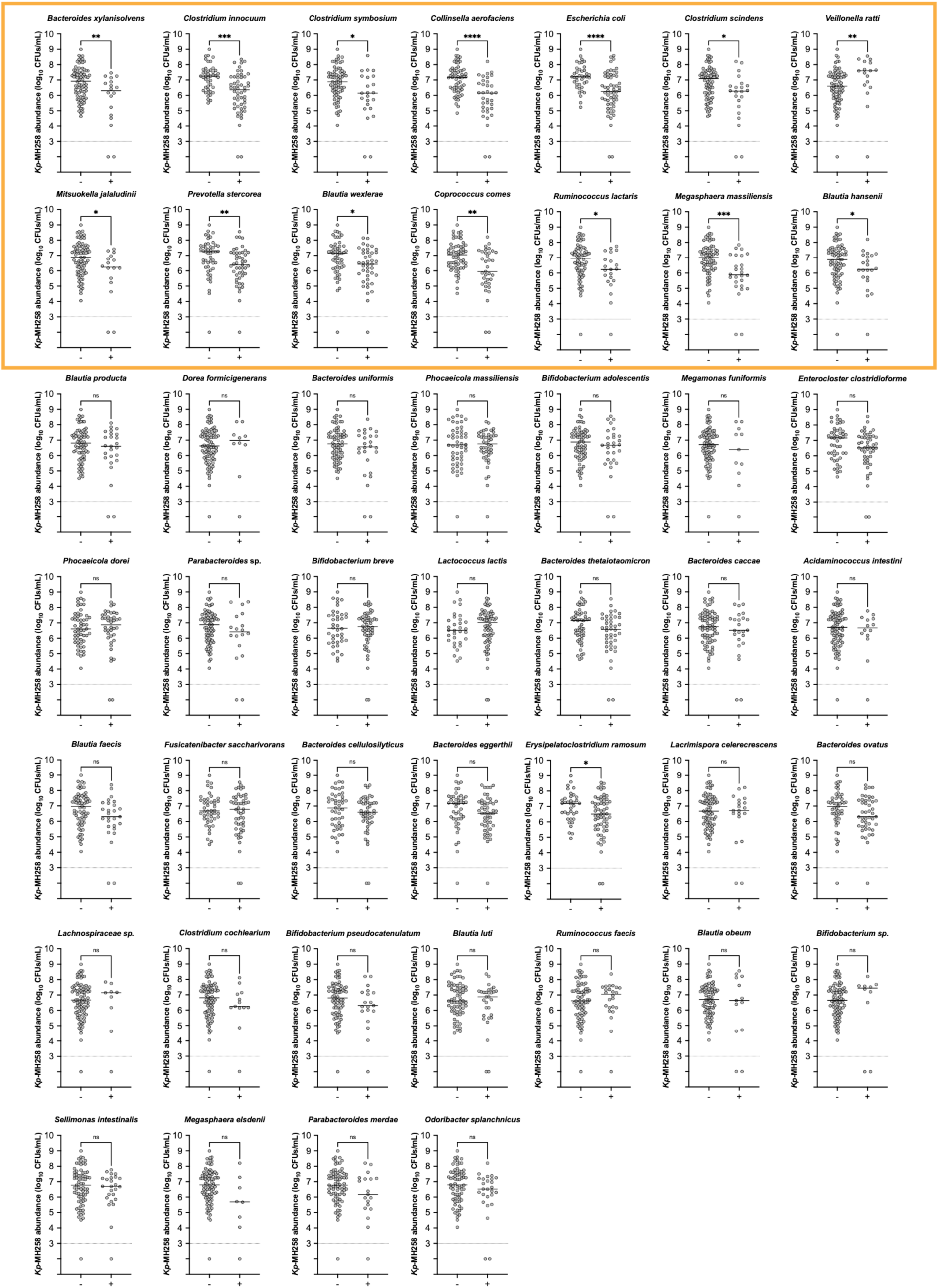
The effect of the presence (‘+’) or absence (‘-‘) (x-axis) of a given strain on the capacity of DMCs (dots) to suppress *Kp*-MH258 (y-axis). Each panel is stratified by each of the 46 strains comprising all DMCs; solid black line indicates median value of *Kp*-MH258 abundance. Yellow box distinguishes set of strains associated with a statistically significant relationship between their respective presence/absence and the capacity for DMCs to suppress *Kp*-MH258. Statistical significance determined by Mann-Whitney test.

**Extended Data Fig. 7.**
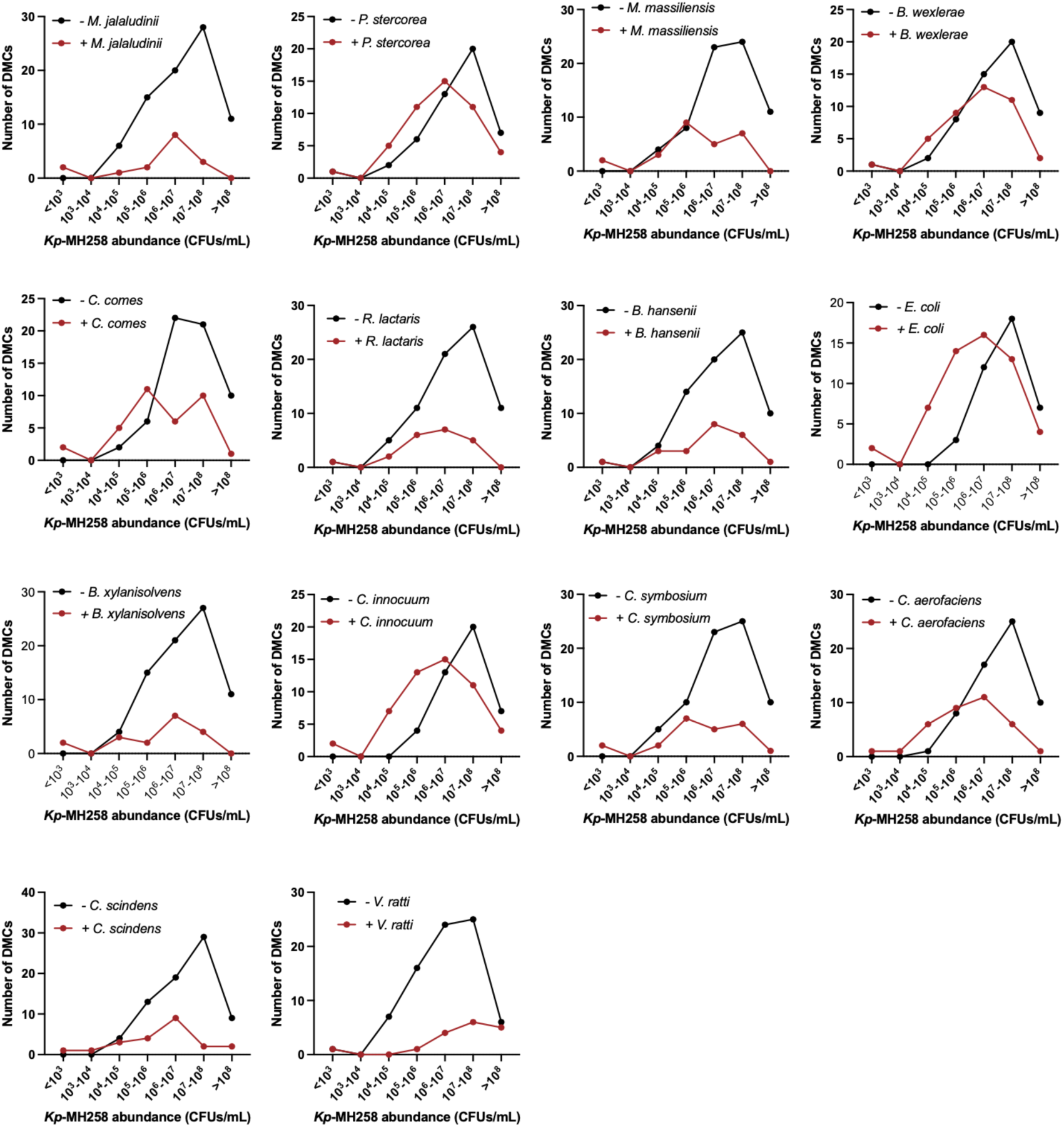
For each of the 14 strains whose individual presence (red line)/absence (black line) was associated with a statistically significant difference in the capacity of DMCs to suppress *Kp*-MH258, histograms of the number of DMCs (y-axis) across the measured range of suppressive capacity (x-axis) are plotted.

**Extended Data Fig. 8.**
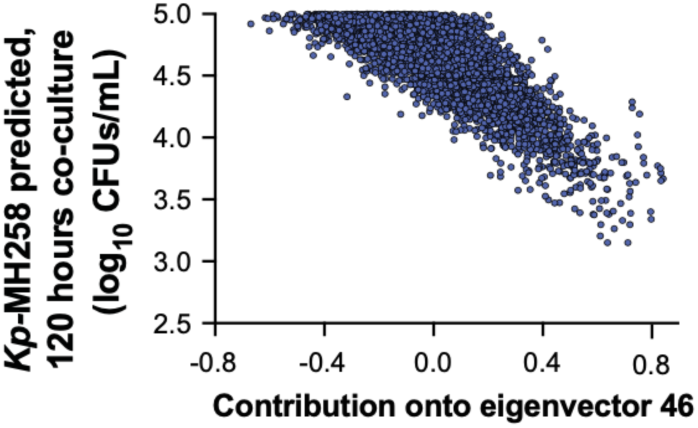
Loading of each of the 5,752 DMCs generated from the RF model (blue dots) onto eigenvector 46 (x-axis) versus predicted abundance of *Kp*-MH258 at 120 hours of co-culture (y-axis).

**Extended Data Fig. 9.**
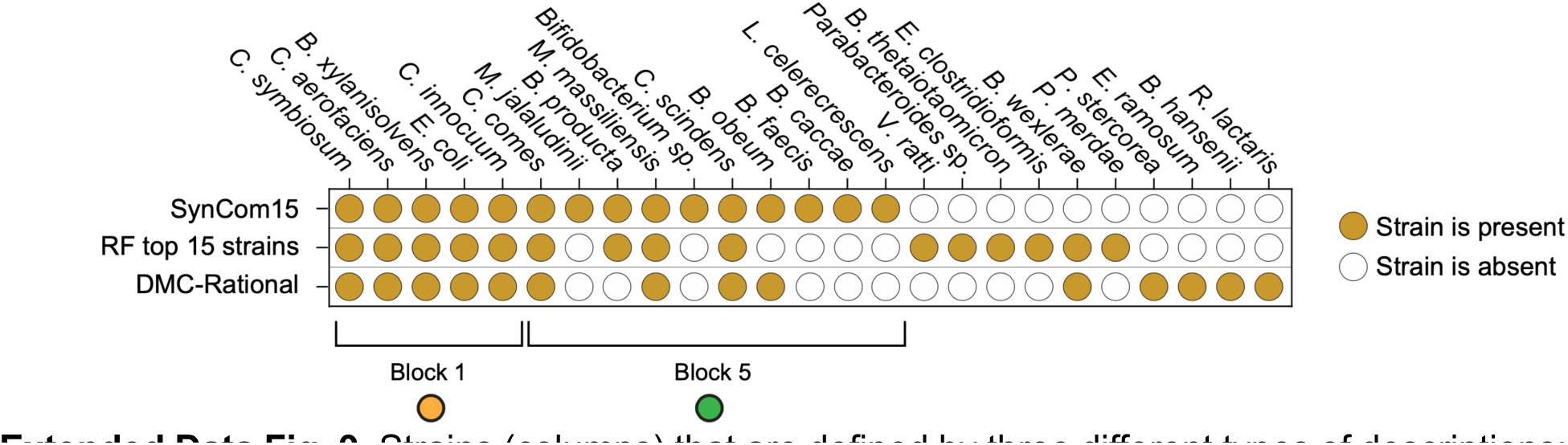
Strains (columns) that are defined by three different types of descriptions: present in SynCom15, present in the top 15 strains as determined by feature importance score of our trained Random Forests (RF) model, present in the DMC-Rational consortium.

**Extended Data Fig. 10.**
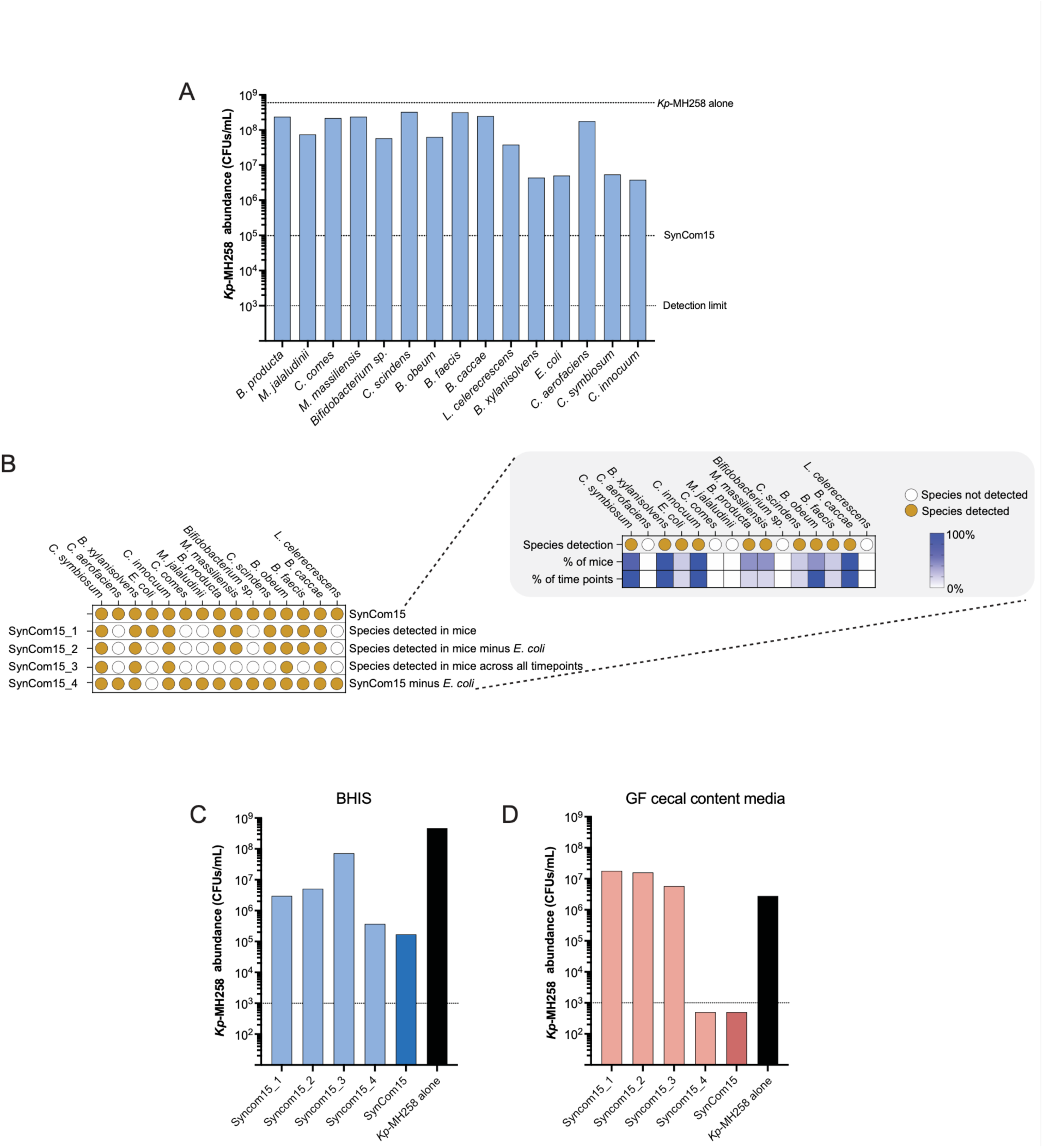
(A) *Kp*-MH258 abundance (y-axis) after co-culture with each SynCom15 strain (x-axis) individually for 120 hours in BHIS media. (B) Construction of several variants of SynCom15 (rows) defined by the presence (yellow dot) or absence (white dot) of SynCom15 strains (columns). Inset illustrates strains that were detected in mice. First row indicates of inset that the strain was found in any mouse at any given timepoint after gavage of SynCom15 post antibiotic treatment and pathogen infection; second row of inset indicates percent of mice in which strain was observed; third row of inset indicates percent of timepoints in which strain was observed. (C,D) *Kp-*MH258 abundance (y-axis) after 120 hours of co-culture with each of the SynCom15 variants shown in panel B in BHIS media (panel C) and GF cecal content media (panel D).

**Extended Data Fig. 11.**
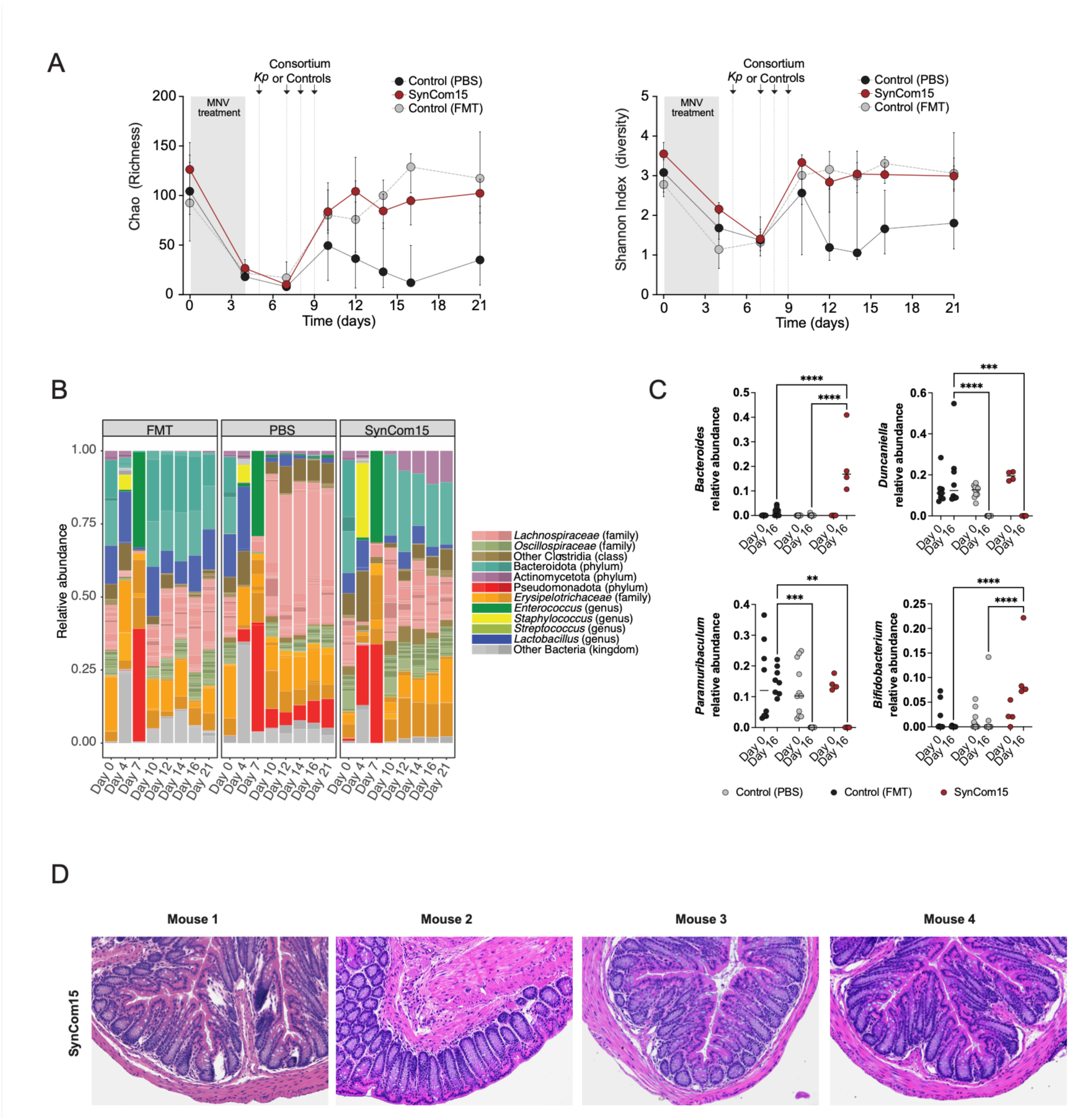
**(A)** Chao and Shannon diversity indices (y-axes) versus time (x-axes) for SPF mice treated with MNV, infected with *Kp-*MH258 (‘*Kp*’) and given PBS, FMT, or SynCom15. Lines represent media values, error bars indicate interquartile range. **(B)** Distribution of average relative abundance for major bacterial phylogenies represented in the fecal microbiota through time (x-axis) for infected mice treated with FMT (left panel), PBS (middle panel), or SynCom15 (right panel). Distributions are defined spanning kingdom to genera-level descriptions. **(C)** Relative abundance of *Bacteroides*, *Duncaniella, Paramuribaculum* and *Bifidobacterium* genera that are differentially abundant amongst infected mice treated with FMT, saline, or SynCom15 prior to antibiotic treatment (day 0) and at day 16 after treatment (equivalent to 11 days after infection with *K. pneumoniae*). Statistical tests performed are two-way ANOVA; **p < 0.01; ***p < 0.001; ****p < 0.0001. **(D)** Hematoxylin and eosin stain of colon of infected mice given SynCom15.

**Extended Data Fig. 12.**
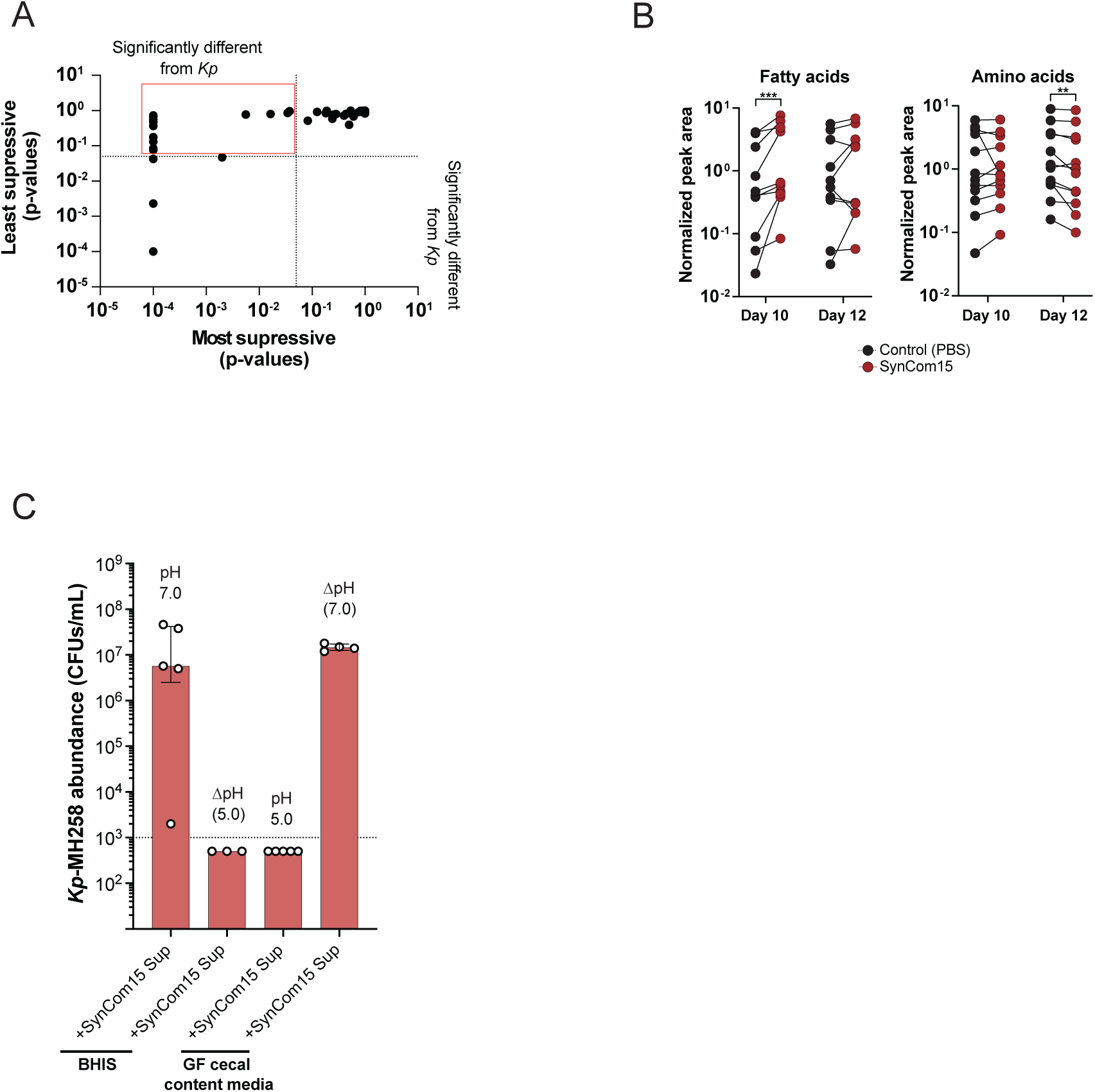
**(A)** p-values for differential enrichment of metabolites were computed for the five DMCs that were the most and least suppressive of *Kp*-MH258 compared to *Kp*-MH258 in monoculture in BHIS media (two-way ANOVA). Each dot in the plot is a metabolite; the x-axis defines the range of p-values observed in the most suppressive DMCs; the y-axis defines the range of p-values observed in the least suppressive DMCs. The red box delineates set of metabolites that are most significantly differentially enriched in the most suppressive DMCs but are not significantly enriched in the least suppressive DMCs. **(B)** Distributions of normalized peak areas (y-axes) of fatty acids and amino acids from fecal samples collected on Day 10 and Day 12 of mouse experiment shown in **Fig. 4C** for mice gavaged with saline (PBS) or SynCom15 (maroon). ***p<0.001; Mann-Whitney. **(C)** Abundance of *Kp*-MH258 in BHIS and GF cecal content media when co-cultured for 120 hours with supernatants collected from SynCom15 cultured alone for 120 hours. Supernatant in BHIS media exhibited a natural pH of 7.0 and was then acidified to a pH of 5.0 (ΔpH). Supernatant in GF cecal content media exhibited a natural pH of 5.0 and was buffered to a pH of 7.0 (ΔpH). Bars represent median values, error bars represent interquartile range.

**Extended Data Fig. 13.**
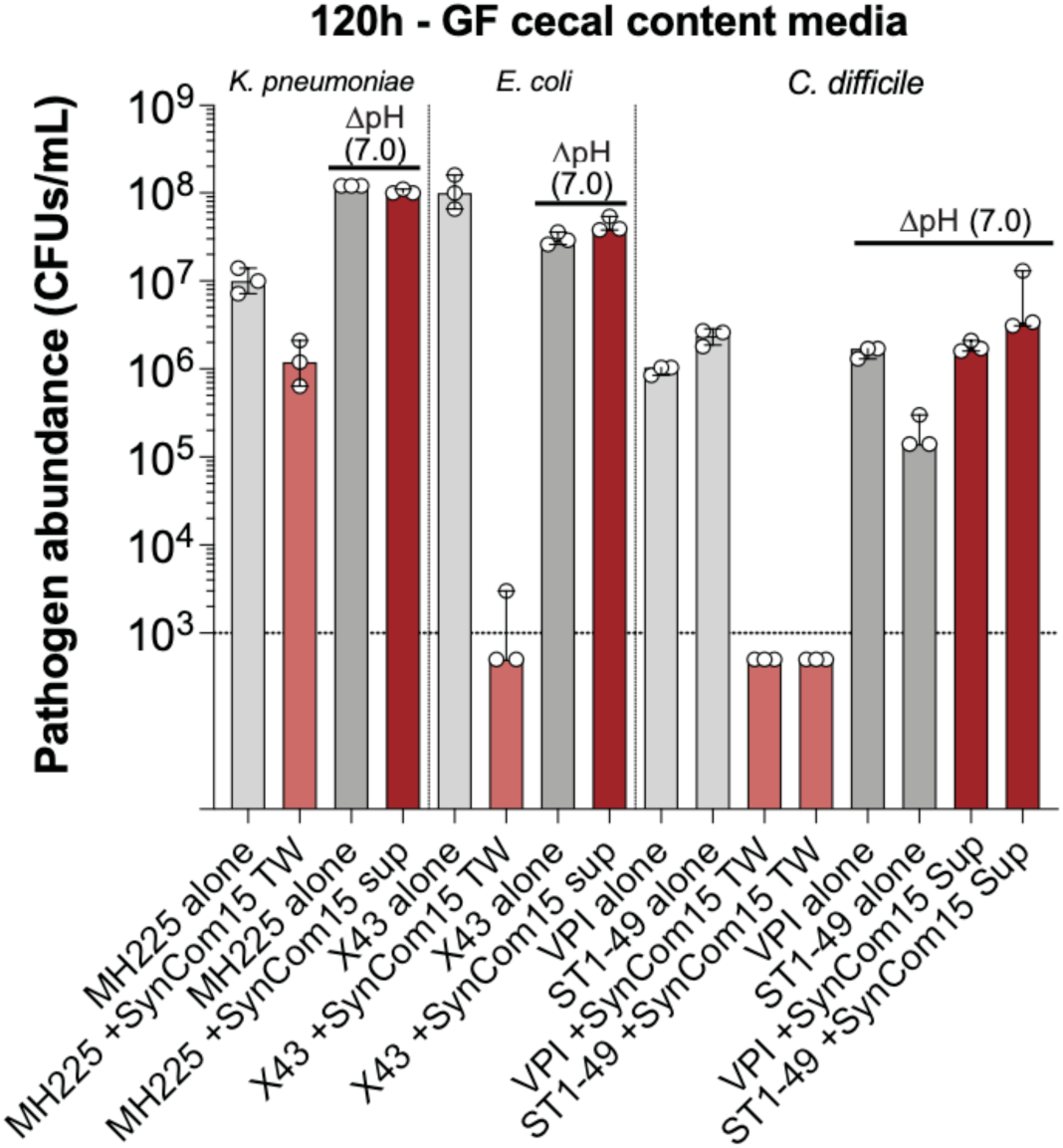
Abundance of various pathogens (y-axis) when monocultured (‘-alone’), or co-cultured in a transwell setup with SynCom15 (‘+SynCom15 TW’) in GF cecal content media at both natural pH and pH adjusted to 7.0. Bars represent media values, error bars represent interquartile range

**Extended Data Fig. 14.**
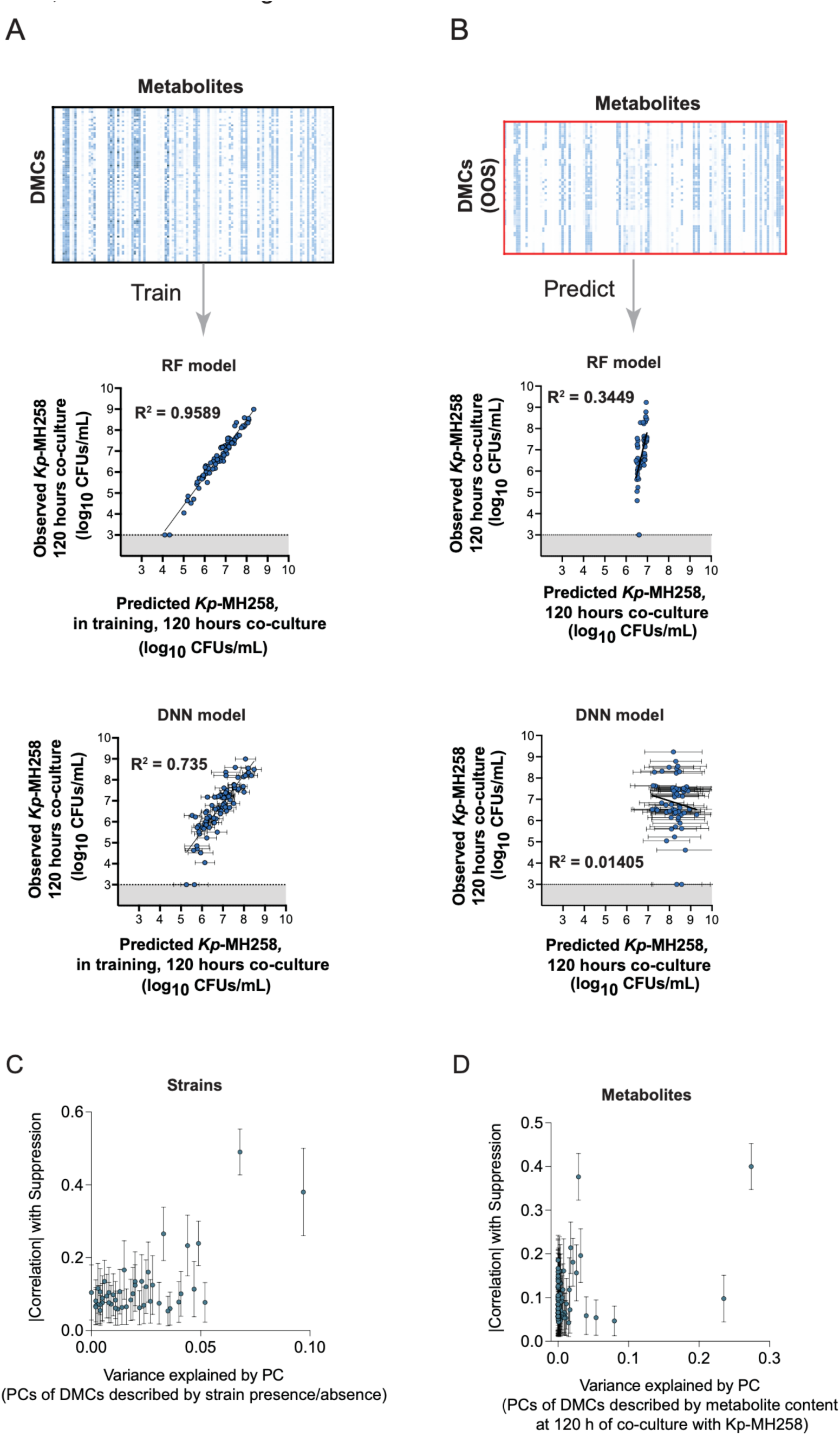
**(A)** Training RF and DNN models of *Kp*-MH258 abundance from metabolite content of DMCs when co-cultured with *Kp*-MH258 for 120 hours. **(B)** Prediction of *Kp*-MH258 abundance from metabolite content of 60 out-of-sample DMCs (see **Fig. 3A**) using trained RF and DNN models from panel A. **(C,D)** Information associated with *Kp*-MH258 suppression (y-axis) versus principal component (‘PC’) (x-axis; ordered from right to left by variance explained by each principal component).

## Methods

### Collection and whole genome sequencing of 848 gut commensals

Fecal samples were obtained from 28 human donors that fell within the age range of 18 to 63 with a median age of 35. Donors were selected as those with no antibiotic use in the past year, no known history of diabetes, colitis, autoimmune disease, cancer, pneumonia, dysentery, or cellulitis at time of consent. Institutions that approved protocols of fecal sample collection were Memorial Sloan Kettering (MSK) and the University of Chicago. Fresh fecal samples were immediately reduced in an anaerobic chamber upon collection and diluted and cultured on various growth media. Agar media types vary, but include any of the following: Columbia Blood Agar, Brain Heart Infusion + Yeast, Brain Heart Infusion + Mucin, Brain Heart Infusion + Yeast + Acetate or N-acetylglucosamine, reinforced Clostridial Agar, Peptone Yeast Glucose, Yeast Casitone Fatty Acids, Defined media M5. Colonies were selected and grown to be sufficiently turbid, 20% glycerol/PBS stocks were created and stored in a -80°C freezer.

Colonies were selected for whole-genome based on pyro-sequencing of the 16S region which provides a rough estimate of genus level designation. For each donor, only colonies that had a sequence identity threshold of less than 99% from CD-Hit (v. 4.8.1) were selected for whole-genome sequencing^82,83^. Bacterial genomic DNA was extracted using QIAamp DNA Mini Kit (QIAGEN) according to manufacturer’s manual. The purified DNA was quantified using a Qubit 2.0 fluorometer. 1000ng of each sample was prepared for sequencing using the QIAseq FX DNA Library Kit (QIAGEN). The protocol was carried out for a targeted fragment size of 550bp. Sequencing was performed on the MiSeq or NextSeq platform (Illumina) with a paired-end (PE) kit in pools designed to provide 1-3 million PE reads per sample with read length of 250 or 150 bp.

Adapters were trimmed off with Trimmomatic (v0.39) with following parameters: the leading and trailing 3 bp of the sequences were trimmed off, quality was controlled by a sliding window of 4, with an average quality score of 15 (default parameters of Trimmomatic)^84^. Moreover, any read that was less than 50 bp long after trimming and quality control were discarded. The remaining high-quality reads were assembled into contigs using SPAdes (v3.15.4)^85^.

Taxonomic classification of the assembled contigs was performed with the following methods: (a) Kraken2 (v2.1.2; (b) full/partial length 16S rRNA gene from each isolated colony’s assembled contigs is extracted and input into BLASTn (v2.10.1+) to query against NCBI’s RNA RefSeq database^86–88^. Top five hits for each query are manually curated to determine an isolate’s identity, with identity and coverage cutoff both at 95%; (c) GTDB-Tk (v1.5.1)^89^. The final taxonomy is determined by the consensus of the three methods. Any colony that did not match initial pyro-sequencing taxonomy or lacked consensus was excluded from the commensal strain bank.

### Dimension-reduction of commensal strain bank

All gut commensal strains were annotated by their Prokka annotations and an alignment was created (848 rows comprising commensal strains, 150181 columns comprising Prokka annotated features)^90^. Each entry in the alignment is a ‘1’ or a ‘0’ indicating the presence or absence of a specific feature in a particular bacterial proteome. Uniform Manifold Approximation and Projection (UMAP) using default parameters (top 10 Principal Components with standard clustering) was used to visualize relative position of each gut commensal strain (see **Extended Data Fig. 2**).

### Algorithm for designing metagenomically diverse consortia from UMAP-based embedding

To design a bacterial consortium comprised of *N* strains, we perform the following steps. Given a UMAP plot of strains, the following algorithm was used to engineer synthetic microbial consortia.

*<u>Step 1</u>*: Create 10,000 communities randomly of size *N*. The ensemble of all 10,000 communities of size *N* is represented as

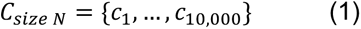

*Step 2*: Each community, *c*_*i*_, is defined by a set of *N* bacterial strains:

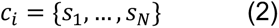

where *s*_*j*_ is strain *j* in *c*_*i*_. Compute all pairwise distances in the UMAP space for all strains in *C*_*i*_. For instance, the pairwise distance between strain 1 and 2 is:

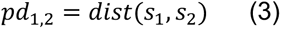

where ‘dist’ is the function that computes the distance between *s*_’_ and *s*_+_ in the UMAP space. We define the distribution of all pairwise distances for *c*_*i*_ as

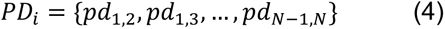

*<u>Step 3</u>*: Order *PD*_*i*_ for a given *c*_*i*_ from largest to smallest values, then compute the mean pairwise distance across the lower 30% of values comprising *PD*_*i*_. We term this value the ‘mean adjusted dispersal’.

*<u>Step 4</u>*: Compute the mean adjusted dispersal for all communities in *C_size N_*.

*<u>Step 5</u>*: Identify the community within the 10,000 communities comprising *C_size N_* with the maximum mean adjusted dispersal. This community is the designed community comprising *N* strains

For downsampling the 848 strains in our commensal strain bank to the 46 strains used to design DMCs (see **Fig. 1A**), the above approach was used where *N* was set to 46 and the UMAP plot used is shown in **Extended Data Fig. 2**. 46 was chosen as the maximum number of strains we were confidently able to manipulate to create bacterial cultures. For creating the DMCs shown in **Fig. 1C**, the above approach was used where *N* spanned a range from 2 to 29 strains and the UMAP plot used is shown in **Fig. 1A**.

A schematic of the process is outlined for a community comprised of three strains in **Extended Data Fig. 3A**.

### Creating the Klebsiella pneumoniae MH258 strain used in experiments

The *Kp*-MH258 isolate was previously described elsewhere^42^. For better in vitro and in vivo selection of this strain, *Kp*-MH258 was transformed by electroporation with pmCherry-sfGFP (86441; addgene).

### Evaluating the effect of DMCs on the fitness of MDR Kp-MH258

The 46 bacterial strains described in **Supplementary Table 1B** were individually inoculated from a frozen stock into 900μL of BHI supplemented with cysteine 0.1% (BHIS) previously reduced. Strains were incubated at 37°C in static conditions for 48h in anaerobiosis to ensure that the most fastidious strains reach stationary phase. *K. pneumoniae*-MH258 sGFP was also inoculated in the same conditions, but only 24h after commensal isolates inoculation due to the fast growth capacity of this species and was incubated for 24h. All strain densities were assessed by taking 100 μL of each culture and measuring OD_600_ in a Biotek Cytation 5. To build all DMCs, isolates were inoculated in 900 μL of BHIS previously reduced in different combinations with an initial OD_600_ of 0.001, so that the densest community reaches a maximum total initial OD_600_ of approximately 0.05. *K. pneumoniae* was added at the same initial OD_600_ of 0.001 (approximately 5 × 10^6^ CFUs/mL) to all DMCs. Cultures were incubated at 37°C in static conditions and anaerobiosis for 5 days. To *assess K.* pneumoniae abundance, 10 μL of each culture were collected daily and homogenized in 90 μL of PBS and serially diluted. Diluted samples were plated in BHIS with kanamycin (50μg/mL). Plates were incubated at 37°C overnight in aerobiosis. GFP expressing *K. pneumoniae-MH258* colony forming units (CFUs) were enumerated. In parallel, 100uL of each culture was also collected to recover the cell phase and the supernatants at 72h, 96h, and 120h. These samples were stored at -80°C to be later processed for shotgun metagenomics and metabolomics.

### Training additive linear regression model on strain presence/absence for suppressing Kp-MH258

We used the LinearRegression module, available under scikit learn (sklearn.linear) python package^91^. The intercept option was set to true which allowed the model to have a non-zero linear intercept if necessary for quality fit. The input data (independent variables) was a vector of 46 1’s and 0’s as shown in the matrix displayed in **Fig. 1C** corresponding to the pattern of presence-absence for each DMC. The dependent variable for model training was the *Kp-MH258* log abundance value for each of these 96 experiments. Both independent and dependent variables associated with the original 96 experiments were used to train the linear model’s coefficients and intercept. After training, we predicated the *Kp*-MH258 log abundances for these 96 experiments. The R^2 value and the Pearson correlation between the prediction and actual abundance values were 0.844 and 0.92 respectively. We extracted and recorded this model’s regression coefficients for each strain. The larger magnitude of this value means a larger effect on the *Kp-MH258* log abundance.

### Animal protocol and husbandry

All mouse experiments were performed in accordance with and approved by the Institutional Animal Care and Use Committee of the University of Chicago under protocol 72599. Male specific-pathogen-free C57BL/6J mice, aged 8 weeks to 10 weeks, from Jackson Laboratories were used for all experiments. Mice were kept within a facility that maintained a 12 hour light and 12 hour dark cycle and controlled humidity (30–70%) and temperature (68–79 °F). Mice were housed in sterile, autoclaved cages with irradiated feed (LabDiets 5K67) and acidified, autoclaved water upon arriving at the on-site mouse facility. Mouse handling and cage changes were performed by investigators wearing sterile gowns, masks and gloves in a sterile biosafety hood. Mice were cohoused with their original shipment group until starting the experiment.

### Preparation of mice stool samples for fecal microbiota transplant (FMT)

Fecal samples from 15-20 mice SPF mice from different cages (to increase sample diversity) were collected to a 50 mL tube. Samples were transferred immediately to the anaerobic chamber (anaerobic exposure was kept under 30 min). Samples were dissolved in 1 mL of PBS 20% glycerol 0.1% cysteine (previously filtered and reduced) per fecal pellet (1mL per ∼20 mg of fecal sample) using a mechanical pestle and vortexing. Samples were aliquoted in cryovials and stored -80°C until use.

### An in-vivo mouse model of MDR Kp-MH258 infection

C57BL/6J male at 8-10 weeks of age were singly housed and placed under an antibiotic regime (0.25g MNV – metronidazole, neomycin, vancomycin) in the drinking water (day 0). Four days later, antibiotic treatment was halted and mice were placed on normal acidified water (day 4). Cages and food were also changed. On day 5 all mice were gavaged with 100μL of PBS containing 500 CFUs of *Kp-*MH258, prepared as previously explained. On days 7, 8, and 9 mice were gavaged with 100uL of either selected defined bacterial consortia, a fecal microbiota transplant from naïve healthy mice, or PBS. Fecal samples were collected on days 0, 4, 7, 10, 12, 14, 16, and 21 (final day of the experiment) for 16s rRNA sequencing and on day 10 and 12 for metabolomics. These were immediately place on dry ice after collection and later stored at - 80°C. To assess for *Kp-*MH258 levels, fecal samples were collected on days 7, 10, 12, 14, 16, and 21. Fecal samples were homogenized in 1mL of PBS and serially diluted. Undiluted and diluted samples were plated in BHIS and kanamycin (50μg/mL).

### Training an RF model and a deep learning-based model on DMC strain presence/absence as input and Kp-MH258 abundance at 120 hours of co-culture with DMC as output

For training the RF model, we used a RandomForestRegressor, available with scikit-learn python package^91^. Tree Depth was set to 12 levels per tree, the number of trees was set to 100, and the maximum number of features was set to “sqrt” (square-root of the number of strains total). Out-of-bag error was measured by a combination of R^2 (where numbers less than 1 indicate more error) and Mean Squared Error (where larger numbers indicate more error). To train and validate our model, we randomly split our dataset into 90% training and 10% true-out-of-sample 100 times. The input data was a vector of 46 1’s and 0’s as shown in the matrix displayed in **Fig. 1C** corresponding to the pattern of presence-absence for each DMC. In each iteration, the RandomForestRegressor was fit to the training set via 6-fold cross validation. Cross-validation accuracy was measured through Pearson Correlation. The true out of sample set was then predicted, and prediction accuracy was measured by computing Mean Squared Error and Pearson Correlation of the predicted versus measured *K. pneumoniae* abundances after 120 hours of co-culture with the DMC. Feature Importance Scores for all features were observed and stored. This process was repeated 100 times, and prediction accuracies and feature significance scores were averaged. An additional RandomForestRegressor model was then trained on the entirety of the dataset with 6-fold cross-validation. Cross-validation accuracy was measured by calculating Mean Squared Error, Pearson Correlation, and R^2. Averaged prediction accuracies and feature significance scores were used to estimate prediction error.

For training the deep learning-based model (‘deep neural network’, DNN), we used the same input data format as described above. We next created 5000 Neural networks using PyTorch each with two hidden layers (50 neurons going to 10 neurons) and a single output neuron tasked with predicting the *Kp-*MH258 log abundance values. For each neural network, we first performed a random 80% -20% training-validation split of the original 96 experiments. This neural network was trained on its training split using a dynamic momenta-dependent backpropagation algorithm where the gradient is calculated on the training split’s loss alone while the momentum parameter is reduced when the loss function on the validation split stagnates. This momentum-based scheme allows for smooth (non-oscillatory) and faster training and reduces the chance of our neural networks getting stuck in local optima.

### Defining SynCom15 through distillation of the RF model

#### Isolating eigenvector most associated with Kp-MH258 clearance

The matrix in **Fig. 3B** was subject to Singular Value Decomposition (SVD) resulting in 46 principal components of data-variance (eigenvectors). We found that the first principal component (PC1) was significantly associated with community complexity (**Supplementary Table 7B**). To isolate the effect of *Kp-*MH258 clearance from community size, we first performed a series of steps to ‘regress out’ the effect of community size. First, let ***x***_*i*_ be the community size of DMC *i*. Let ***y***_***i***_ be the predicted *Kp-*MH258 clearance from the RF model for DMC *i*. A linear model is then created regressing community size against *Kp-*MH258 clearance taking the form:

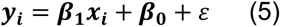

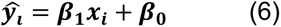

where ***y***6_1_ is the *Kp-*MH258 clearance of DMC *i* as a function of its size. The residuals of this linear model are given by

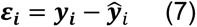

where *ε*_*i*_ is the degree of *Kp-*MH258 clearance of DMC *i* after removing linearly modeled information related to the size of the DMC. All principal components were regressed against 𝒓_*i*_and principal component 46 (PC46) was found to be the most significantly associated with predicted, residualized *Kp-*MH258 clearance (**Supplementary Table 7C**).

#### Defining the matrix in **Fig. 3C**

Let 𝒖 ∈ ℝ^23^ be the vector of projections of each strain on eigenvector 46 of the matrix defined in **Fig. 3B**.

Let *s* be the scalar value denoting the maximum value of 𝒖

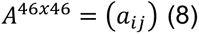

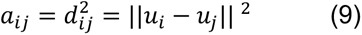

Where || · || denotes the Euclidian norm on ℝ^23^

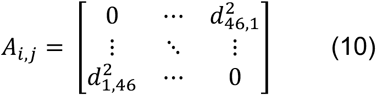

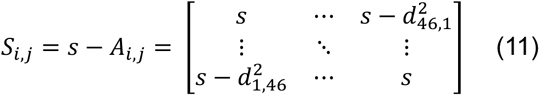

The resulting symmetric similarity matrix, 𝑆_*i,j*_, with rows and columns indicating each strain and each element representing the similarity between strain *i* and strain 𝑗 describes how strains are related to one another based on their projections along eigenvector 46. Hierarchical clustering on the resulting similarity matrix was then performed to identify groups of strains. Strains that are more similar are often found in communities that suppress *Kp-MH258* and those that are more distant are rarely found in communities that suppress *Kp-MH258*.

### Creation of ex-vivo media

8–10-week-old wild-type male C57BL/6J mice were used for all studies. Mice were initially obtained from The Jackson Laboratory and subsequently bred and raised in a GF isolator. After removal from the GF isolator, mice were handled in a sterile manner and individually housed in sealed negative pressure bio-containment unit isolators. Throughout breeding, mice were housed within the University of Chicago Gnotobiotic Research Animal Facility (GRAF) and maintained at a 12 hour light and 12 hour dark cycle and controlled humidity (30–70%) and temperature (68–79 °F). Gnotobiotic mice were fed an ad libitum diet of autoclaved Teklad Global 18% Protein Rodent Diet (Sterilizable) (2018S/2018SC). To create ‘GF cecal content media’, mice were euthanized and cecal contents were collected, weighted, and homogenized in 10mL of sterile distilled water on a of per gram of content. Cecal suspension was centrifuged, and supernatants were filtered through a 0.22 mm filter. GF cecal extract media was stored at -80°C.

To create ‘Ab-SPF cecal content media’ C57BL/6J SPF male mice at 8-10 weeks of age were singly housed and placed under an antibiotic regime (0.25g MNV – metronidazole, neomycin, vancomycin) in the drinking water (day 0). Four days later, antibiotic treatment was halted and mice were placed on normal acidified water (day 4). Cages and food were also changed. On day 7 were euthanized and cecal contents were collected, weighted, and homogenized in 10mL of sterile distilled water on a of per gram of content. Cecal suspension was centrifuged, and supernatants were filtered through a 0.22 mm filter. Ab-SPF cecal extract media was stored at -80°C.

### Determining engraftment of SynCom15 strains in SPF mice

To determine SynCom15 strain engraftment, 16s rRNA sequences from all 15 strains were blasted against 16S rRNA sequences derived from fecal samples of antibiotic-treated SPF mice gavaged with SynCom15 consortium. Fecal-derived sequences were assigned to a SynCom15 strain if their 16s rRNA percentage sequence identity was 100% with a minimum of a 95% coverage.

### Determining structure of microbiota in infected SPF mice given saline, FMT, or SynCom15

DNA was extracted using the QIAamp PowerFecal Pro DNA kit (Qiagen). Before extraction, samples were subjected to mechanical disruption using a bead beating method. Briefly, samples were suspended in a bead tube (Qiagen) along with lysis buffer and loaded on a bead mill homogenizer (Fisherbrand). Samples were then centrifuged, and supernatant was resuspended in a reagent that effectively removed inhibitors. DNA was then purified routinely using a spin column filter membrane and quantified using Qubit.

16S sequencing was performed for murine studies, where V4–V5 region within 16S rRNA gene was amplified using universal bacterial primers—563F (5′-nnnnnnnn-NNNNNNNNNNNN-AYTGGGYDTAAA-GNG-3′) and 926R (5′-nnnnnnnn-NNNNNNNNNNNN-CCGTCAATTYHT-TTRAGT-3′), where ‘N’ represents the barcodes and ‘n’ are additional nucleotides added to offset primer sequencing. Approximately 412-bp region amplicons were then purified using a spin column-based method (Minelute, Qiagen), quantified and pooled at equimolar concentrations. Illumina sequencing-compatible Unique Dual Index adapters were ligated onto the pools using the QIAseq 1-step amplicon library kit (Qiagen). Library quality control was performed using Qubit and TapeStation and sequenced on Illumina MiSeq platform to generate 2 × 250 bp reads.

Raw V4–V5 16S rRNA gene sequence data were demultiplexed and processed through the dada2 pipeline (v1.18.0) into amplicon sequence variants (ASVs) with minor modifications in R (v4.0.3)^92^. Specifically, reads were first trimmed at 190 bp for both forward and reverse reads to remove low-quality nucleotides. Chimeras were detected and removed using the default consensus method in the dada2 pipeline. Then, ASVs with length between 320 bp and 365 bp were kept and deemed as high-quality ASVs. Taxonomy of the resultant ASVs was assigned to the genus level using the RDP Classifier (v2.13) with a minimum bootstrap confidence score of 80^93^.

### Metabolic profiling of DMCs

For metabolite extraction from liquid cultures, samples were incubated at −80 °C between 1 h and 12 h. Four volumes of methanol spiked with internal standards were added to each culture supernatant. Samples were then centrifuged at −10 °C and 20,000 × g for 15 min followed by the transfer of 100 μL of supernatant to pre-labelled mass spectrometer autosampler vials (MicroLiter, 09-1200).

For metabolite extraction from fecal samples, extraction solvent (80% methanol spiked with internal standards and stored at -80 °C) was added at a ratio of 100 mg of material/mL of extraction solvent in beadruptor tubes (Fisherbrand; 15-340-154). Samples were homogenized at 4 °C on a Bead Mill 24 Homogenizer (Fisher; 15-340-163), set at 1.6 m/s with 6 thirty-second cycles, 5 seconds off per cycle. Samples were then centrifuged at -10 °C, 20,000 × g for 15 min and the supernatant was used for subsequent metabolomic analysis.

Short chain fatty acids were derivatized as described by Haak *et al*. with the following modifications^94^. The metabolite extract (100 μL) was added to 100 μL of 100 mM borate buffer (pH 10) (Thermo Fisher, 28341), 400 μL of 100 mM pentafluorobenzyl bromide (Millipore Sigma; 90257) in Acetonitrile (Fisher;A955-4), and 400 μL of n-hexane (Acros Organics; 160780010) in a capped mass spec autosampler vial (Microliter; 09-1200). Samples were heated in a thermomixer C (Eppendorf) to 65 °C for 1 hour while shaking at 1300 rpm. After cooling to RT, samples were centrifuged at 4 °C, 2000 × g for 5 min, allowing phase separation. The hexanes phase (100 μL) (top layer) was transferred to an autosampler vial containing a glass insert and the vial was sealed. Another 100 μL of the hexanes phase was diluted with 900 μL of nhexane in an autosampler vial. Concentrated and dilute samples were analyzed using a GC-MS (Agilent 7890A GC system, Agilent 5975C MS detector) operating in negative chemical ionization mode, using a HP-5MSUI column (30 m × 0.25 mm, 0.25 μm; Agilent Technologies 19091S-433UI), methane as the reagent gas (99.999% pure) and 1 μL split injection (1:10 split ratio). Oven ramp parameters: 1 min hold at 60 °C, 25 °C per min up to 300 °C with a 2.5 min hold at 300 °C. Inlet temperature was 280 °C and transfer line was 310 °C. A 10-point calibration curve was prepared with acetate (100 mM), propionate (25 mM), butyrate (12.5 mM), and succinate (50 mM), with 9 subsequent 2x serial dilutions.

Metabolites were also analyzed using GC-MS with electron impact ionization. The metabolite extract (100 μL) mass spec autosampler vials (Microliter; 09-1200) and dried down completely under nitrogen stream at 30 L/min (top) 1 L/min (bottom) at 30 °C (Biotage SPE Dry 96 Dual; 3579M). To dried samples, 50 μL of freshly prepared 20 mg/mL methoxyamine (Sigma; 226904) in pyridine (Sigma; 270970) was added and incubated in a thermomixer C (Eppendorf) for 90 min at 30 °C and 1400 rpm. After samples are cooled to room temperature, 80 μL of derivatizing reagent (BSTFA + 1% TMCS; Sigma; B-023) and 70 μL of ethyl acetate (Sigma; 439169) were added and samples were incubated in a thermomixer at 70 °C for 1 hour and 1400 rpm. Samples were cooled to RT and 400 μL of Ethyl Acetate was added to dilute samples. Turbid samples were transferred to microcentrifuge tubes and centrifuged at 4 °C, 20,000 × g for 15 min. Supernatants were then added to mass spec vials for GCMS analysis. Samples were analyzed using a GC-MS (Agilent 7890A GC system, Agilent 5975C MS detector) operating in electron impact ionization mode, using a HP-5MSUI column (30 m × 0.25 mm, 0.25 μm; Agilent Technologies 19091S- 433UI) and 1 μL injection. Oven ramp parameters: 1 min hold at 60 °C, 16 °C per min up to 300 °C with a 7 min hold at 300 °C. Inlet temperature was 280 °C and transfer line was 300 °C.

Data analysis was performed using MassHunter Quantitative Analysis software (version B.10, Agilent Technologies) and confirmed by comparison to authentic standards. Normalized peak areas were calculated by dividing raw peak areas of targeted analytes by averaged raw peak areas of internal standards.

### Evaluating suppressive capacity of SynCom15 on various strains of K. pneumoniae, E. coli, and C. difficile

The 15 bacterial strains that compose SynCom15 were individually inoculated from a frozen stock into 900 μL of BHIS previously reduced. Strains were incubated at 37°C in static conditions for 48h in anaerobiosis to ensure that the most fastidious strains reach stationary phase. Pathogen strains of *K. pneumoniae* (MH258-sGFP, MH225, MH189, MH1867, and Kp43816), *E. coli* (X43, T18002), VRE, and *C. difficile* (VPI, ST-49, R20291, ST-6, ST-75) were also inoculated in the same conditions, but only 24h after commensal isolates inoculation due to the fast growth capacity of these species and was incubated for 24h. All strain densities were assessed by taking 100 μL of each culture and measuring OD_600_ in a Biotek Cytation 5. To build SynCom15, SynCom15 minus commensal *E. coli* MSK19.28, Block 1, and Block 1 minus *E. coli* MSK19.28, isolates were inoculated in 900 μL of GF cecal content media previously reduced with an initial OD_600_ of 0.001, as described above. *K. pneumoniae, E. coli, VRE, or C. difficile* strains were added at the same initial OD_600_ of 0.001 to the different DMCs or to media alone. Cultures were incubated at 37°C in static conditions and anaerobiosis for 5 days. To *assess K*. *pneumoniae*, VRE, and *C. difficile* abundance, 10 μL of each culture were collected at 24h, 96, and 120h of culture and homogenized in 90 μL of PBS and serially diluted. Diluted samples were plated in: BHIS with kanamycin (50mg/mL) for the *Kp*-MH258-sfGFP strain; BHIS with ampicillin (100mg/mL) for *Kp*-MH225, *Kp*-MH189, *Kp*-MH1867, Kp43816, *E. coli* X43, and T18002; BHIS with Vancomycin (50mg/mL)/ Streptomycin (100mg/mL) for the VRE strain; and BHIS with norfloxacin [6g/L], moxalactam [16g/L], D-Cycloserine [62.5g/L], and Cefoxitin [16g/L] for all *C. difficile* strains. Plates were incubated at 37°C overnight in aerobiosis for all strains except for the *C. difficile* strains, which dilutions, plating and incubation of the plates were done under anaerobiosis. CFUs were enumerated 24h later for the *K. pneumoniae* and *E. coli* strains, and 48h later for VRE and *C. difficile* strains. In parallel, 100 μL of each culture was also collected to recover the cell phase and the supernatants at 24h, 96h, and 120h. These samples were stored at -80°C to be later processed for shotgun metagenomics and metabolomics.

### Elucidating indirect, pH dependent mechanism of suppression

For testing SynCom15 dependency on contact for pathogen suppression, the 15 bacterial strains that compose SynCom15 were individually inoculated from a frozen stock into 900 μL of BHIS previously reduced. Strains were incubated at 37°C in static conditions for 48h in anaerobiosis to ensure that the most fastidious strains reach stationary phase. Pathogen strains of *K. pneumoniae* (MH258-sGFP and MH225), *E. coli* (X43), and *C. difficile* (VPI and ST-49) were also inoculated in the same conditions, but only 24h after commensal isolates inoculation due to the fast growth capacity of these species and was incubated for 24h. Pathogen cultures were diluted into BHIS or GF cecal content media in 24-well plates, and DMCs cultures were diluted into upper chambers of Transwells inserts (0.4 *μ* m ThinCert for 24-well plates, greiner bio-one) such that each strain started at an OD_600_ of 0.001. To *assess K*. *pneumoniae*, *E. coli*, and *C. difficile* abundance, 10 μL of each culture were collected at 24h, 96h, and 120h of culture and homogenized in 90 μL of PBS and serially diluted. Samples were diluted, plated, incubated, and stored as described above.

DMCs plus pathogen or pathogen alone cultures were also collected to measure pH after 120h (pH probe Fisherbrand). When indicated culture’s pH was adjusted to 5.0 using HCl or 7.0 using NaOH. Cell-free spent medium from these cultures (with or without adjusted pH) were obtained using 0.22 *μ* m Millex-GP syringe filters (Millipore). Supernatants were used as culture media to inoculate the different pathogens. A supernatant derived from a culture that contains a specific pathogen was only used to inoculate the same pathogen. As before, 10 μL of each culture were collected at 24h, 96h, and 120h of culture and homogenized in 90 μL of PBS and serially diluted for plating the appropriated plates and conditions for the different pathogens, as indicated before. In parallel, 100 μL of each culture was also collected to recover the cell phase and the supernatants at 24h, 96h, and 120h. These samples were stored at -80°C to be later processed for shotgun metagenomics and metabolomics.

### Comparison of SynCom15 with microbiotas of healthy human donors

To investigate the presence of SynCom15 strains in samples from healthy human donors, SynCom15 strains taxonomic names were searched in the 22 fecal samples obtained from the DFI 22 human donors. For SynCom15 strain unclassified to species level *Bifidobacterium* sp., the most closely related species annotated by GTDB with an 98.21% ANI (*Bifidobacterium pseudocatenulatum*) was used^89,95^.

### Training an RF- and DNN-based model for predicting Kp-MH258 suppression from metabolite content

#### Data Handling

Our metabolite data consists of 118 metabolites measured as well as the Kp-MH258 CFUs at the end of 120 hours. The Kp-MH258 CFU measurement has a resolution of 1000 CFU. or any CFU value recorded as 0, we regularized it by adding a 1000. The log of this regularized CFU as well as the log fold changes are then unit norm-centered and used to train both Random Forest (RF) and Deep Neural Network (DNN) Models. Since norm-centering is an invertible transformation, we undo this transformation on the final model prediction value to get the predicted log suppression.

#### Training RF model

For training the RF model, we used a RandomForestRegressor, available with scikit-learn python package. Tree Depth was set to 12 levels per tree, the number of trees was set to 5000, and the maximum number of features was set to “sqrt” (square-root of the number of strains total). For any tree in a RF model, a randomly chosen 1/3^rd^ of the training datapoints is internally withheld as validation set. Since the RF model internally performs this cross validation using decisions from trees that don’t use a given datapoint, and the number of trees we have is large compared to the number of datapoints, the out-of-bag (OOB) accuracy score—the average accuracy of prediction on a datapoint across these trees—is our metric for generality of the model. For RF model for predicting Kp-MH258 suppressionbuilt on metabolites, the OOB score is **0.20** on a scale of 0 to 1 strongly suggesting that metabolites are a poor choice of basis. The score was stable to any additional trees added to the model. With the model trained, we noted the R^2 score on the training set. Using this RF model, we predicted and recorded the log suppression on the 60 true-out-of-sample experiments.

#### Training DNN based model

For training the metabolite based deep learning-based model DNN, we used the same input data format as described above. We next created 5000 Neural networks using PyTorch each with two hidden layers (50 neurons going to 10 neurons) and a single output neuron tasked with predicting the *Kp-MH258* log abundance values. For each neural network, we first performed a random 80% -20% training-validation splits. This neural network was trained on its training split using a dynamic momenta-dependent backpropagation algorithm where the gradient is calculated on the training split’s loss alone while the momentum parameter is reduced when the loss function on the validation split stagnates. This momentum-based scheme allows for smooth (non-oscillatory) and faster training and reduces the chance of our neural networks getting stuck in local optima. Once trained, we defined the generalization capacity of each neural network as the 𝑅^+^ value on the prediction of the validation split with floor set at 0. Higher 𝑅^+^ means that the neural network has properly generalized over the validation split. These 5000 trained neural networks were used to predict log suppression on the 60 true-out-of-sample experiments. We aggregated the result of this neural network ensemble experiment by weighing each prediction with the generalization capacity of the respective neural network to calculate the weighted mean and standard deviation of the predicted *Kp-MH258* log abundance values for all DMCs included (training/validation set as well as out-of-sample set).

## Supplementary Tables

**Supplementary Table 1**. Data pertaining to **Extended Data Figures 1** and **2**.

**Supplementary Table 2**. Phylogenetic description of 46 strains used to construct bacterial communities.

**Supplementary Table 3**. Data pertaining to design of DMCs.

**Supplementary Table 4**. Data pertaining to **Fig. 2B** and **2C**.

**Supplementary Table 5**. Validation data for ML models.

**Supplementary Table 6**. Data regarding validating ML models on out-of-sample DMCs.

**Supplementary Table 7**. Defining SynCom15 using machine learning and statistical inference of model constraints; data pertaining to **Fig. 4B** and **4D**.

**Supplementary Table 8**. Taxonomic compositional analysis of mouse experiments.

**Supplementary Table 9**. Metabolomic data associated with DMCs.

**Supplementary Table 10**. Mechanism of *Kp*-MH258 suppression by SynCom15.

**Supplementary Table 11**. SynCom15 suppression of phylogenetically distinct pathogens.

**Supplementary Table 12**. Fractional abundance of SynCom15 strain-associated species in fecal samples of healthy donors.

**Supplementary Table 13**. Data and analysis associated with **Extended Data Fig. 14**.

**Supplementary Table 14**. A summary of select previous microbiome design efforts.

## Supplementary Discussion

### On constructing consortia based on phylogenetic diversity using the UMAP-based embedding

As our goal was to design phylogenetically diverse consortia, we sought to do so in an algorithmic, quantitative manner. Typically, phylogeny is reflected as a hierarchy of relatedness between bacterial species^89,95–97^. While such a view provides a reasonable way to classify bacteria, hierarchies are intrinsically difficult to use as a parametrization of defining distance^32^. Conveyed using intuition: what does ‘different’ mean when objects are hierarchically related to each other? By virtue of being hierarchically related, two objects can be different but be relatively more similar than two other bacteria. As such, ensuring diverse sampling off a tree is a challenging concept to encode into an algorithm.

As an alternative to hierarchical parametrizations of ensembles, non-linear embeddings compress hierarchical relationships into an *N*-dimensional space where *N* is determined by the user^98–100^. In our case, we compressed the phylogeny of our strain bank into two UMAP-based dimensions, yielding a dimension-reduced space. Typically, such spaces are used purely for visualization. The reason for this is that distance in the space cannot be ascribed to hamming distance between objects due to the non-linear compression intrinsic to UMAP. What does this consideration practically mean for downstream analyses? First, new data cannot be projected into an existing UMAP space without a reparameterization of the space. Second, two UMAP plots comprised of distinct data cannot be compared to each other from a geometric standpoint. Third, computing distance within the UMAP space is meaningless with respect to the relative similarity between two objects.

However, in our case, the UMAP-based embedding directly captured phylogenetic differences between groups of strains. Note, this did not mean that distance in the UMAP scaled with phylogenetic distance, but that, by definition, large distances in the UMAP space corresponded to sampling of different phylogenies. As such, using dispersion in the UMAP space was a data-driven way to create diverse consortia. We note that this approach is likely imperfect and future work will be needed to create diverse ensembles directly off the inferred phylogenetic tree.

### Model Distillation

The motivation behind distillation in our work stems from a broader challenge in machine learning (ML): predictive models often succeed in fitting data but remain opaque with respect to the principles, if any, they have internalized. In fields such as computer vision and natural language processing, this has spurred the development of ‘model distillation’ frameworks which attempt to extract simplified rules, constraints, or low-dimensional representation from complex black-box models^49^. The underlying ethos is to bridge the gap between prediction and understanding. That is, move from models that merely output correct answers and towards models that reveal generative rules that can be reused across contexts.

We adapted this philosophy in our work to understand how the RF model was learning to predict the suppressive capacity of DMCs. A key difference in our approach from the typical approaches of distillation used in AI is that we did not focus on the architecture of the models themselves, but rather inferred constraints that the models learned within their latent representation of the data. The motivation behind this choice was inspired by previous studies that have interrogated the structure of biological systems using statistical inference spanning the fields of protein science, neuroscience, genome science, microbiome science, and cancer genomics^10,47,68,74,80,81,101–103^. The essence of the idea has been the following. Evolution creates diversity through the process of iterative selection and variation. However, the constraints governing this process are opaque and unintuitive from the perspective of a researcher and therefore we are not informed about a ‘forward model’ of biology. We therefore collect the outputs of this process—extant diversity—create an alignment to compare these outputs, and statistically learn constraints describing the outputs^104^. The result of this analysis provides an understanding of what component parts in complex biological systems that arise from the process of evolution are relatively important versus unimportant for the survival of the system (**Fig. SD1A**). In analogy to this approach, we derived a synthetic generating function in the form of a machine-learning model that can be used to engineer consortia with desired *Kp*-MH258 suppression. As such, we could mirror the process of creating extant diversity from the evolutionary process by creating lots of consortia that were predicted to achieve pathogen suppression by running the ML model. While the logic of suppression would not be revealed observing the composition of any single consortia, the collection of suppressive consortia would contain conserved taxonomic signatures reflecting that all consortia are predicted to suppress *Kp*-MH258. Thus, statistical decomposition of the consortia reflecting the ‘extant diversity’ of communities that suppress *Kp*-MH258 would reveal key taxonomic features learned by the synthetic generating function (**Fig. SD1B**).

**Figure SD1.**
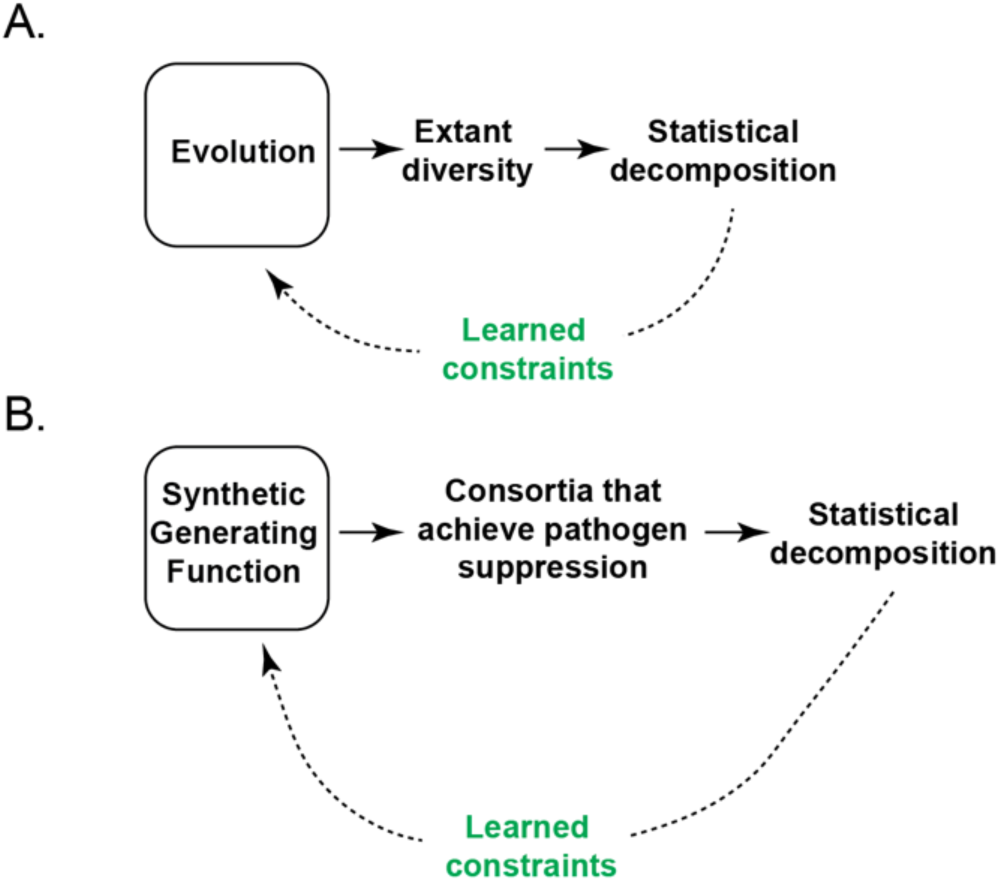
(A) Evolution creates extant diversity. The ensemble of extant diversity is subject to statistical decomposition to learn constraints about the evolutionary process. (B) In our work, a synthetic generating function was created in the form of an ML model. This ML model then created >5,000 consortia that were predicted to achieve pathogen suppression of greater than 10^5^ CFUs/ml. The resulting consortia are analogous to ‘extant diversity’—synthetic communities that survived the selection criteria of needing to achieve a high degree of pathogen suppression. The resulting consortia are subject to statistical decomposition to learn what constraints, if any, are latent within the statistical generating function.

### Assessing the compressive power of our approach

The process by which we converged on SynCom15 as a community that clears *K. pneumoniae* involved a six-step process: <u>C</u>ompress/<u>D</u>esign-<u>B</u>uild-<u>T</u>est-<u>L</u>earn-Infer. Within this process there were two points at which biological and experimental information was used for progressing through steps: (i) execution of the <u>C</u>ompress step utilized whole genomes of strains to reduce the size of the design space from spanning 848 possible strains to 46 possible strains and (ii) execution of the <u>D</u>esign-<u>B</u>uild-<u>T</u>est-<u>L</u>earn-Infer steps required screening 96 DMCs. We therefore sought to compute the equivalent of a ‘compression ratio’ for converging on a single complex community from a bank of 848 strains. In evaluating computational algorithms, the compression ratio is a measure of data complexity prior to compression relative to after compression. As our process considered biological information in the form of bacterial genome sequences and experiments, we normalized the compression ratio by the amount of information needed to perform the compression. We therefore defined an ‘effective’ compressive power as

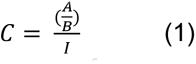

where C is the effective compressive power of a process, A is the complexity of data prior to compression, B is the complexity of data after compression, and I is the information needed for compression from A to B.

For our first step, we reduced the strain bank from 848 strains to 46 strains representative of the full phylogenetic diversity by genome sequencing each of the 848 strains, annotating each genome by their gene content, and performing dimension-reduction via a UMAP analysis. Therefore, the total complexity prior to compression was 2^848^/2, the total complexity after compression was 2^46^/2, and the information needed to be collected for compression were all base pairs of the 848 commensal strains (8.5 × 10^11^ basepairs). Considering these values, the effective compression of our first step was ∼10^230^—a substantial compression driven by the sizeable drop in complexity of the strain bank. For our second step, we used the diversity of the 46 strains to create 96 DMCs, 23 experiments to demonstrate the reproducibility of our assay, and 60 experiments to evaluate the ‘out-of-sample’ DMCs—a total of 179 experiments. Therefore, the total complexity prior to compression was 2^46^/2, the total complexity after compression was 1 (SynCom15), and the information needed to be collected for compression 179 experiments. Considering these values, the effective compression for our second step was ∼10^11^. Collectively, this analysis showed that despite the apparently immense amount of data reflected in the whole genome sequences of 848 bacterial strains, this complexity is offset by many orders of magnitude through Constraint Distillation.

### Contextualizing Constraint Distillation amongst prior microbiome design strategies

A broad range of strategies have been proposed for designing synthetic microbial communities with desired functions. As summarized in **Supplementary Table 14**, most efforts fall along distinct categories of design approaches. In one category are approaches that treat communities as reaction networks whose collective behavior can be captured by dynamical systems models. These studies typically use extensive information about individual strains—growth in mono- and co-culture, inferred or measured metabolic qualities, curated genome-scale metabolic models, or experimentally parametrized interaction matrices—to simulate community dynamics and identify candidate consortia for a target function. Such models can provide mechanistic insight and can enable *in silico* exploration of design space, but in practice require substantial prior knowledge, are difficult to parametrize at scale, and are often limited to specific conditions or considering small numbers of strains. Typically, newer approaches of statistical modeling—machine learning and artificial intelligence approaches like deep-learning—have been used within this category to augment Design-Build-Test-Learn paradigms. Although powerful, these methods are typically data hungry, requiring hundreds to thousands of tested communities, and are often optimized for prediction within a single experimental context. As a result, generalization of rules across environments, particularly regarding synthetic design of consortia towards gut microbiome-related applications, has remained largely untested using this strategy.

A second category is rooted in empirical reduction of natural communities. Here, complex microbiotas (e.g. whole fecal communities) are iteratively diluted, fractionated, or rationally trimmed to yield a defined consortium. This strategy implicitly treats the natural ecosystem as a blueprint: composition and structure of the starting community constrain which taxa are even considered, and down-sampling tends to preserve high-abundance or strongly interacting members. While this has yielded designed consortia, this strategy is predicated on the existence of natural microbiomes that have already solved the problem. Thus, design towards possibly novel target functions is not possible through using only this approach. Moreover, this strategy is not meant to create principles of design—the capacity to design a new, functional microbiome in a predictive manner.

A third category is leveraging existing sequencing efforts between human or mouse cohorts to perform comparative analysis and define bacteria that differentiate the cohorts from each other. A typical example would be to compare a ‘diseased’ cohort with a ‘healthy’ cohort then create a consortium based on these differences. This strategy has also been historically effective at creating consortia but requires (i) the presence of taxonomic differences between cohorts, (ii) the taxonomic differences faithfully map to differences in phenotype, and (iii) the method of comparative analysis is accurately capturing taxonomic differences from a statistical perspective without compressive loss of information. Like the second category, this strategy is also not meant to create principles of consortia design such that new consortia can be engineered to function in a predictive manner.

Constraint Distillation occupies a distinct point in this landscape (**Fig. SD2**). Our approach deliberately dispenses with several underlying foundations of prior work. We do not (i) model dynamical interactions between community members, (ii) encode mechanism-specific features related to *K. pneumoniae* suppression, (iii) rely on pathway-level annotations or curated metabolic networks at the scale of individual strains or designed communities, (iv) use human microbiome composition as a template, or (v) start from naturally occurring communities that already display the desired phenotype. Instead, we represent each designed community only by a binary vector of strain presence/absence, where the communities were initially designed solely based on genome-encoded diversity. From this sparse and functionally agnostic description along with a modestly sized screen of 96 consortia, we trained an ML model and then inferred functionally relevant statistical constraints learned by the model that defined SynCom15. Conceptually, our work complements rather than replaces prior efforts. Mechanistic, dynamical, and approaches embracing ‘-omics’ remain essential for elucidating how specific consortia function and for de-risking translation of live biotherapeutics into clinical settings. However, the SynCom15 case study demonstrates that generative design of a sparse, broadly effective community does not require these ingredients as prerequisites. Thus, Constraint Distillation offers a distinct mode of microbiome engineering: one that treats community design primarily as an inference problem on compositional space with mechanistic understanding as an output rather than an input of the design process.

**Figure SD2.**
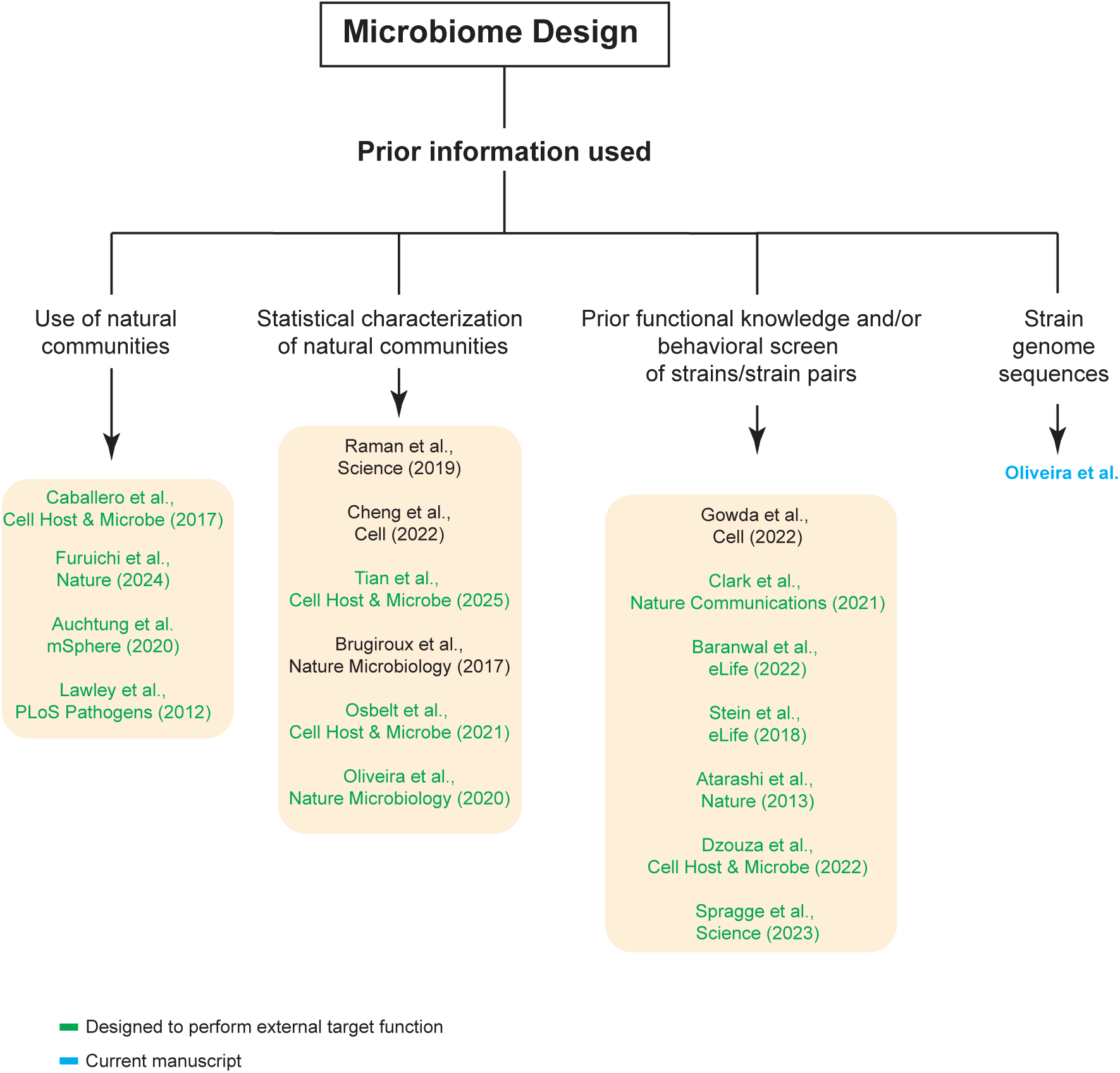
Previous efforts at community design can be conceptualized into three categories with respect to use of prior knowledge: (i) Use of natural communities to subsequently perform down-sampling, (ii) Statistical characterization of natural communities that reflect phenotypes of interests from cohorts, and (iii) Use of strain phenotypes/strain-strain interactions. Our manuscript illustrates the utility of using genome sequences of strains to <u>C</u>ompress the space of design and <u>D</u>esign metagenomically diverse consortia without any behavioral or functional knowledge of the constituent strains or communities. These communities are then <u>B</u>uilt, <u>T</u>ested for function, a model is <u>L</u>earned between strain presence/absence and output function, Inference is performed to distill taxonomic constraints learned by the model. Together these steps define the process we term ‘Constraint Distillation’.

## Data Availability

The datasets generated in our study are available within Supplementary Tables. Metagenomic data generated from profiling human fecal microbiomes used in this study are publicly available on NCBI under BioProject ID PRJNA838648. 16S data generated from mouse experiments used in this study will be publicly available on NCBI under BioProject ID PRJNA1074807. Raw data files associated with metabolomic data used in this study will be found on MassIVEW repository MSV000094183.

## Code availability

All code was written in either Python or R; code for all analysis will be found on Github (https://github.com/aramanlab/Oliveira_et_al_2024).

Figures. Figure panels associated with data were generated using either the Prism software (v10.2.0), various available packages in R, or Python. Figure schemes were generated using BioRender (BioRender.com) or Adobe Illustrator.

## Acknowledgements

We thank members of the Pamer, Kuehn, and Raman Laboratories for helpful discussion. We thank D. Pincus, M. Mani, A. Murugan, R. Ranganathan, and E. Pamer for helpful discussion. We thank members of the biobank, genomics, and metabolomics core services within the Duchossois Family Institute (DFI) at the University of Chicago and E. Pamer for their help in isolate collection, sequencing, and metabolomic profiling of all samples described in this manuscript. We thank the late E. Littman for his bioinformatic contributions. This work was supported by the Duchossois Family Institute (DFI) at the University of Chicago, the Dr. Ralph and Marian Falk Medical Research Trust, and NIH grant RM35GM146702.

## Author Information

E.M. aided in many of the experiments involving pathogen suppression and elucidating mechanism of SynCom15. B.P. aided in modeling of strain presence/absence with linear regression and deep-learning models as well as metabolomic-based modeling of consortia. K.L. wrote the code to implement the UMAP-based design strategy. M.Y. wrote the code to create the RF model based on strain presence/absence and scored 100,000 DMCs using the resulting RF model. R.Y.C. performed statistical analysis of the 100,000 DMCs to ultimately yield SynCom15; C.T. aided in the assay of all DMCs for evaluating their capacity to clear *Kp*-MH258 as well as the effect of selected communities in an *in vivo* setting. F.H. and V.A. aided with establishing the plate-based assay to evaluate clearing *Kp*-MH258; R.P. and K.S. aided in evaluating the suppressive capacity of SynCom15 against pathogen strains of *K. pneumoniae*, *E. coli*, and *C. difficile*. R.R. aided in analysis of taxonomic composition of fecal pellets procured from mice and analysis of fecal samples collected from healthy human donors. R.O. conducted all experiments involving DMCs across different conditions, all *in vivo* experiments, all experiments involving characterization of SynCom15, provided material for metabolomic and genomic analysis; S.K. and A.S.R. conceived of the statistical approach for community design involving creating a statistical model from an ensemble of diverse consortia; R.Y.C. and A.S.R. conceived of performing statistical inference on the learned model to distill model constraints. R.O. and A.S.R. conceived of the *in vitro* and *in vivo* experiments, experiments involved in understanding mechanism of SynCom15 action, the analysis of healthy human samples, metabolic profiling of DMCs, and sensitivity analysis of Constraint Distillation as a process. R.O. and A.S.R. wrote the manuscript.

## Ethics Declarations

Patents (63/543,XXX & 63/543XXX) related to this research have been filed by the University of Chicago with S.K., R.O., and A.S.R. as inventors.

## Materials and Correspondence

Author to whom correspondence and materials request should be addressed is A.S.R.

## References

1. Lawson, C. E. et al. Common principles and best practices for engineering microbiomes. Nat. Rev. Microbiol. 17, 725–741 (2019).

2. Goldford, J. E. et al. Emergent simplicity in microbial community assembly. Science 361, 469–474 (2018).

3. Faith, J. J., Ahern, P. P., Ridaura, V. K., Cheng, J. & Gordon, J. I. Identifying gut microbe-host phenotype relationships using combinatorial communities in gnotobiotic mice. Sci. Transl. Med. 6, 220ra11 (2014).

4. Inda, M. E., Broset, E., Lu, T. K. & de la Fuente-Nunez, C. Emerging frontiers in microbiome engineering. Trends Immunol. 40, 952–973 (2019).

5. Venturelli, O. S. et al. What is the key challenge in engineering microbiomes? Cell Syst. 14, 85–90 (2023).

6. Faust, K. & Raes, J. Microbial interactions: from networks to models. Nat. Rev. Microbiol. 10, 538–550 (2012).

7. Zomorrodi, A. R. & Segrè, D. Synthetic Ecology of Microbes: Mathematical Models and Applications. J. Mol. Biol. 428, 837–861 (2016).

8. Alivisatos, A. P. et al. MICROBIOME. A unified initiative to harness Earth’s microbiomes. Science 350, 507–508 (2015).

9. Dubilier, N., McFall-Ngai, M. & Zhao, L. Microbiology: Create a global microbiome effort. Nature Publishing Group UK 10.1038/526631a (2015) doi:10.1038/526631a.

10. Raman, A. S. et al. A sparse covarying unit that describes healthy and impaired human gut microbiota development. Science 365, (2019).

11. Cheng, A. G. et al. Design, construction, and in vivo augmentation of a complex gut microbiome. Cell 185, 3617–3636.e19 (2022).

12. Tian, S. et al. A designed synthetic microbiota provides insight to community function in Clostridioides difficile resistance. Cell Host Microbe 33, 373–387.e9 (2025).

13. Brugiroux, S. et al. Genome-guided design of a defined mouse microbiota that confers colonization resistance against Salmonella enterica serovar Typhimurium. Nat. Microbiol. 2, 16215 (2016).

14. Osbelt, L. et al. Klebsiella oxytoca causes colonization resistance against multidrug-resistant K. pneumoniae in the gut via cooperative carbohydrate competition. Cell Host Microbe 29, 1663–1679.e7 (2021).

15. Oliveira, R. A. et al. Klebsiella michiganensis transmission enhances resistance to Enterobacteriaceae gut invasion by nutrition competition. Nat Microbiol 5, 630–641 (2020).

16. Furuichi, M. et al. Commensal consortia decolonize Enterobacteriaceae via ecological control. Nature 633, 878–886 (2024).

17. Honda, K. et al. Rationally-defined microbial consortia suppress multidrug-resistant proinflammatory Enterobacteriaceae via ecological control. Res. Sq. (2023) doi:10.21203/rs.3.rs-3462622/v1.

18. Auchtung, J. M., Preisner, E. C., Collins, J., Lerma, A. I. & Britton, R. A. Identification of simplified microbial communities that inhibit Clostridioides difficile infection through dilution/extinction. mSphere 5, (2020).

19. Lawley, T. D. et al. Targeted restoration of the intestinal microbiota with a simple, defined bacteriotherapy resolves relapsing Clostridium difficile disease in mice. PLoS Pathog. 8, e1002995 (2012).

20. Petrof, E. O. et al. Stool substitute transplant therapy for the eradication of Clostridium difficile infection: “RePOOPulating” the gut. Microbiome 1, 3 (2013).

21. Gowda, K., Ping, D., Mani, M. & Kuehn, S. Genomic structure predicts metabolite dynamics in microbial communities. Cell 185, 530–546.e25 (2022).

22. Clark, R. L. et al. Design of synthetic human gut microbiome assembly and butyrate production. Nat. Commun. 12, 3254 (2021).

23. Baranwal, M. et al. Recurrent neural networks enable design of multifunctional synthetic human gut microbiome dynamics. Elife 11, (2022).

24. Stein, R. R. et al. Computer-guided design of optimal microbial consortia for immune system modulation. Elife 7, (2018).

25. Atarashi, K. et al. Treg induction by a rationally selected mixture of Clostridia strains from the human microbiota. Nature 500, 232–236 (2013).

26. Dsouza, M. et al. Colonization of the live biotherapeutic product VE303 and modulation of the microbiota and metabolites in healthy volunteers. Cell Host Microbe 30, 583–598.e8 (2022).

27. Spragge, F. et al. Microbiome diversity protects against pathogens by nutrient blocking. Science 382, eadj3502 (2023).

28. Morin, M. A., Morrison, A. J., Harms, M. J. & Dutton, R. J. Higher-order interactions shape microbial interactions as microbial community complexity increases. Sci. Rep. 12, 22640 (2022).

29. Gallardo-Navarro, O., Aguilar-Salinas, B., Rocha, J. & Olmedo-Álvarez, G. Higher-order interactions and emergent properties of microbial communities: The power of synthetic ecology. Heliyon 10, e33896 (2024).

30. Feng, L. et al. Identifying determinants of bacterial fitness in a model of human gut microbial succession. Proc. Natl. Acad. Sci. U. S. A. 117, 2622–2633 (2020).

31. 31. Front Matter. Inverse Problem Theory and Methods for Model Parameter Estimation i–xii Preprint at 10.1137/1.9780898717921.fm (2005).

32. Doran, B. A. et al. Subspecies phylogeny in the human gut revealed by co-evolutionary constraints across the bacterial kingdom. Cell Syst. (2025) doi:10.1016/j.cels.2024.12.008.

33. Skwara, A. et al. Statistically learning the functional landscape of microbial communities. Nat Ecol Evol 7, 1823–1833 (2023).

34. Lee, K. K., Park, Y. & Kuehn, S. Robustness of microbiome function. Curr. Opin. Syst. Biol. 36, 100479 (2023).

35. Gubbins, D. Book reviews. Geophys. J. Int. 94, 167–168 (1988).

36. Clermont, G. & Zenker, S. The inverse problem in mathematical biology. Math. Biosci. 260, 11–15 (2015).

37. Cocco, S., Feinauer, C., Figliuzzi, M., Monasson, R. & Weigt, M. Inverse statistical physics of protein sequences: a key issues review. Rep. Prog. Phys. 81, 032601 (2018).

38. Sanchez-Lengeling, B. & Aspuru-Guzik, A. Inverse molecular design using machine learning: Generative models for matter engineering. Science 361, 360–365 (2018).

39. Mora, T. & Bialek, W. Are biological systems poised at criticality? J. Stat. Phys. 144, 268–302 (2011).

40. Park, H., Li, Z. & Walsh, A. Has generative artificial intelligence solved inverse materials design? Matter 7, 2355–2367 (2024).

41. Odenwald, M. A. et al. Bifidobacteria metabolize lactulose to optimize gut metabolites and prevent systemic infection in patients with liver disease. Nat Microbiol 8, 2033–2049 (2023).

42. Xiong, H. et al. Distinct Contributions of Neutrophils and CCR2+ Monocytes to Pulmonary Clearance of Different Klebsiella pneumoniae Strains. Infect. Immun. 83, 3418–3427 (2015).

43. Hernández, U., Posadas-Vidales, L. & Espinosa-Soto, C. On the effects of the modularity of gene regulatory networks on phenotypic variability and its association with robustness. Biosystems. 212, 104586 (2022).

44. Hartwell, L. H., Hopfield, J. J., Leibler, S. & Murray, A. W. From molecular to modular cell biology. Nature 402, C47–52 (1999).

45. Newman, M. E. J. Modularity and community structure in networks. Proc. Natl. Acad. Sci. U. S. A. 103, 8577–8582 (2006).

46. Halabi, N., Rivoire, O., Leibler, S. & Ranganathan, R. Protein sectors: evolutionary units of three-dimensional structure. Cell 138, 774–786 (2009).

47. McLaughlin, R. N., Jr, Poelwijk, F. J., Raman, A., Gosal, W. S. & Ranganathan, R. The spatial architecture of protein function and adaptation. Nature 491, 138–142 (2012).

48. Espinosa-Soto, C. & Wagner, A. Specialization can drive the evolution of modularity. PLoS Comput. Biol. 6, e1000719 (2010).

49. 49. Frosst, N. & Hinton, G. Distilling a neural network into a soft decision tree. arXiv [cs.LG] (2017) doi:10.48550/arXiv.1711.09784.

50. Boix-Adsera, E. Towards a theory of model distillation. arXiv [cs.LG] (2024).

51. Arrieta, A. B. et al. Explainable Artificial Intelligence (XAI): Concepts, taxonomies, opportunities and challenges toward responsible AI. arXiv [cs.AI] (2019) doi:10.48550/arXiv.1910.10045.

52. Stutz, M. R. et al. Immunomodulatory fecal metabolites are associated with mortality in COVID-19 patients with respiratory failure. Nat. Commun. 13, 6615 (2022).

53. de Porto, A. P. et al. Fecal metabolite profiling identifies critically ill patients with increased 30-day mortality. Sci. Adv. 11, eadt1466 (2025).

54. Sorbara, M. T. et al. Inhibiting antibiotic-resistant Enterobacteriaceae by microbiota-mediated intracellular acidification. J. Exp. Med. 216, 84–98 (2019).

55. Bachman, M. A. et al. Genome-Wide Identification of Klebsiella pneumoniae Fitness Genes during Lung Infection. MBio 6, e00775 (2015).

56. Silver, R. J. et al. Amino acid biosynthetic pathways are required for Klebsiella pneumoniae growth in immunocompromised lungs and are druggable targets during infection. Antimicrob. Agents Chemother. 63, e02674–18 (2019).

57. Vornhagen, J. et al. The Klebsiella pneumoniae citrate synthase gene, gltA, influences site specific fitness during infection. PLoS Pathog. 15, e1008010 (2019).

58. Pickard, J. M., Zeng, M. Y., Caruso, R. & Núñez, G. Gut microbiota: Role in pathogen colonization, immune responses, and inflammatory disease. Immunol. Rev. 279, 70–89 (2017).

59. Oliphant, K. & Allen-Vercoe, E. Macronutrient metabolism by the human gut microbiome: major fermentation by-products and their impact on host health. Microbiome 7, 91 (2019).

60. Horrocks, V., King, O. G., Yip, A. Y. G., Marques, I. M. & McDonald, J. A. K. Role of the gut microbiota in nutrient competition and protection against intestinal pathogen colonization. Microbiology 169, 001377 (2023).

61. Baktash, A. et al. Mechanistic insights in the success of fecal Microbiota transplants for the treatment of Clostridium difficile infections. Front. Microbiol. 9, 1242 (2018).

62. Gehrig, J. L. et al. Effects of microbiota-directed foods in gnotobiotic animals and undernourished children. Science 365, (2019).

63. Delannoy-Bruno, O. et al. Evaluating microbiome-directed fibre snacks in gnotobiotic mice and humans. Nature 595, 91–95 (2021).

64. Kennedy, M. S. et al. Diet outperforms microbial transplant to drive microbiome recovery in mice. Nature 1–9 (2025).

65. 65. Plata, G., Srinivasan, K., Krishnamurthy, M., Herron, L. & Dixit, P. Designing host-associated microbiomes using the consumer/resource model. mSystems 10, e0106824 (2025).

66. Thompson, J. C., Zavala, V. M. & Venturelli, O. S. Integrating a tailored recurrent neural network with Bayesian experimental design to optimize microbial community functions. PLoS Comput. Biol. 19, e1011436 (2023).

67. Russ, W. P., Lowery, D. M., Mishra, P., Yaffe, M. B. & Ranganathan, R. Natural-like function in artificial WW domains. Nature 437, 579–583 (2005).

68. Socolich, M. et al. Evolutionary information for specifying a protein fold. Nature 437, 512–518 (2005).

69. 69. Bialek, W. & Ranganathan, R. Rediscovering the power of pairwise interactions. arXiv [q-bio.QM] (2007) doi:10.48550/arXiv.0712.4397.

70. Cavagna, A. et al. New statistical tools for analyzing the structure of animal groups. Math. Biosci. 214, 32–37 (2008).

71. Cavagna, A. et al. Scale-free correlations in starling flocks. Proc. Natl. Acad. Sci. U. S. A. 107, 11865–11870 (2010).

72. Caballero, S. et al. Cooperating Commensals Restore Colonization Resistance to Vancomycin-Resistant Enterococcus faecium. Cell Host Microbe 21, 592–602.e4 (2017).

73. Sulaiman, J. E. et al. Elucidating human gut microbiota interactions that robustly inhibit diverse Clostridioides difficile strains across different nutrient landscapes. Nat. Commun. 15, 7416 (2024).

74. Russ, W. P. et al. An evolution-based model for designing chorismate mutase enzymes. Science 369, 440–445 (2020).

75. Yeh, A. H.-W. et al. De novo design of luciferases using deep learning. Nature 614, 774–780 (2023).

76. Anishchenko, I. et al. De novo protein design by deep network hallucination. Nature 600, 547–552 (2021).

77. Watson, J. L. et al. De novo design of protein structure and function with RFdiffusion. Nature 620, 1089–1100 (2023).

78. Wang, J. et al. Scaffolding protein functional sites using deep learning. Science 377, 387–394 (2022).

79. Brixi, G. et al. Genome modeling and design across all domains of life with Evo 2. bioRxiv 2025.02.18.638918 (2025) doi:10.1101/2025.02.18.638918.

80. Zaydman, M. A. et al. Defining hierarchical protein interaction networks from spectral analysis of bacterial proteomes. Elife 11, (2022).

81. 81. Behera, V. et al. Conserved principles of spatial biology define tumor heterogeneity and response to immunotherapy. bioRxiv 2024.10.18.619136 (2024) doi:10.1101/2024.10.18.619136.

82. Li, W. & Godzik, A. Cd-hit: a fast program for clustering and comparing large sets of protein or nucleotide sequences. Bioinformatics 22, 1658–1659 (2006).

83. Huang, Y., Niu, B., Gao, Y., Fu, L. & Li, W. CD-HIT Suite: a web server for clustering and comparing biological sequences. Bioinformatics 26, 680–682 (2010).

84. Bolger, A. M., Lohse, M. & Usadel, B. Trimmomatic: a flexible trimmer for Illumina sequence data. Bioinformatics 30, 2114–2120 (2014).

85. Prjibelski, A., Antipov, D., Meleshko, D., Lapidus, A. & Korobeynikov, A. Using SPAdes De Novo Assembler. Curr. Protoc. Bioinformatics 70, e102 (2020).

86. Wood, D. E., Lu, J. & Langmead, B. Improved metagenomic analysis with Kraken 2. Genome Biol. 20, 257 (2019).

87. Camacho, C. et al. BLAST+: architecture and applications. BMC Bioinformatics 10, 421 (2009).

88. O’Leary, N. A. et al. Reference sequence (RefSeq) database at NCBI: current status, taxonomic expansion, and functional annotation. Nucleic Acids Res. 44, D733–45 (2016).

89. Chaumeil, P.-A., Mussig, A. J., Hugenholtz, P. & Parks, D. H. GTDB-Tk: a toolkit to classify genomes with the Genome Taxonomy Database. Bioinformatics 36, 1925–1927 (2019).

90. Seemann, T. Prokka: rapid prokaryotic genome annotation. Bioinformatics 30, 2068–2069 (2014).

91. Liaw, A. & Wiener, M. Classification and Regression by randomForest. (2007).

92. Callahan, B. J. et al. DADA2: High-resolution sample inference from Illumina amplicon data. Nat. Methods 13, 581–583 (2016).

93. Wang, Q., Garrity, G. M., Tiedje, J. M. & Cole, J. R. Naive Bayesian classifier for rapid assignment of rRNA sequences into the new bacterial taxonomy. Appl. Environ. Microbiol. 73, 5261–5267 (2007).

94. Haak, B. W. et al. Impact of gut colonization with butyrate-producing microbiota on respiratory viral infection following allo-HCT. Blood 131, 2978–2986 (2018).

95. Parks, D. H. et al. GTDB: an ongoing census of bacterial and archaeal diversity through a phylogenetically consistent, rank normalized and complete genome-based taxonomy. Nucleic Acids Res. 50, D785–D794 (2022).

96. 96. Guindon, S., Delsuc, F., Dufayard, J.-F. & Gascuel, O. Estimating Maximum Likelihood Phylogenies with PhyML. in Bioinformatics for DNA Sequence Analysis (ed. Posada, D.) 113–137 (Humana Press, Totowa, NJ, 2009).

97. Stamatakis, A. RAxML version 8: a tool for phylogenetic analysis and post-analysis of large phylogenies. Bioinformatics 30, 1312–1313 (2014).

98. McInnes, L., Healy, J. & Melville, J. UMAP: Uniform Manifold Approximation and Projection for Dimension Reduction. arXiv [stat.ML] (2018).

99. Kobak, D. & Berens, P. The art of using t-SNE for single-cell transcriptomics. Nat. Commun. 10, 5416 (2019).

100. Becht, E. et al. Dimensionality reduction for visualizing single-cell data using UMAP. Nat. Biotechnol. 37, 38–44 (2018).

101. Stringer, C., Pachitariu, M., Steinmetz, N., Carandini, M. & Harris, K. D. High-dimensional geometry of population responses in visual cortex. Nature 571, 361–365 (2019).

102. Eisen, J. A. Phylogenomics: improving functional predictions for uncharacterized genes by evolutionary analysis. Genome Res. 8, 163–167 (1998).

103. Pellegrini, M., Marcotte, E. M., Thompson, M. J., Eisenberg, D. & Yeates, T. O. Assigning protein functions by comparative genome analysis: protein phylogenetic profiles. Proc. Natl. Acad. Sci. U. S. A. 96, 4285–4288 (1999).

104. Stein-O’Brien, G. L. et al. Enter the Matrix: Factorization Uncovers Knowledge from Omics. Trends Genet. 34, 790–805 (2018).

